# ZBP1’s Inability to Convert Unmodified RNAs to the Z-form Underlies a Balanced Mechanism of RNA Recognition with ADAR1

**DOI:** 10.64898/2026.04.21.719927

**Authors:** Jeffrey B. Krall, Lily G. Beck, Parker J. Nichols, Quentin Vicens, Morkos A. Henen, Beat Vögeli

## Abstract

Z-DNA Binding Protein 1 (ZBP1) is a critical pattern recognition receptor within the innate immune response to viral infection. ZBP1 senses foreign nucleic acids in the unusual, left-handed Z-conformation via binding through its N-terminal Zα1 and Zα2 domains and activates downstream pro-pyroptotic, -apoptotic, and -necroptotic pathways to initiate cell death and allow for viral clearance. Both dsDNA and dsRNA can adopt the Z-conformation, however, the conformational change is energetically expensive, especially for dsRNA, and typically requires chemical modifications or protein binding to induce a right-to-left-handed conversion and stabilization. ZBP1 has been previously shown to bind and convert B-DNA to the Z-conformation and was assumed to be able to convert A-RNA as well, despite the lack of experimental validation. Here, we use a variety of Nuclear Magnetic Resonance (NMR) and other biophysical and biochemical experiments to characterize the Z-DNA and Z-RNA binding properties of ZBP1’s Zα1 and Zα2 domains. While ZBP1’s Zα domains are able to convert and stabilize unmodified dsDNA in the Z-conformation, both domains are incapable of flipping unmodified A-conformation dsRNA. We show that ZBP1’s Zα domains require dsRNAs with Z-promoting chemical modification in order for them to bind and stabilize the Z-conformation. These results contrast with the Zα domain from Adenosine Deaminase Acting on RNA 1 (ADAR1), which can bind and flip both dsDNA and dsRNA into the Z-conformation, potentially indicating finely tuned competition between ADAR1 and ZBP1 for pro-survival and pro-death outcomes, respectively. This work highlights the functional variability of Zα domains and narrows down the potential physiological substrates of ZBP1 in infection and disease.

**Graphical:** 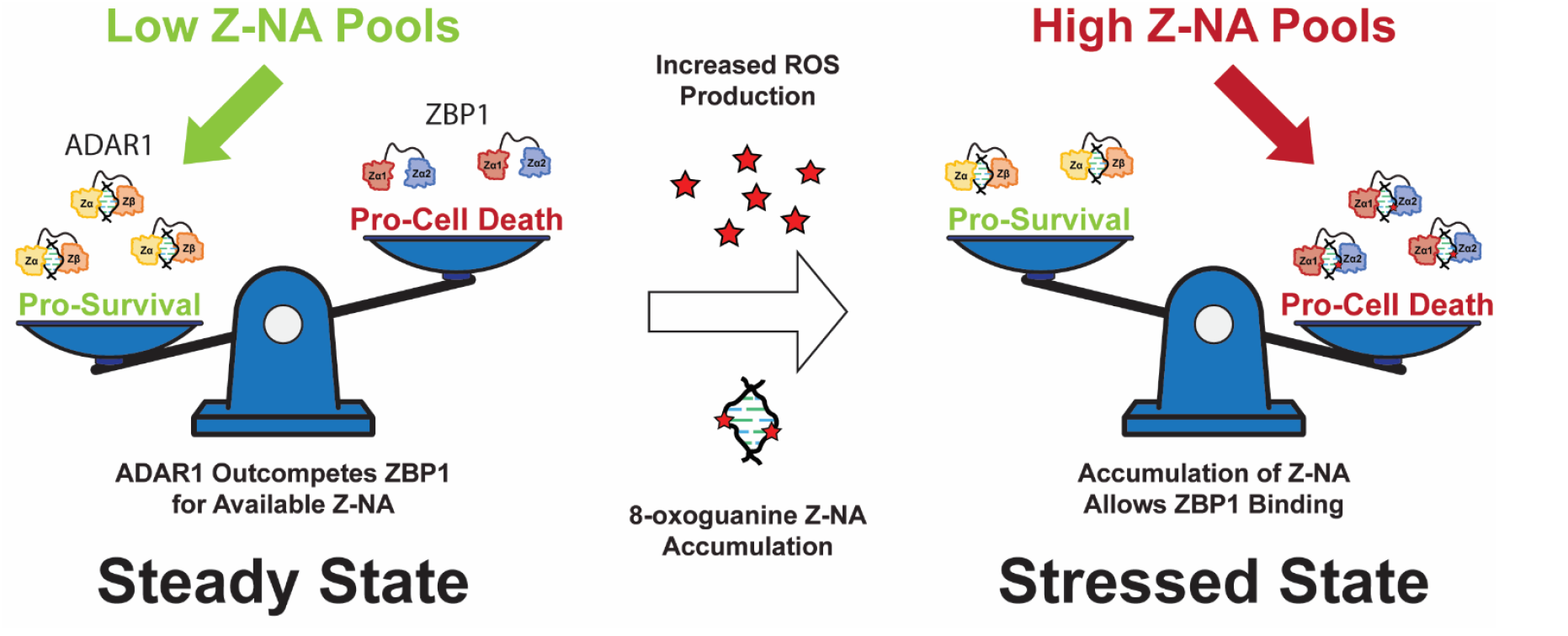

## INTRODUCTION

The presence of atypical nucleic acids or nucleic acid conformations is a key indicator of viral infection, and thus, the host cell utilizes a wide variety of nucleic acid pattern recognition receptors (PRRs) to recognize pathogens and mount an immune response [1, 2]. In addition to right-handed B-DNA and A-RNA, left-handed Z-conformation duplexes are generated during settings of viral infection, autoinflammation, and cancer [2], and can be sensed by specific PRRs as they are highly immunogenic. Whereas right-handed B-DNA and A-RNA adopt distinct duplex conformations, both Z-DNA/Z-RNA (collectively termed Z-NA) adopt a similar Z-conformation [3]. In the Z-conformation, the backbone forms a left-handed helix with glycosidic bonds alternating between *anti*- and *syn*-conformations and sugar puckers alternating between C2⍰-*endo* and C3⍰-*endo* [4, 5]. The unique features of the Z-form helix result in a “zig-zag” pattern of the phosphate backbone, from which it gets its name, and results in a closer approach of the two strands leading to electrostatic repulsions and contributes to the inherent instability of the Z-conformation. Z-RNA is generally more unstable than Z-DNA due to the unfavorable adoption of a C2⍰-*endo* sugar puckers. Consequently, the adoption of Z-RNA has an ~1.2 kcal/(mol•bp) higher activation energy when compared to its Z-DNA counterpart [6]. Although the Z-conformation is generally unfavorable under physiological conditions, certain factors, such as chemical modifications (extensively reviewed in [3]), protein binding [7–12], and negative superhelical stress [13, 14], can allow its stable adoption.

Z-Binding Domains (ZBDs) are the cognate binders of Z-NAs and can be found in a variety of innate immune response proteins, including Adenosine Deaminase Acting on RNA 1 (ADAR1; Figure 1A), Z-DNA Binding Protein 1 (ZBP1; Figure 1B), and in viral proteins such as E3L and ORF112 [15]. The function of these domains is necessary for the modulation of host immune responses and the virus’s ability to evade them. The nomenclature of ZBDs is not definitive; however, they can be divided into two main classes: the binding competent Zα domains and the structurally homologous, but functionally deficient Zβ domains. ZBP1 is the only known PRR for Z-NAs in mammals, and while it has low basal expression under homeostatic conditions, it is greatly upregulated during immunostimulation [2, 16]. The full-length gene product includes two N-terminal ZBDs (Zα1 and Zα2) responsible for Z-NA binding and three RIPK Homotypic Interaction Motifs (RHIMs) responsible for mediating the formation of the PANoptosome and downstream cell death signaling through concurrent activation of pyroptosis, apoptosis, and necroptosis (Figure 1B; [17–19]). Human ZBP1 is predominantly found in two main isoforms, ZBP1-Short and ZBP1-Long, in which the only difference stems from the excision of Exon 2 containing the first Zα1 domain [20]. While the isoforms perform a similar role sensing and responding to Z-NAs, their cellular localization and functionalities differ slightly [21, 22]. While both isoforms are predominantly cytoplasmic, ZBP1-Short is more prone to localizing to phase separates and forms large cytoplasmic granules in overexpression systems. ZBP1-Long, on the other hand, shows diffuse cytoplasmic distributions dependent on Z-binding ability, indicating that the Zα1 domain has a dominant effect on localization over Zα2 and that Zα2 may be responsible for ZBP1’s ability to undergo liquid-liquid phase separation [21].

**Figure 1.**
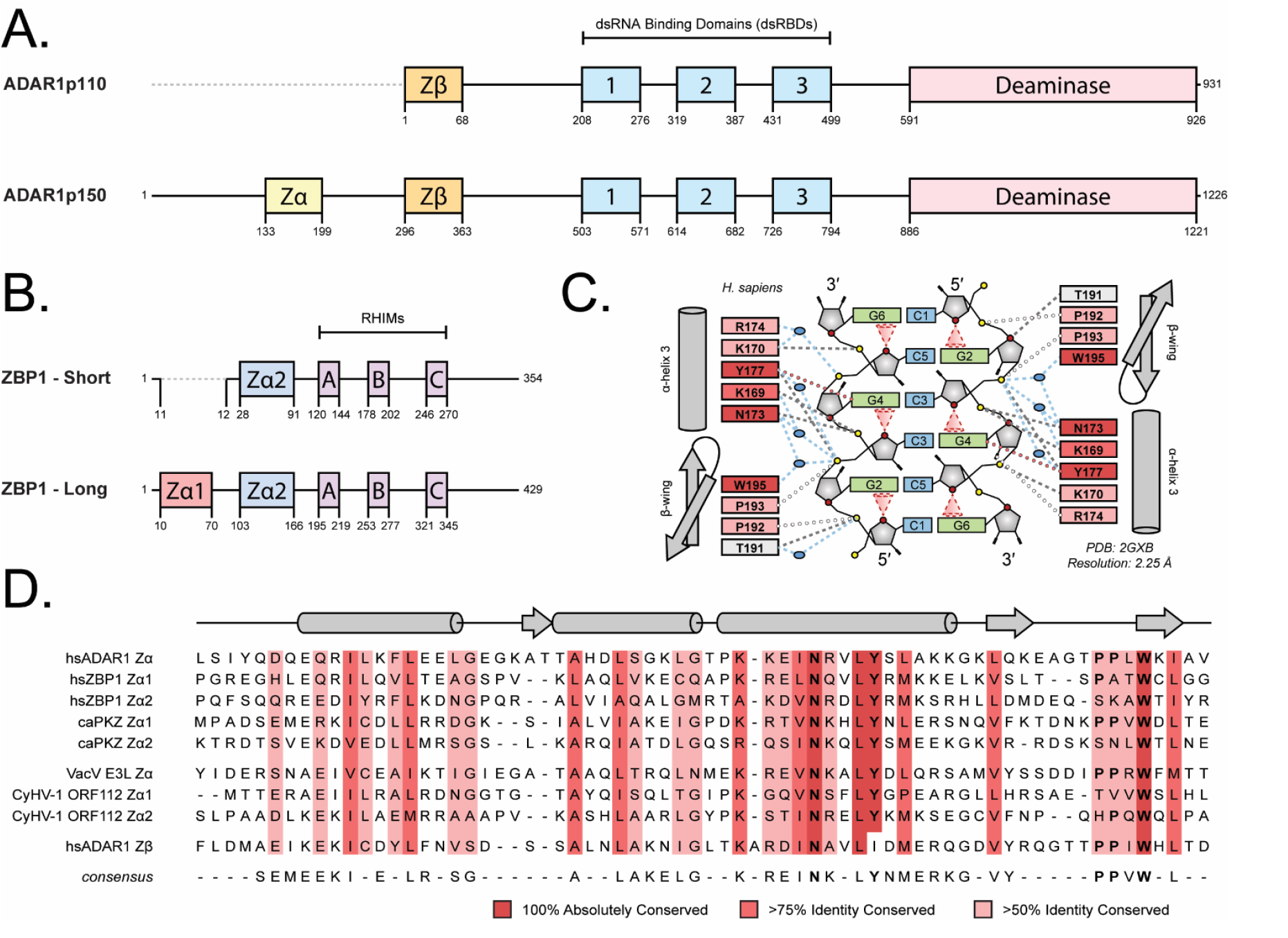
ZBDs in metazoan and viral Z-Binding Proteins (ZBPs). **(A)** Domain architecture of the two predominant isoforms of human Adenosine Deaminase Acting on RNA 1 (ADAR1). **(B)** Domain architecture of the two predominant isoforms of human Z-DNA Binding Protein 1 (ZBP1). RHIMs stand for RIP Kinase Homotypic Interact Motifs. **(C)** Schematic diagrams of the protein-Z-RNA interaction in Zα^ADAR1^ found in the crystal structure 2GXB generated by NUCPLOT [102]. Direct and water-mediated hydrogen bonds (<3.2 Å) are represented by grey and blue dashed lines, respectively. Lone pair···π contacts between ribose O4⍰ and guanine nucleobases are represented by red cones. Weak CH···π hydrogen bonds and close van der Waals interactions (<3.6 Å) are represented by red and white circles, respectively. **(D)** Multiple sequence alignments of the metazoan ZBDs from human ADAR1, human ZBP1, and goldfish PKZ and from the viruses, vaccinia virus and cyprinid herpesvirus, that infect humans and goldfish, respectively. Residues with high identity conservation are shaded in red according to the legend above.

ADAR1 was the first protein identified to harbor a ZBD and is considered the master negative regulator of the innate immune response to viral infection [23, 24]. Similar to ZBP1, ADAR1 has two predominant isoforms: ADAR1p110 and ADAR1p150 (Figure 1A) [25]. ADAR1p110 is constitutively expressed and contains an N-terminal Zβ domain, three tandem dsRNA Binding Domains (dsRBDs), and a C-terminal Adenosine Deaminase domain (Figure 1A). ADAR1p150 expression is driven by an alternative, interferon-inducible promoter and encodes an additional Zα domain N-terminal to the shared Zβ domain [25, 26]. Unlike ZBP1, both ADAR isoforms act to negatively regulate the innate immune response and prevent activation of cell death pathways. The deaminase domain catalyzes A-to-I editing in dsRNA substrates [27, 28], creating I:U wobble pairs thereby destabilizing dsRNA structures and preventing their recognition by the dsRNA PRRs, primarily the RLRs [1, 29, 30]. Additionally, the dsRBDs of ADAR1 are also known to directly compete with the dsRNA sensors, primarily PKR, for A-form nucleic acids to prevent their activation [1, 31]. ADAR1’s tandem ZBDs are thought to form a bipartite ZBD [32, 33], and similarly block ZBP1 activation through the competitive binding of Z-NAs [34, 35]. Because the p110 isoform is constitutively expressed, it is mostly responsible for preventing aberrant activation of cytosolic PRRs through the editing of nuclear dsRNAs before their export to the cytoplasm [36, 37]. The p150 isoform, on the other hand, is more important for the attenuation of ongoing IFN responses through editing and the sequestration of RNA substrates (extensively reviewed in [1]).

All Zα domains are comprised of three α-helices packed against three antiparallel β-sheets and use a similar interface to contact and stabilize the Z-conformation [10]. In addition to binding pre-formed Z-NAs, Zα domains have been shown to have A- and B-to-Z conversion activities [38–42], both of which require specific structural features. For Zα binding, there are several key amino acids critical for their interaction with Z-NAs. Using ADAR1’s Zα domain as reference (Figure 1C): Tyr177 allows for the recognition of nucleotides in the *syn* conformation by forming a CH···π hydrogen bond with the H8 proton of the central purine base. Additionally, Asn173 and Trp195 form both direct and water-mediated hydrogen bonds to the phosphate backbone of Z-NA and to themselves [7, 8, 15]. The Tyr, Asn, and Trp residues, being crucial to both the stability of the domain and to its interactions with Z-NAs, are 100% conserved in all Zα domains solved in complex with Z-DNA or Z-RNA (Figure 1D, Supplementary Figure 1) [15]. Mutation of any of these three residues abolishes binding and/or conversion activity [43, 44], and the lack of the conserved Tyr in Zβ explains its inability to bind to or stabilize the Z-form [45]. Auxiliary residues, including positively charged or amphipathic residues at positions 169, 170, and 174 within α-helix 3, display a lower identity conservation but still contribute to interactions with Z-NA.

Additionally, Pro192 and Pro193 lie at the apex of the β-wing, with Pro192 adopting the rare *cis* conformation[7]. This *cis*-*trans* proline-proline motif is important for the correct positioning of the β-wing along the Z-NA phosphate backbone and the overall function of the domain [7, 46]. The composition of residues within the β-wing immediately prior to the proline motif is also known to greatly affect the domain’s conversion activity, with the introduction of positively charged or hydrogen bonding-competent amino acids increasing both the rate of conversion and the conversion efficiency [47]. Recent studies on novel viral ZBDs show that while the lack of the proline motif does not affect B-to-Z conversion activity, A-to-Z conversion activity is absent [48]. Similarly, proline-to-alanine point mutation (P193A) within the proline-proline motif of ADAR1’s Zα domain abolishes A-to-Z conversion activity [46]. Consequently, the P193A mutation within ADAR1p150 is well known to cause severe autoimmune pathologies, including Bilateral Striatal Necrosis (BSN) [49] and Aicardi-Goutières Syndrome (AGS) when paired with a null allele [50]. Despite the functional consequences, these prolines only exhibit ~70% identity conservation [15]. Interestingly, both Zα1 and Zα2 of ZBP1 are lacking the double proline motif, with Zα1 retaining the first proline and Zα2 retaining neither. ZBP1’s Zα1 and Zα2 domains have been shown to induce or stabilize the Z-form of dsDNA in vitro [21], and are suspected to bind both Z-DNAs and Z-RNAs in vivo, as shown by the anti-Z antibody (Z22) [21, 35, 51–53].

Despite growing literature implicating ZBP1 in sensing both Z-DNA and Z-RNA across multiple disease settings, neither ZBP1’s ability to convert A-RNA to the Z-conformation nor its specific interactions with Z-RNA have been characterized structurally or biophysically. Instead, most studies expect ZBP1’s Zα1 and Zα2 domains to behave similarly to ADAR1’s Zα domain. Here, we use a variety of biophysical techniques, including Nuclear Magnetic Resonance (NMR), Circular Dichroism (CD), and Isothermal Titration Calorimetry (ITC) experiments to characterize the isolated and tandem ZBDs. We show that ZBP1 ZBDs can convert B-DNA to Z-DNA, albeit less efficiently than ADAR1 Zα, but cannot convert A-RNA, instead binding it in an A-form–like mode. ZBP1 constructs form unstable complexes with DNA and show limited ability to stabilize Z-conformations unless the substrate is predisposed. However, ZBP1 can bind and stabilize Z-RNA when modifications (e.g., m^8^G or o^8^G) lower the energetic barrier, unlike ADAR1 which actively induces Z-form NAs. This functional difference likely restricts ZBP1 sensing to pre-Z-forming nucleic acids, preventing aberrant immune activation and reflecting mechanistic differences from ADAR1.

## RESULTS

### Tandem ZBP1 Zα Domains do not Act Like a Bipartite ZBD

The expression of ZBDs in tandem can drastically change the functional properties of the protein [32, 33], and thus it becomes important to characterize the domains in isolation as well as in tandem in order to fully understand their properties. To characterize the behavior of the ZBP1 protein constructs (Supplementary figure 2A), we performed NMR relaxation experiments sensitive to different timescales to report on both global and local dynamics. Picosecond to nanosecond dynamics can be probed by both R1 and R2 relaxation rates, whereas microsecond to millisecond dynamics, which are often indicative of domain motion, are probed by R1ρ experiments or exchange contributions to R2 rates. Average ^15^N-R1 relaxation rates (Table 1; Supplementary Figure 3A-C) were remarkably similar for both Zα1^ZBP1^ (2.217 ± 0.003 s^−1^) and Zα2^ZBP1^ (2.089 ± 0.004 s^−1^). R1 relaxation rate averages for both Zα1 and Zα2 are lower in the Tandem^ZBP1^ construct (1.561 ± 0.004 s^−1^ and 1.563 ± 0.005 s^−1^, respectively), likely as a result of its rotational freedom being limited due to being in a tethered system.

**Table 1.** Measured relaxation rates^a^ and calculated rotational correlation times.

| Constructs | Domain | R1 [s<sup>-1</sup>] | R1ρ [s<sup>-1</sup>] | R2 [s<sup>-1</sup>] | R2/R1ρ | τ<sub>c</sub> [ns] |
| --- | --- | --- | --- | --- | --- | --- |
| Zα1<sup>ZBP1</sup> | Zα1 | 2.127 ± 0.003 | 6.216 ± 0.012 | 7.618 ± 0.052 | 1.261 ± 0.030 | 4.445 ± 0.068 |
| Zα2<sup>ZBP1</sup> | Zα2 | 2.089 ± 0.004 | 6.411 ± 0.015 | 6.766 ± 0.028 | 1.026 ± 0.025 | 4.663 ± 0.109 |
| Tandem<sup>ZBP1</sup> | Zα1 | 1.561 ± 0.004 | 11.094 ± 0.064 | 11.882 ± 0.103 | 1.100 ± 0.031 | 8.158 ± 0.229 |
|  | Linker | 1.530 ± 0.007 | 3.134 ± 0.011 | 4.033 ± 0.025 | 1.232 ± 0.031 | 3.075 ± 0.158 |
|  | Zα2 | 1.563 ± 0.005 | 9.948 ± 0.046 | 10.501 ± 0.047 | 1.051 ± 0.016 | 7.618 ± 0.016 |

<sup>a</sup>averaged over all residues of a domain

Similarly, average ^15^N-R2 relaxation rates (Table 1; Supplementary Figure 3A-C) were lower in the isolated Zα1^ZBP1^ (7.62 ± 0.05 s^−1^) and Zα2^ZBP1^ (6.77 ± 0.03 s^−1^) constructs than in the Tandem construct (11.88 ± 0.10 s^−1^ and 10.50 ± 0.05 s^−1^ for Zα1 and Zα2, respectively). ^15^N-R1ρ relaxation rate averages for Zα1^ZBP1^ (6.22 ± 0.01 s^−1^), Zα2^ZBP1^ (6.41 ± 0.02 s^−1^), and the two domains in the Tandem^ZBP1^ construct (11.09 ± 0.06 s^−1^ and 9.95 ± 0.05 s^−1^, respectively) largely mirrored the trends seen in the R2 rates (Supplementary Figure 3A-C). Taken broadly, the close agreement between the ZBP1 domain-average R2 and R1ρ rates indicate a general lack of dynamics in the μs to ms time regime covered by the R2 experiment and not quenched by the spin lock used in the R1ρ experiment (Supplementary Figure 3D-F). Interestingly, the isolated Zα2^ZBP1^ construct contains a region of consistently increased R2 over R1ρ rates corresponding to residues 135-142 within the N-terminus of its recognition helix, indicating the presence of μs-ms dynamics in this region (Supplementary Figure 3E). Additionally, residues Ala137 and Lys138 within this region show significant peak broadening in the ^15^N-HSQCs (Supplementary Figure 4), supporting the presence of μs-ms dynamics. Previous structural studies of Zα2 indicate that this region adopts a 3_10_-helical fold [10, 54], which is less stable than a typical α-helix and is likely to undergo transient folding/unfolding events [55]. The dynamic folding/unfolding of the 3_10_-helix within this region would explain both the increased R2 relaxation rates and the broadened peaks observed in the ^15^N-HSQCs.

Rotational correlation times (τ_C_; Table 1, Supplementary Figure 3G-I), which report on the molecular tumbling of molecules in solution, were calculated from R1 and R1ρ rates to minimize contributions from R2_Ex_, inflating τ_C_. Zα1^ZBP1^ and Zα2^ZBP1^ behave as small, well-folded domains with τ_C_ values of 4.45 ± 0.07 ns and 4.66 ± 0.11 ns, respectively, and fall within the expected range for proteins of their molecular weight (6.87 kDa and 7.11 kDa, respectively). τ_C_ values for Zα1 and Zα2 in the Tandem^ZBP1^ construct were 8.16 ± 0.23 ns and 7.62 ± 0.17 ns, respectively. The increase in tumbling time for both domains in the larger construct is likely a result of them existing in a higher molecular weight construct with reduced rotational freedom rather than as a result of dimerization. The calculated tumbling times of the ZBDs are far below those expected from a fully globular protein with a molecular weight of 20.66 kDa, let alone that of a dimer. These findings are in contrast to previous studies [56], which suggested that a Tandem ZBP1 construct forms dimers at concentrations similar to those in our experiments (~500 μM). Previous studies from our lab also suggest that the ZBDs of ZBP1 do not interact with one another and do not adopt distinct secondary structures when expressed in a tandem construct [57]. Taken together, the NMR relaxation rates confirm the structure and behavior of the isolated domains reported previously [9, 10, 54] and suggest that the Tandem^ZBP1^ construct does not act like a bipartite ZBD, unlike ADAR1’s tandem ZαZβ domain [32, 58].

### Circular Dichroism Reveals that ZBP1 is Incapable of Flipping Unmodified RNAs to the Z-Conformation

The functional variety between Zα domains is well documented [33, 40, 47, 48, 59], with some ZBDs being well characterized to be able to flip both dsDNA and dsRNA oligos to the Z-form [6], whereas others are only capable of flipping dsDNA [48], or only being able to stabilize pre-formed Z-conformations [47]. To test the ability of each ZBP1 construct to facilitate the right-to-left handed conversion and stabilization of the Z-conformation for both DNA and RNA oligos (Supplementary Figure 2B), we measured circular dichroism (CD) timecourses and spectra for a variety of oligos (Figure 2, Supplementary Figure 5). Due to the large difference in activation energies between the right-to-left-handed conversion of dsDNA and dsRNA [6], all DNA oligos were measured at 15°C, and all RNA oligos were measured at 42°C to roughly match their rates according to previous studies [6, 38]. For ease of comparison between the protein constructs measured within this study and to rates previously reported, all conversion activities were calculated using pseudo-first-order kinetics (Table 2). The ability of each construct to stabilize the Z-conformation can also be approximated from the total CD sign change. Here, the difference between the CD of the Z-conformation at equilibrium and that of the native right-handed A- and B-conformations can be calculated and used to compare Z-stabilization efficacy between domain constructs. Zα^ADAR1^ will be used as the point of comparison (100%), as it is the most well characterized ZBD and has been shown to convert both B-DNA and A-RNA oligos to the Z-form [5, 6, 38].

**Figure 2.**
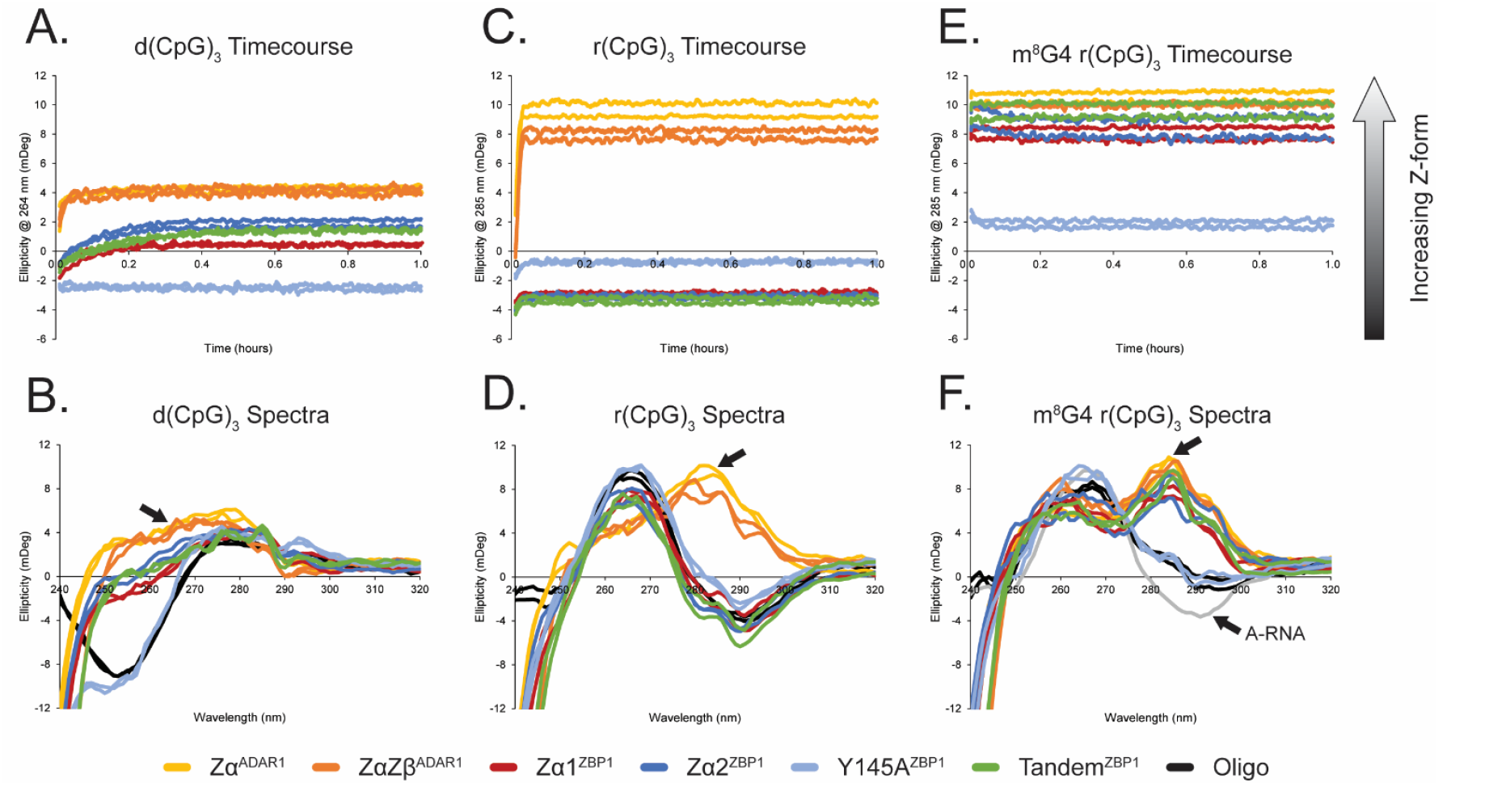
Circular Dichroism of ZBD constructs with small DNA and RNA oligos. **(A)** CD time course of ZBDs with d(CpG)_3_ oligos at 15°C. Formation and stabilization of Z-DNA is indicated by a growing positive ellipticity at 264 nm [black arrow in (B)]. **(C)** CD time course of ZBDs with r(CpG)_3_ oligos at 42°C. Formation and stabilization of Z-RNA is indicated by a growing positive ellipticity at 295 nm [back arrow in (D)]. **(E)** CD time course of ZBDs with m^8^G4 r(CpG)_3_ oligos at XXX°C. Formation and stabilization of Z-RNA is indicated by a growing positive ellipticity at 285 nm [back arrow in (F)]. **(B, D, F)** Full spectrum CD of (A-B) confirms the presence or absence of the Z-form after 1 hour. Black lines represent the native oligo conformations in the absence of ZBDs and the grey line in (F) represents a native A-conformation 6-mer RNA.

**Table 2.** A(B)-to-Z conversion rate constants (*k*) and conversion efficiencies extracted from circular.

| Oligo | Zα<sup>ADAR1</sup> |  | ZαZβ<sup>ADAR1</sup> |  | Zα1<sup>ZBP1</sup> |  | Zα2<sup>ZBP1</sup> |  | Y145A<sup>ZBP1</sup> |  | Tandem<sup>ZBP1</sup> |  |
| --- | --- | --- | --- | --- | --- | --- | --- | --- | --- | --- | --- | --- |
|  | *k* (hr<sup>-1</sup>) | % Z-form | *k* (hr<sup>-1</sup>) | % Z-form | *k* (hr<sup>-1</sup>) | % Z-form | *k* (hr<sup>-1</sup>) | % Z-form | *k* (hr<sup>-1</sup>) | % Z-form | *k* (hr<sup>-1</sup>) | % Z-form |
| d(CpG)<sub>3</sub><br>(15°C) | 38.15 ± 6.31 | 100.0 ± 0.4 | 52.60 ± 10.87 | 98.3 ± 0.5 | 12.04 ± 0.89 | 48.7 ± 0.3 | 9.28 ± 0.50 | 67.7 ± 0.4 | N/A |  | 6.21 ± 0.37 | 61.0 ± 0.4 |
| d(CpG)<sub>6</sub><br>(15°C) | 3.19 ± 0.07 | 100.0 ± 0.6 | 2.45 ± 0.07 | 96.3 ± 0.6 | 1.16 ± 0.03 | 86.4 ± 1.4 | LLPS<sup>‡</sup> |  | LLPS |  | LLPS<sup>‡</sup> |  |
| d(CpG)<sub>12</sub><br>(15°C) | 2.61 ± 0.06 | 100.0 ± 0.6 | 2.75 ± 0.09 | 111.0 ± 0.7 | 1.13 ± 0.03 | 81.6 ± 0.7 | LLPS<sup>‡</sup> |  | LLPS |  | LLPS<sup>‡</sup> |  |
| m<sup>8</sup>G4 r(CpG)<sub>3</sub><br>(15°C) | * | 100.0 ± 0.6 | * | 96.5 ± 0.6 | * | 81.3 ± 0.6 | * | 84.5 ± 0.6 | N/A |  | * | 92.9 ± 0.6 |
| r(CpG)<sub>3</sub><br>(42°C) | 107.00 ± 4.20 | 100.0 ± 0.3 | 96.82 ± 4.10 | 86.2 ± 0.3 | N/A |  | N/A |  | N/A |  | N/A |  |
| r(CpG)<sub>6</sub><br>(42°C) | 1.19 ± 0.01 | 100.0 ± 0.5 | 1.05 ± 0.02 | 92.5 ± 0.6 | N/A |  | N/A |  | N/A |  | N/A |  |
| r(CpG)<sub>12</sub><br>(42°C) | 3.00 ± 0.04 | 100.0 ± 0.5 | 2.70 ± 0.03 | 96.7 ± 0.4 | N/A |  | LLPS |  | LLPS |  | LLPS |  |
| LNA (CpG)<sub>12</sub><br>(42°C) | N/A |  | N/A |  | N/A |  | LLPS |  | LLPS |  | LLPS |  |

Both Zα^ADAR1^ and ZαZβ^ADAR1^ were able to quickly and efficiently convert the B-form dsDNA to the Z-conformation with rates of 38.15 ± 6.31 hr^−1^ and 52.6 ± 10.87 hr^−1^, respectively (Figure 2A). Zα^ADAR1^ and ZαZβ^ADAR1^ were able to effectively stabilize the Z-conformation to similar extents (100.0 ± 0.04% and 98.3 ± 0.5%, respectively; Figure 2B). When the ZBP1 constructs were incubated with a short d(CG)_3_ oligo, Zα1^ZBP1^, Zα2^ZBP1^, and Tandem^ZBP1^ were able to partially convert the DNA to the Z-form but did so at a much slower rate and with reduced conversion efficiency. Zα1^ZBP1^ was the fastest of the ZBP1 constructs with B-to-Z conversions rates of 12.04 ± 0.89 hr^−1^, followed by Zα2^ZBP1^ and Tandem^ZBP1^ with rates of 9.28 ± 0.50 hr^−1^ and 6.21 ± hr^−1^, respectively. Unintuitively, Tandem^ZBP1^, which has two functional ZBDs, was the slowest in converting the short d(CpG)_3_ oligo, suggesting the presence of either negative cooperation between the domains or another mechanism of self-regulation. Interestingly, Zα2^ZBP1^ was the most efficient ZBP1 construct in stabilizing the Z-form with 67.7 ± 0.4%, followed by Tandem^ZBP1^ and Zα1^ZBP1^ with 61.0 ± 0.4% and 48.7 ± 0.3% efficiency, respectively. As expected, the tyrosine-to-alanine mutant of Zα2 (Y145A^ZBP1^) showed no conversion activity or stabilization of the d(CpG)_3_ oligo in the Z-form as the ellipticity never crossed zero throughout the timecourse measurement and the CD spectra remained in the B-conformation. Overall, while ZBP1’s functional ZBD constructs were able to convert a short B-DNA oligo to the Z-form, they were much slower and less efficient when compared to ADAR1’s ZBD constructs.

We next tested each ZBD construct’s ability to convert an A-RNA oligo of the same length (Figure 2C, D). Similar to the 6-mer B-DNA, Zα^ADAR1^ and ZαZβ^ADAR1^ were able to quickly and efficiently convert an r(CpG)_3_ oligo to the Z-conformation with rates of 107.00 ± 4.20 hr^−1^ and 96.82 ± 4.10 hr^−1^, respectively. Recent studies have shown that A- and B-to-Z conversions proceed through a melted intermediate [38], and the higher temperatures at which the RNA oligos were measured (42°C versus 15°C) would lower the activation energy needed to proceed through this intermediate state, explaining their higher rates compared to the d(CpG)_3_. Despite the higher temperatures, Zα^ADAR1^ and ZαZβ^ADAR1^ were still able to efficiently stabilize the Z-conformation (100.0 ± 0.3% and 86.2 ± 0.3%, respectively). Unexpectedly, when we tested the ZBP1 constructs, none of them were able to convert the 6-mer RNA into the Z-conformation (Figure 2C). Although there was a small initial increase in the ellipticity, it never crossed zero, so rates were not fit. Additionally, full spectra measurements of the r(CpG)_3_ confirmed that the RNA remained in the A-conformation (Figure 2D). These results came as a surprise, as ZBP1 has historically been implicated in specifically sensing and responding to Z-RNA rather than its Z-DNA counterpart [60].

### Increasing Length and Flipping Cooperativity Does Not Promote ZBP1-Induced Z-RNA Conversion

Both A- and B-to-Z conversions are extremely cooperative after an initial nucleation event occurs and will proceed down the length of the oligo if the sequence allows it [38, 61, 62]. Therefore, we wondered whether the inability of ZBP1’s ZBDs to convert the r(CpG)_3_ RNA could be explained by the RNAs length. Subsequently, we also tested 12-mer and 24-mer DNA and RNA oligos to see if ZBP1 required cooperativity to flip RNA (Supplementary Figure 5). Doubling the length of a DNA (CpG) repeat to 12 bps drastically reduces the conversion rate induced by Zα^ADAR1^ (Supplementary Figure 5B), similar to previous reports [38]. Zα^ADAR1^ and ZαZβ^ADAR1^ were still able to effectively convert the d(CpG)_6_ oligo to the Z-form, but with rates an order of magnitude lower than the short d(CpG)_3_ oligo (3.19 ± 0.07 hr^−1^ and 2.45 ± 0.07 hr^−1^, respectively). Similar to the d(CpG)_3_ oligo, Zα^ADAR1^ and ZαZβ^ADAR1^ were nearly equally capable of stabilizing the Z-form with efficiencies of 100.0 ± 0.6% and 96.3 ± 0.6%, respectively. Doubling the length of the DNA again to create a d(CpG)_12_ 24-mer did not greatly affect Zα^ADAR1^ or ZαZβ^ADAR1^ rates (2.61 ± 0.06 hr^−1^ and 2.75 ± 0.09 hr^−1^, respectively) or efficiencies of stabilization (100 ± 0.6% and 111.0 ± 0.7%, respectively; Supplementary Figure 5C).

Similar to the ADAR1 ZBD constructs, increasing the length of the DNA oligos from a 6-mer to a 12-mer and 24-mer resulted in a significant decrease in Zα1^ZBP1^’s conversion rates (1.16 ± 0.03 hr^−1^ and 1.13 ± 0.03 hr^−1^, respectively). However, in contrast to the ADAR1 ZBD constructs, Zα1^ZBP1^’s efficiency in stabilizing the Z-form was greatly increased as length was increased from the d(CpG)_3_ oligo to the d(CpG)_6_ oligo, 48.7 ± 0.3% versus 86.4 ± 1.4%, indicating that ZBP1’s Zα1 domain relies on increased cooperativity to effectively stabilize the Z-conformation in DNA oligos (Supplementary Figure 5B). A further increase in length to the d(CpG)_12_ 24-mer did not result in a greater stabilization efficiency (81.6 ± 0.7%; Supplementary Figure 5C). Interestingly, when any ZBP1 ZBD construct containing Zα2 was incubated with the longer d(CpG)_6_ and d(CpG)_12_ DNA oligos, they precipitated out of solution and formed a clear gel-like pellet when centrifuged at 10,000rpm (Table 2). Unfortunately, the presence of phase separation hampers our ability to extract reliable rates for the Zα2-containing ZBD constructs as the particulates non-specifically block both right- and left-handed polarized light, resulting in a net zero ellipticity at the beginning of the measurement. Despite this limitation, Zα2^ZBP1^ and Tandem^ZBP1^ still convert the d(CpG)_6_ and d(CpG)_12_ oligos to the Z-form as evidenced by their full spectra CD profiles after the timecourse measurement, which show the characteristic peaks of the Z-DNA, albeit to a much lower extent (Supplementary Figure 5B, C).

While increasing the length of the DNA oligos had a subtle effect on each ZBP1 construct’s ability to convert and stabilize Z-DNA, increasing the length of the RNA oligos yielded no improvement on their A-to-Z conversion ability (Supplementary Figure 5E, F). Similar to incubation with a DNA 12-mer, incubation of Zα^ADAR1^ and ZαZβ^ADAR1^ with an r(CpG)_6_ oligo resulted in overall lower A-to-Z conversion rates of 1.19 ± 0.01 hr^−1^ and 1.05 ± 0.02 hr^−1^, respectively, when compared to the short 6-mer RNA oligos (Supplementary Figure 5E). With the 12-mer RNA (Supplementary Figure 5F), ZαZβ^ADAR1^ was slightly less effective in stabilizing the Z-conformation (92.5 ± 0.6%) when compared to Zα^ADAR1^ (100.0 ± 07%). Similar to previous reports [38], doubling the number of RNA bps again, from 12 to 24, results in an appreciable increase in the conversion rates. When Zα^ADAR1^ and ZαZβ^ADAR1^ were incubated with the r(CpG)_12_ oligo, both constructs conversion rates increased roughly 2.5x to 3.00 ± 0.04 hr^−1^ and 2.70 ± 0.03 hr^−1^, respectively. This increase in rate reflects the increase in cooperativity of the A-to-Z flip measured previously [38]. Unlike the ADAR1 ZBD constructs, when Zα1^ZBP1^, Zα2^ZBP1^, Y145A^ZBP1^, or Tandem^ZBP1^ were incubated with the r(CpG)_6_, no appreciable conversion activity was measurable (Supplementary Figure 5E). Similar to the short 6-mer RNA, there was an initial increase in ellipticity but no further increase or zero-crossing. Additionally, the full spectrum CD profiles confirmed that the 12-mer RNA remained in the A-conformation. Doubling the number of bps again to 24, thereby increasing the potential cooperativity, did not facilitate its conversion to the Z-form by any ZBP1 ZBD construct (Supplementary Figure 5F). Similar to the d(CpG)_12_ oligo, any Zα2-containing protein construct immediately phase separated with the r(CpG)_12_ oligo (Figure 3C), again confirming that Zα2’s propensity to induce LLPS is independent of its conversion activity.

**Figure 3.**
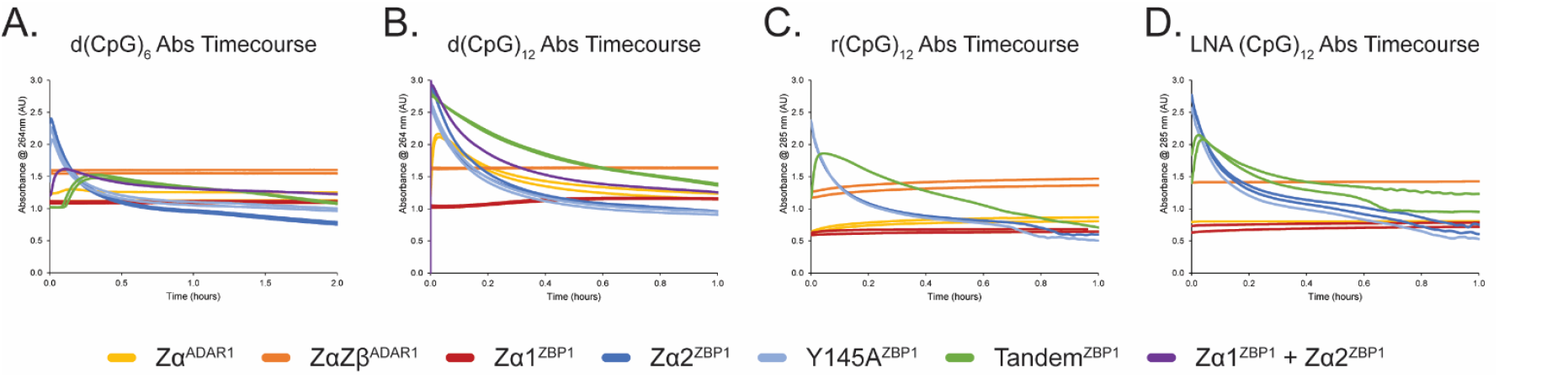
Absorbance time course measurements of oligonucleotides that display visible phase separation. Each color represents a different ZBD construct or combination and follow the legend below. **(A, B)** Absorbance time course measurements of the d(CpG)_6_ and d(CpG)_12_ oligos at 15°C, respectively. **(C, D)** Absorbance time course measurements of the r(CpG)_12_ and LNA (CpG)_12_ oligos at 42°C, respectively.

### *Cis-Trans* Proline-Proline Motif is Indispensable for A-to-Z Conversion Activity

Previous studies have shown that residues within the β-wings of ZBDs are critically important for A-to-Z conversion activity [46], and B-to-Z conversion efficiency [47, 63]. Of note, the *cis*-*trans* proline-proline motif at the apex of the β-wing is crucial for the proper folding and positioning the wing to contact the Z-conformation [7, 44, 46] (Supplementary Figure 6A-B). Neither Zα1^ZBP1^ nor Zα2^ZBP1^ contain the double proline motif and are incapable of converting (CpG)_n_ RNAs to the Z-conformation, but are still able to convert DNA, albeit with slower rates, as seen in this study and others [64]. Similarly, introduction of a proline-to-alanine mutation (P193A) within ADAR1’s Zα domain abolishes A-to-Z conversion activity [46], but does not greatly affect its affinity for Z-DNA [44]. However, to our knowledge, B-to-Z conversion rates have not been measured for the P193A mutant of ADAR1’s Zα domain (P193A^ADAR1^) using DNA oligos. Thus, we have also measured P193A^ADAR1^ with the d(CpG)_6_ and r(CpG)_6_ oligos (Supplementary Figure 6C-D) to gain a better understanding of the role of the proline-proline motif. When P193A^ADAR1^ was incubated with the d(CpG)_6_ oligo, its conversion rate and Z-stabilization efficiency were strikingly similar to Zα1^ZBP1^ (Supplementary Figure 6C). P193A^ADAR1^ had a B-to-Z conversion rate of 0.85 ± 0.02 hr^−1^ and a stabilization efficiency of 83.6 ± 1.2%, which represents a 3.8x decrease in conversion activity compared to the WT Zα^ADAR1^ construct and a 20% loss in Z-form stabilization. In contrast, P193A^ADAR1^ was only 1.2x slower than Zα1^ZBP1^ in converting the d(CpG)_6_ oligo and had a similar stabilization efficiency (83.6 ± 1.2 versus 86.4 ± 1.4%, respectively). Similar to previous reports [46], the P193A point mutation completely abolishes ADAR1’s A-to-Z conversion activity and resembles the results seen with ZBP1 ZBDs and the r(CpG)_6_ oligo (Supplementary Figure 6D). Therefore, the *cis*-*trans* proline-proline motif appears to be crucial for A-to-Z conversion activity but is largely dispensable for B-to-Z conversion. Lack of the motif reduces conversion activity but still allows for the stabilization of the Z-conformation, albeit to a lower extent. Zα^ADAR1^’s β-wing displays similar contacts between Z-DNA and Z-RNA in their respective structures (Supplementary Figure 1A & B), so it is unknown whether the lack of conversion activity in ZBP1 is caused by differential contacts between the A- and B-conformations and the proline motif for example, the lack of the proline motif and the resulting loss of favorable interactions may be sufficient to prevent the stabilization of the relatively higher energy Z-RNA conformation [6].

### ZBP1 Requires Chemically Modified RNA to Stabilize Z-RNA

Given ZBP1’s inability to convert unmodified r(CpG)_n_ repeats to the Z-conformation, we wondered if ZBP1’s ZBDs required chemical modifications to do so. Any chemical adduct at the C8-position of purine nucleobases heavily disfavors the *anti*-conformation, instead favoring the *syn*-conformation [3, 65, 66], thereby destabilizing the A- and B-conformations of RNA and DNA, respectively, and favoring the adoption of the Z-form [3]. Of note, 8-methylguanosine (m^8^G) modifications are known to greatly promote the adoption of the Z-conformation [67–69], with a single inclusion of an m^8^G at the central G4 position of a 6-mer RNA resulting in substantial Z-RNA adoption at physiological salt concentrations [38]. The m^8^G4 r(CpG)_3_ oligo has been previously reported to have distinct A- and Z-form populations and exists in a dynamic equilibrium dependent on temperature and salt concentration [38]. Under our experimental conditions at 15°C, we observe a similar phenomenon in which the m^8^G4 r(CpG)_3_ oligo displays characteristics of both A- and Z-form CD profiles (Figure 2F), indicating that it is also transiently sampling both A- and Z-conformations. Incubation of any of the ZBD constructs, with the exception of Y145A^ZBP1^, resulted in the immediate stabilization of the Z-conformation (Figure 2E). This conversion/stabilization occurred on a faster time scale before the cuvette could be placed into the spectrophotometer, and therefore, rates could not be measured (Table 2). With the m^8^G4 chemical modification, Zα1^ZBP1^, Zα2^ZBP1^, and Tandem^ZBP1^ were now nearly as efficient in stabilizing Z-RNA (81.3 ± 0.6%, 84.5 ± 0.6%, and 92.9 ± 0.6%, respectively) as the Zα^ADAR1^ and ZαZβ^ADAR1^ constructs were (100 ± 0.6% and 96.5 ± 0.6%, respectively). Taken together, our Circular Dichroism results suggest that ZBP1 is incapable of flipping unmodified RNAs into the Z-conformation, instead requiring chemical modifications to lower the high activation energy of melting the duplex or the Z-conversion.

### Zα2 Drives Phase Separation that is Independent of Z-Conversion Activity

ZBP1 has been well characterized for its ability to partition into liquid-liquid phase separates in overexpression systems and in physiological responses [21, 33, 56]. Our CD data showed that any ZBP1 construct containing Zα2 phase separated with the longer DNA and RNA oligos, whereas Zα1^ZBP1^ did not, indicating that Zα2 is the main driver of phase separation (Supplementary Figure 5B,C,F). This phenomenon was independent of Zα2’s Z-conversion ability, as the Y145A mutant, which is incapable of stabilizing the Z-conformation, phase separated similarly to the wild-type construct. Additionally, all Zα2-containing ZBP1 constructs, including Y145A^ZBP1^ similarly phase separated when incubated with an LNA (CpG)_12_ oligonucleotide, which is incapable of converting to the Z-form (Supplementary Figure 5H; [70, 71]), confirming that Zα2’s Z-conversion ability is not necessary for its phase separating abilities.

Although all Zα2-containing ZBD constructs phase separated with the longer oligos, each construct demonstrated different nucleation and settling rates apparent in the absorbance timecourse measurements (Figure 3). Absorbance measurements were taken simultaneously with the CD timecourse measurements and provided additional information on the interactions between the ZBD constructs and the oligonucleotides. The nucleation of LLPS droplets resulted in an initial increase in the absorbance measured as the light was scattered by the large droplets rather than being absorbed by the macromolecules in solution. The droplets, being denser than water, slowly settled out of solution over the course of the experiment, resulting in a slow decrease in light scattering and thus an apparent decrease in absorbance. When Zα2^ZBP1^ and Y145A^ZBP1^ were incubated with the d(CpG)_6_ oligo at 15°C, LLPS immediately formed and began to rapidly settle out of solution (Figure 3A). Interestingly, when the Tandem^ZBP1^ construct was mixed with the same oligo, there was an initial lag period of ~6 min before phase separation began, suggesting that the presence of Zα1 negatively affected the initiation of LLPS. Furthermore, nucleation of phase separation and the settling of the Tandem^ZBP1^ droplets occurred much more slowly than for the isolated Zα2 constructs, indicating that the smaller domain form larger or more dense phase separates than the tandem domains. To confirm that Zα1 had a negative effect on Zα2-driven phase separation, we also incubated equimolar amounts of Zα1^ZBP1^ and Zα2^ZBP1^ with the d(CpG)_6_ oligo. The presence of Zα1^ZBP1^ in *trans* slowed down the nucleation and settling of LLPS, however not to the same extent as Tandem^ZBP1^, confirming Zα1’s negative effect on LLPS that is enhanced when ZBP1’s ZBDs are expressed in tandem.

Doubling the length of the DNA oligo to 24 bps resulted in a drastic increase in the nucleation of droplets (Figure 3B). With the d(CpG)_12_ oligo, Zα2^ZBP1^, Y145A^ZBP1^, and Tandem^ZBP1^ had all nucleated phase separation immediately upon mixing and before the measurement could be started, suggesting that phase separation was partially dependent on oligo length. Despite similar fast nucleation times, the Zα2-containing constructs displayed similar trends in their settling rates, with Zα2^ZBP1^ and Y145A^ZBP1^ being nearly identical and having the fastest setting rates, followed by equimolar combinations of Zα1^ZBP1^ and Zα2^ZBP1^, and finally Tandem^ZBP1^. Interestingly, Zα^ADAR1^ also displayed an increase in absorbance that could be indicative of phase separation; however, no cloudiness or partitioning was observed at the end of the measurements, indicating that the increase in absorbance was due to a separate phenomenon or that Zα^ADAR1^ was able to resolve its LLPS by the end of the measurement period.

Interestingly, no phase separation was observed when any Zα2-containing protein construct was incubated with the r(CpG)_6_ oligo at 42°C as opposed to the d(CpG)_6_ oligo at 15°C. The lack of phase separation at higher temperatures could indicate that Zα2-driven phase condensates display Upper Critical Solution Temperature (UCST), as opposed to Lower Critical Solution Temperature (LCST), behavior. UCST phase transitions are enthalpically driven [72], in which specific interactions between charged, polar, and aromatic residues play a crucial role [73]. Doubling the length of the r(CpG)_6_ oligo to 24 bps results in a return of phase separation, confirming that phase separation is length dependent and indicates that the UCST can be increased with length as well. The phase separation of the r(CpG)_12_ oligo showed similar characteristics to the DNA 12- and 24-mers in which Zα2^ZBP1^ and Y145A^ZBP1^ showed nearly identical nucleation and settling rates and where Tandem^ZBP1^ had much slower nucleation and settling rates. The same trends were observed with the LNA (CpG)_12_, once again confirming that flipping ability is not necessary for the initiation of LLPS.

It is also important to note that all ZBDs displayed a minor gradual increase in absorbance over the course of the measurements (Supplementary Figure 5). These increases could be due to differences in base stacking interactions between the A/B- and Z-conformations that affect their respective electronic excitations, leading to differences in the UV-Vis absorbance profiles. This could be the case for the RNA oligos, as there is a moderate increase in the absorbance of the Z-RNA spectrum at 285 nm when compared to the A-RNA absorbance spectrum; however, for the DNA oligos, there is little to no change in the absorbance spectra at 264 nm [74]. Additionally, Zα1^ZBP1^, which does not phase separate nor flip RNA to the Z-conformation, similarly shows a moderate increase in absorbance over the course of the experiment. Alternatively, the increase in absorbance could be due to the generation and maintenance of a melted intermediate species. Both DNA and RNA experience hyperchromic effects when transitioning from ordered, native structures to disordered, denatured structures. Extinction coefficients increase across the visible spectrum for denatured oligos, however, maximal increases for GC-rich oligos occur around 280 nm for both DNA and RNA [75, 76]. As previously discussed, right-to-left-handed conversion of nucleic acids proceeds through melted intermediates [38], and thus, the increases in absorbance that we see for all oligos could be in part caused by the promotion or adoption of melted intermediates.

### B- and A-to-Z Conversions are Comprised of Distinct Processes Occurring Simultaneously

Although we used pseudo first-order kinetics to model each ZBDs conversion of right-handed oligos for simplicity, we found that our CD data was best explained by fitting multi-exponential rate equations (Supplementary Table 1). The total number of potential rates varied from one to three depending on the protein construct and the length of the nucleic acid but could generally be classified into fast (≥ 20 hr^−1^), intermediate (~10 hr^−1^), or slow (< 5 hr^−1^). The increased model complexity was justified using F-tests, as described in the methods section. The presence of multiple rates hints at different biological processes occurring simultaneously, although currently we can only hypothesize about what those processes are. In general, we assumed that the rate corresponding to the largest change in CD signal amplitude corresponds to the conversion rate as the right-to-left handed transition results in the largest difference in the CD spectra. A/B-to-Z conversion rates of the multi-exponential fits largely agreed with the single exponential rate of the pseudo-first order kinetics fits (Table 2, Supplementary Table 1). For the majority of oligos, the conversion rates fell within the slow processes, which agree with previous studies. Additionally, Zα1^ZBP1^, which lacks A-to-Z conversion activity, did not have a similar slow rate that would be indicative of flipping. For reasons similar to those encountered in the pseudo-first order kinetics analysis, many of the rates for Zα2^ZBP1^, Y145A^ZBP1^, and Tandem^ZBP1^ could not be fit due to the presence of LLPS interfering with measurements. The second and/or third rate constants accounted for much less of the overall amplitude of the data but greatly decreased the residuals of the fits.

The intermediate processes, when capable of being fit, accounted for the next largest contribution to the overall amplitude but were ~5x weaker on average. The rates for intermediate processes varied slightly across the measurable ZBD constructs; however, for each ZBD construct, there was a notable difference between DNA and RNA rates. On average, the intermediate rates for Zα^ADAR1^, ZαZβ^ADAR1^, and Zα1^ZBP1^ were 1.8x faster for DNA than for RNA (Supplementary Table 1), despite the RNA measurements being conducted at 42°C compared to the DNA measurements at 15°C. Z-RNA adoption has a much higher activation energy (3.2 kcal•mol^−1^•bp^−1^) compared to its DNA counterpart (2.0 kcal•mol^−1^•bp^−1^) [6], with much of this energy difference coming from the unfavorable adoption of the C2⍰-*endo* sugar pucker [3, 77]. Therefore, we think that this intermediate process is likely involved in the transient melting and/or rearrangement of the duplex into a Z-like state before full conversion to the Z-conformation. We also observed an extremely fast process that was dominant at the beginning of the measurements, but which accounted for the smallest percentage of the overall sign change in most cases outside of the short 6-mer oligos. When present, these rates were remarkably similar between all of the oligos for each ZBD, suggesting that the process governing it is inherent to the domain itself. This is corroborated by the fact that the rates are similar for DNA, RNA, and LNA regardless of each domain’s conversion capabilities. Given the timescales (~20-150 hr^−1^), these processes are unlikely to be involved in initial binding to the right-handed conformations but may still be involved in the nucleation of Z-forming regions.

Counterintuitively, the ADAR1 constructs, relative to the ZBP1 constructs, have extremely fast (~100 hr^−1^) processes that also account for the majority of the signal when incubated with the short, 6-mer RNA and DNA oligos and a second, slower process (~10 hr^−1^) that accounts for much less of the data. The ZBP1 constructs do not show the same trends with the 6-mer oligos; instead, they have a fast initial process with lower signal amplitudes, and a slower process that accounts for the majority of the signal, consistent with the longer oligos. It is currently unknown why the ADAR1 constructs display this characteristic, but it may be due to their ability to rapidly nucleate melted intermediates. The 6-mer DNA and RNA oligos have very low melting temperatures (T_M_’s) relative to the longer 12- and 24-mer oligos according to similar studies [38]. On the short oligos, Zα^ADAR1^ and ZαZβ^ADAR1^ may be able to fully melt the duplex and reform the strands into pseudo- or unstable Z-conformations. Under equilibrium conditions, these unstable complexes can slowly adopt more stable Z-conformations. Fitting multi-exponential rates, while much more complex, reveal consistent properties of each ZBD and their interactions with nucleic acids. However, the physical phenomenon underlying these rates is currently unknown, and future studies need to be conducted to more comprehensively understand the steps of Z-conversion.

### NMR Confirms Zα^ADAR1^’s Ability to Convert and Stabilize DNA and RNA Oligos in the Z-Form

To gain a better understanding of each ZBD’s interactions with right- and left-handed nucleic acids, we measured a series of 2D ^15^N-heteronuclear single-quantum coherence (HSQC) and 1D imino proton spectra of each ZBD with increasing concentrations of 6-mer oligos. Figure 4A-D shows the chemical shift perturbations (CSPs) induced in Zα^ADAR1^’s ^15^N-HSQC spectrum upon titration of stabilized Z-form m^8^G4 r(CpG)_3_, B-form d(CpG)_3_, A-form r(CpG)_3_, and A-like LNA (CpG)_3_ oligonucleotides, respectively, and in order of Z-conformation stability. Zα^ADAR1^ shows remarkably similar CSPs across all three convertible oligos, suggesting that the ZBD contacts the m^8^G4 r(CpG)_3_, d(CpG)_3_, and r(CpG)_3_ oligos in a similar manner. Of note, the critical Tyr177 residue in the recognition helix and Thr191, which sits at the apex of the β-wing, have large downfield chemical shifts to nearly the same positions for all of the Z-adopting oligos (Figure 4A-C). In contrast, Zα^ADAR1^ displays a distinct set of chemical shifts for the LNA 6-mer (Figure 4D), which is incapable of converting to the Z-conformation. Despite two distinct binding modes to the flippable and unflippable oligos, Zα^ADAR1^ uses nearly the same interface to contact both left- and right-handed oligos (Figure 5A). The most significant CSPs occurred primarily along the recognition helix and the β-wing, with similar large magnitudes and directionalities of chemical shifts for the m^8^G4 r(CpG)_3_, d(CpG)_3_, and r(CpG)_3_ oligos and much lower magnitudes and directionalities for the LNA (CpG)_3_ oligo.

**Figure 4.**
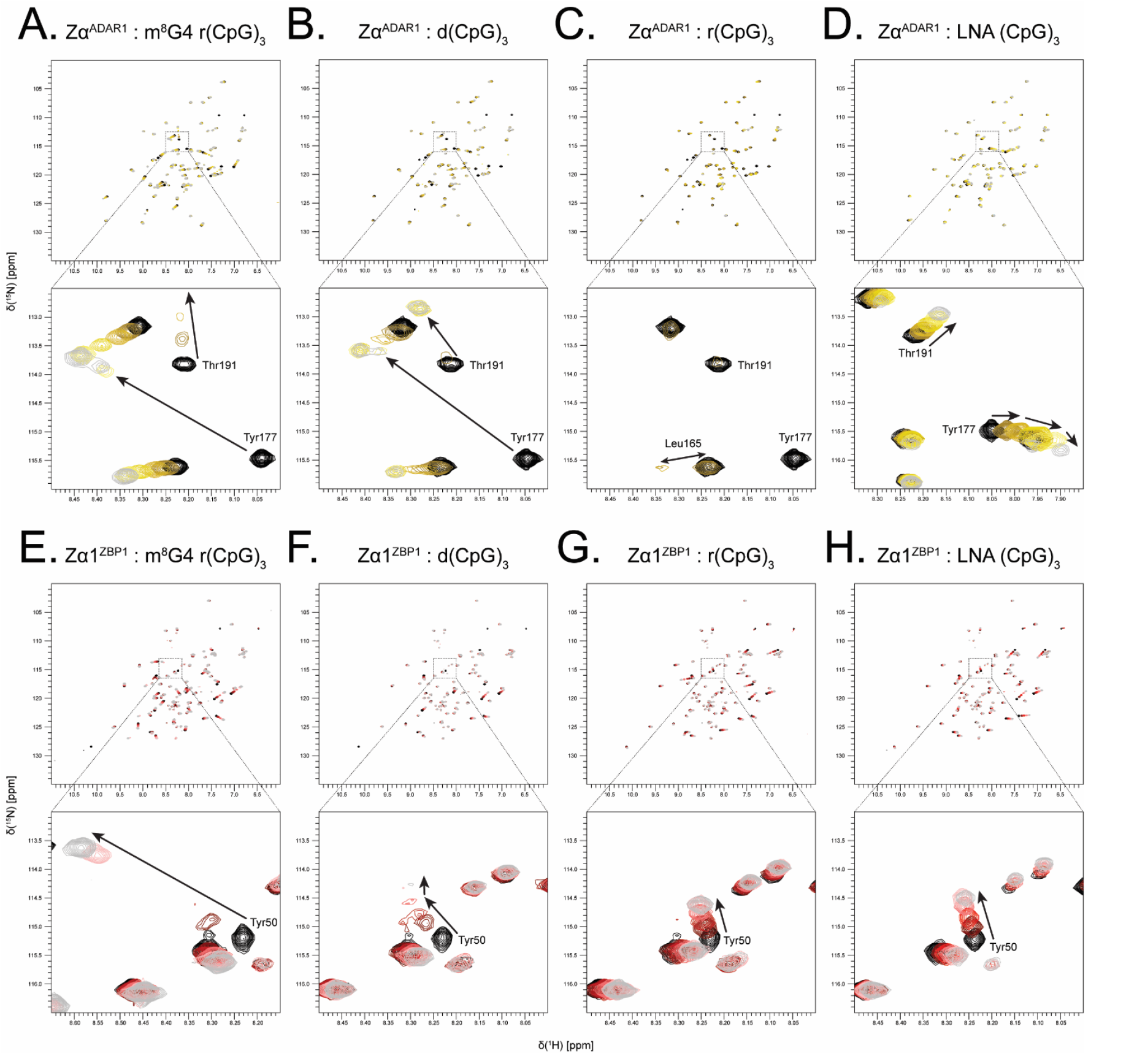
^15^N-HSQC titration series of the representative ZBDs of ADAR1 and ZBP1 with different 6-mer DNA/RNA oligos. **(A-D)** Titration series of Zα^ADAR1^ with the Z-form m^8^G4 r(CpG)_3_, B-form d(CpG)_3_, A-form r(CpG)_3_, and A-Like LNA (CpG)_3_, respectively. Titration points are color coded from black (Apo), through shades of yellow (1:0.25, 1:0.50, 1:1, 1:2), to grey (1:4). Inset boxes highlight residues Y177 and T191 which show significant peak shifts and are critical for flipping abilities. Y177 and T191 in the r(CpG)_3_ series shift to similar positions as seen with the m^8^G4 r(CpG)_3_ and d(CpG)_3_ residues but are heavily broadened. **(E-H)** Titration series of Zα1^ZBP1^ with the same oligos as shown in (A). Titration points are color coded from black (Apo), through shades of red (1:0.25, 1:0.50, 1:1, 1:2), to grey (1:4). Inset boxes highlight the CSPs of the critical Y50 residue.

**Figure 5.**
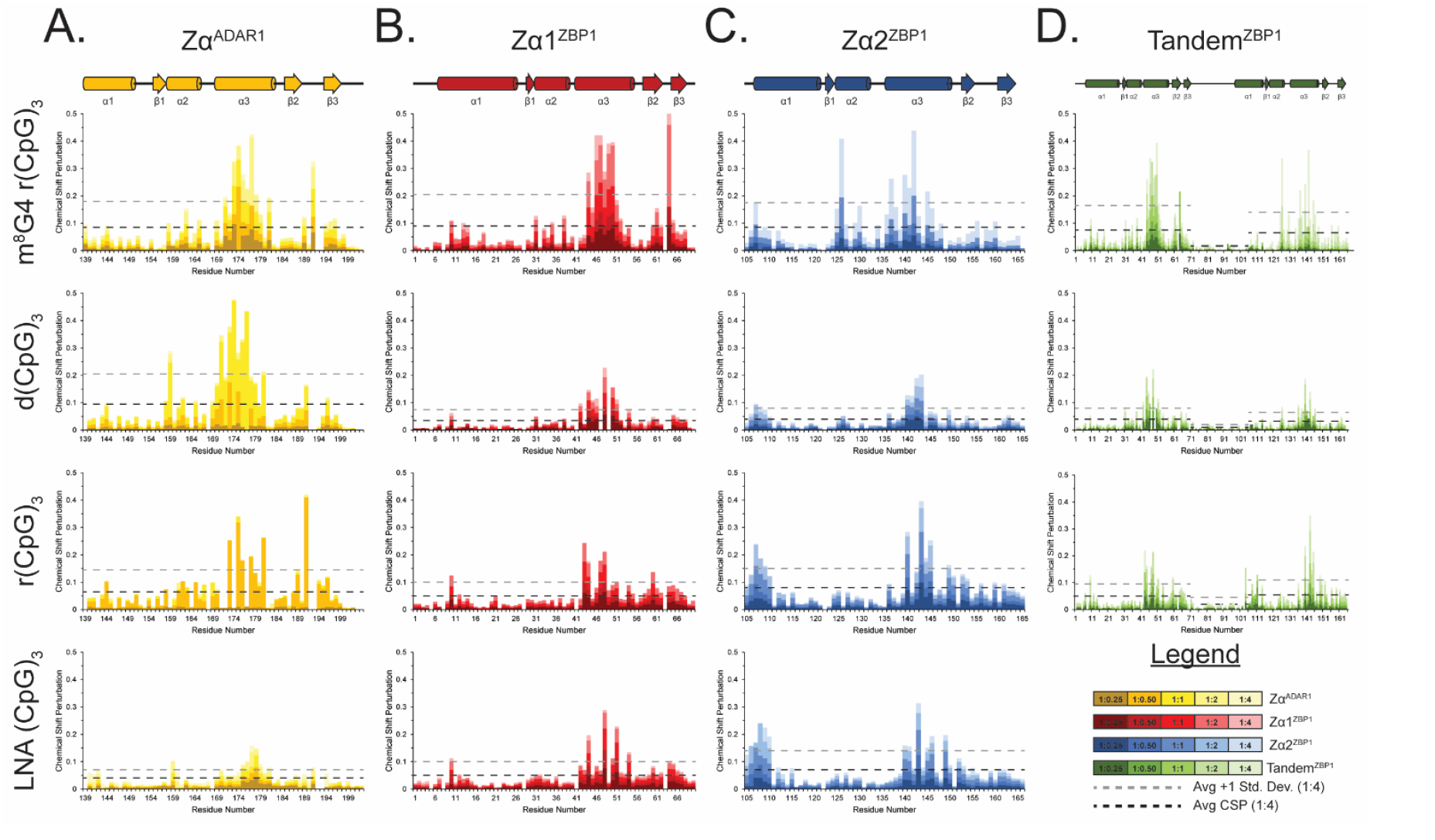
Waterfall plots of CSPs calculated from the ZBD construct ^15^N-HSQC titration series with the different 6-mer DNA/RNA oligos. CSPs were calculated using **Equation (2)** and represent the total shift from each titration point to the Apo protein ^15^N-HSQCs and are shaded according to the legend. Average CSPs (black dashes) were calculated for each domain and significantly shifted peaks were determined to be peaks one standard deviation above the average (grey dashes) or more. **(A)**. CSPs calculated from the Zα^ADAR1^ titration series with the different oligos. CSPs for the r(CpG)_3_ series show all or nothing behavior typical of peaks undergoing slow exchange. **(B)**. CSPs calculated from the Zα1^ZBP1^ titration series with the different oligos **(C)**. CSPs calculated from the Zα2^ZBP1^ titration series with the different oligos **(D)** CSPs calculated from the Tandem^ZBP1^ titration series with the different oligos.

Despite largely similar CSPs for the Z-adopting oligos, there were minor differences in the binding modes of the DNA and both RNA oligos, but the patterns closely matched those documented in previous studies [78]. Similarly, the Zα^ADAR1^ – LNA (CpG)_3_ titration series yielded chemical shifts identical to previous studies [71], including multidirectional chemical shift changes at high nucleic acid concentrations characteristic of multimodal binding modes. Interestingly, each titration series displayed binding behaviors occurring on different timescales. Most amide signals in the m^8^G4 r(CpG)_3_ series retained comparable strong intensities throughout titrations, indicating that binding to the modified RNA is occurring on the fast time scale. However, residues in the interaction interface along α3 and the β-wing experienced intermediate exchange characterized by the fading of signal intensity at intermediate titration points with linear peak shifts (Figure 4A, Supplementary Figure 7A). Largely, exchange regimes are slowed down in the Zα^ADAR1^ – d(CpG)_3_ titration series, with most amide signals experiencing fast-intermediate exchange regimes. Residues in the interaction interface are likewise slower, experiencing intermediate exchange (Figure 4B, Supplementary Figure 7B). The Zα^ADAR1^ – r(CpG)_3_ titration series, on the other hand, shows characteristics of both intermediate and slow exchange. Residues outside the interaction interface exhibit mostly slow exchange, characterized by the gradual disappearance of a peak at one location and its gradual appearance at a new position (Figure 4C, Supplementary Figure 8A). Amide signals in the interaction interface also experience slow exchange processes but also experience heavy peak broadening due to additional contributions from intermediate exchange or increased macromolecular size (Figure 4C). Interestingly, only fast exchange processes were observed in the Zα^ADAR1^ – LNA (CpG)_3_ titration series, including residues within the interaction interface (Figure 4D, Supplementary Figure 7D). Additionally, residues in the LNA series showed many nonlinear chemical shifts (Supplementary Figure 8B), consistent with previous reports of multiple LNAs binding the same Zα^ADAR1^ domain [71].

Lee et al. [78] observed similar trends in their titrations of Zα^ADAR1^ into d(CpG)_3_ and r(CpG)_3_ oligos under similar experimental conditions. They proposed that the slower exchange processes seen in the RNA series were due to its much slower conversion rate compared to DNA. Our results closely match their observations and extend their findings. The conversion rate of the m^8^G4 r(CpG)_3_ oligo is extremely fast by CD, and similarly, the majority of the amide signals experience only fast exchange. For all three convertible oligos, residues within the interaction interface experienced additional contributions from the intermediate exchange regime, possibly reflecting the on/off rates of the Z-NA – protein complex (Supplementary Figure 7A-C). Likewise, the nonconvertible LNA titration series only experienced fast exchange, which corroborates a lack of conversion activity, and therefore, a lack of additional exchange processes (Supplementary Figure 7D). The fast exchange seen likely reflects the affinity of Zα^ADAR1^ for the A-type helix, which is an order of magnitude lower than the domain’s affinity for convertible oligos [71]. Alternatively, the decrease in amide signal intensities at intermediate titration points could be due to increased complex size. Under protein saturating conditions, two ZBDs would be able to bind each 6-mer duplex, increasing molecular size and decreasing signal intensity. As nucleic acid concentrations increase to the 1:1 titration point, one ZBD would be titrated off, decreasing complex size and resulting in a return of signal. However, this is unlikely as these decreases in signal intensity should be consistent across the entire domain, especially for well-folded proteins. Overall, the ^15^N-HSQC titrations of Zα^ADAR1^ confirmed its ability to convert both DNA and RNA to the Z-form and inability to convert LNAs. Additionally, the different exchange regimes seen in each titration series reflect Zα^ADAR1^’s conversion rates for each oligo or lack thereof.

### NMR Reveals that ZBP1 ZBDs Form Unstable Complexes with DNA and are Incapable of Converting RNA

Unlike ADAR1’s Zα domain, which contacts all convertible oligos in a similar manner, ZBP1’s ZBDs have different binding modes for the m^8^G4 r(CpG)_3_, d(CpG)_3_, and the r(CpG)_3_ and LNA oligos. When Zα1^ZBP1^ was incubated with the m^8^G4 r(CpG)_3_ oligo, amide signals experienced large chemical shifts, similar to Zα^ADAR1^’s titration with the same oligo (Figure 4E). Of note, Zα1’s critical Tyr50 residue in the recognition helix (Figure 4E inset) and Ala64, which occupies a structurally similar position as Zα^ADAR1^’s double proline motif [9], have extreme downfield chemical shifts in the proton dimension, indicating the presence of significant deshielding effects likely caused by the close packing of these residues alongside the negatively charged backbone of the m^8^G4 oligo (Figure 5B). Interestingly, both Zα^ADAR1^ and Zα1^ZBP1^ have nearly identical CSP magnitudes and directions for their respective Tyr residues, indicating highly similar binding modes to the Z-form oligo. Unlike the m^8^G4 r(CpG)_3_ oligo, the Zα1^ZBP1^ – d(CpG)_3_ titration series has very different CSPs compared to ADAR1’s Zα domain. Many residues within the interaction interface, including Tyr50, have distinct bimodal binding modes that shift directions after the 1:1 titration point (Figure 4F). At low nucleic acid concentrations, the magnitudes and directions of the chemical shifts roughly match the m^8^G4 titration series, indicating the first binding mode is to the Z-conformation. As the titration reaches equivalent protein:nucleic acid concentrations, most peaks in the interaction interface shift directions, suggesting the presence of a second binding mode. Despite the appearance of a second binding mode, Zα1^ZBP1^ contacts both oligos using a similar interface, albeit with much lower CSPs, especially in the β-wing for the DNA series. Overall, Zα1^ZBP1^’s CSPs were 2.7x and 3.4x lower in the d(CpG)_3_ titration series compared to the m^8^G4 r(CpG)_3_ series across the entire domain and the interaction interface, respectively.

Unlike Zα^ADAR1^, which had similar chemical shift magnitudes and directions for all convertible oligos, the Zα1^ZBP1^ – r(CpG)_3_ titration series was unique compared to both the DNA and m^8^G4 r(CpG)_3_ series (Figure 4G). As with the m^8^G4 series, the unmodified r(CpG)_3_ titration series showed only one set of linear shifts, indicating a single binding mode. However, the chemical shifts were much smaller than the m^8^G4 modified RNA oligo and had significantly different final positions. Average CSPs at the 1:4 titration point were 1.8x and 2.5x lower across the entire domain and within the interaction interface, respectively, when compared to the modified Z-RNA oligo (Figure 5B). Additionally, Zα1^ZBP1^ uses a slightly modified interaction interface to contact r(CpG)_3_ oligos compared to the m^8^G4 and DNA 6-mers. Residues within the α3-helix more sparsely contact the r(CpG)_3_ oligo, however, much like Zα^ADAR1^, Zα1’s β-wing is more involved in binding the RNA oligos compared to DNA oligos. Consistent with the inability of any ZBP1 construct to convert RNA to the Z-form, Zα1^ZBP1^ has nearly identical binding modes to the LNA and unmodified RNA 6-mers (Figure 4H). Interestingly, the final positions of both A-form duplexes closely match the final positions of the second binding mode seen in the bimodal DNA shifts (Figure 4F-H). This suggests that at saturating (>2:1) protein:nucleic acid stoichiometries, Zα1^ZBP1^ fully stabilizes the DNA in the Z-conformation and shifts towards an initial left-handed binding mode (Supplementary Figure 9). As nucleic acid concentrations increase up to an equimolar stoichiometry, Zα1 domains begin to be titrated off and lose the ability to fully stabilize the Z-conformation. Increasing nucleic acid concentrations further results in the reversion of the DNA conformation back to the B-form and the appearance of the second, right-handed binding mode. This interpretation is consistent with the previous CD results, which showed that Zα1^ZBP1^ was less efficient than Zα^ADAR1^ and ZαZβ^ADAR1^ in stabilizing the short 6-mer DNA in the Z-form. It is currently unclear if the reversion to the B-conformation occurs while still bound to Zα1^ZBP1^ or if the dissociation of Z-form oligos from Zα1^ZBP1^ results in the reversion.

Zα1^ZBP1^’s interactions with the different 6-mer nucleic acids also show the presence of different time regimes compared to Zα^ADAR1^. Amide signals in the Zα1^ZBP1^ – m^8^G4 r(CpG)_3_ still closely match the behaviors seen in the Zα^ADAR1^ series with most peaks experiencing fast exchange and residues in the interaction interface experiencing additional contributions from intermediate exchange processes (Supplementary Figure 7E). Similar to the ADAR1 series, Zα1^ZBP1^ also experiences increased contributions from intermediate exchange in the d(CpG)_3_. Still, these contributions are limited to only the residues in the interaction interface with the rest of the domain largely experiencing fast exchange. However, unlike Zα^ADAR1^, amide signals in the interaction interface of Zα1^ZBP1^ remained broadened throughout the titration with the DNA 6-mer and intensity did not increase at the 1:2 and 1:4 titration points (Supplementary Figure 7F). The increased intermediate exchange seen at later titration points is likely caused by the constant exchange between free/bound nucleic acid as a result of the formation of unstable complexes. Similar to the CSP results, the time regimes of both the r(CpG)_3_ and LNA (CpG)_3_ are remarkably similar to each other (Supplementary Figure 7G-H). Both oligos shift almost exclusively in the fast time regime with no additional intermediate exchange contributions in the interaction interface, consistent with the domain’s inability to flip either LNA or RNA. Interestingly, for some peaks in the Zα1^ZBP1^ – LNA (CpG)_3_ titration series, there was evidence for both fast and slow exchange (Supplementary Figure 8C). Amide signals shifted linearly indicating fast exchange processes, however, there was notable splitting of the peaks at the intermediate titration points, consistent with two different species or slow exchange. The ^15^N-HSQCs of free Zα1^ZBP1^ has two unique sets of peaks because of slow exchange caused by the *cis*-*trans* isomerization of a proline residue [57], however, this is likely a separate phenomenon as the split peaks have roughly the same signal intensity as opposed to the different populations of *cis*-versus *trans*-proline seen in the free spectra (~11% v 89%, respectively).

### Zα2^ZBP1^ and Tandem^ZBP1^ Behave Nearly Identically to Zα1^ZBP1^ by NMR

Mirroring the Circular Dichroism results, the Zα2^ZBP1^ and Tandem^ZBP1^ ^15^N-HSQC titration series are largely similar to the Zα1^ZBP1^ titration series with some notable differences (Supplementary Figure 10). Like Zα1^ZBP1^, Zα2^ZBP1^ shows large chemical shifts with the m^8^G4 oligo with similar magnitudes and directions to the Zα^ADAR1^ – m^8^G4 r(CpG)_3_ titration series, indicating similar binding to the Z-form oligo (Supplementary Figure 10A). Zα2’s interaction interface with Z-form oligonucleotides differs slightly from the canonical ZBD binding mode as the β1-sheet plays a more significant role in contacting the Z-form (Figure 5C, Supplementary Figure 1E). The crystal structure of Zα2 with Z-DNA shows that Arg124 makes a water mediated hydrogen bond to the phosphate backbone [10], and similarly, we observed large CSPs around Arg124 in the m^8^G4 r(CpG)_3_ titration series (Figure 5C). Like Zα1^ZBP1^, Zα2^ZBP1^ displayed bimodal binding behaviors with the d(CpG)_3_ oligos, consistent with Zα2^ZBP1^ forming unstable complexes with short 6-mer DNA sequences. Likewise, the magnitudes of Zα2^ZBP1^’s CSPs to the d(CpG)_3_ ligand were much lower across both the entire domain and the interaction interface when compared to the Z-RNA ligand (2.1x and 2.6x, respectively). Zα2^ZBP1^ also displays very similar CSP magnitudes and directions for both the RNA and LNA ligands. Much like Zα1^ZBP1^, Zα2^ZBP1^ uses a slightly modified interaction interface to contact the A-form oligos, in which the β1-sheet is no longer used and the CSPs are more evenly distributed along the entire length of the recognition helix (Figure 5C). Additionally, residues at the N-terminus of the α1-helix experience large CSPs with the RNA and LNA oligos. This portion of the α1-helix is in close proximity to the 3_10_-helix at the N-terminus of the recognition helix in its crystal structure [10]. Thus, the high CSPs in this region are likely a propagation of structural perturbations made to the α3-helix. Interestingly, titration of all oligos except the DNA 6-mer resulted in significant quenching of the intermediate exchange seen in the 3_10_-helix (Supplementary Figure 11). As opposed to signal intensities in the rest of the domain, Ala137 and Lys138 peak heights increased with increasing concentrations of the m^8^G4 r(CpG)_3_, r(CpG)_3_, and LNA (CpG)_3_ oligos, suggesting that binding to these nucleic acids preferentially stabilized one of the states being sampled by the 3_10_-helix. Chemical shift patterns were significantly different for the m^8^G4 r(CpG)_3_ oligo (Supplementary Figure 11A) compared to the A-form r(CpG)_3_ and LNA (CpG)_3_ oligos (Supplementary Figure 11C & D, respectively), indicating that the two states are influenced by the binding of right-versus left-handed oligos. Similar results were seen by Ha et. al., in which the 3_10_ helix rotates to engage the phosphate backbone of the Z-conformation in the crystal structure compared to the position of the 3_10_-helix seen in the Apo NMR structure [10, 54]. The titration of d(CpG)_3_ into Zα2^ZBP1^ did not result in the quenching of the intermediate exchange (Supplementary Figure 11B); however, this is likely due to the wide-spread intermediate exchange experienced by the rest of the domain as a consequence of forming unstable complexes with the short 6-mer DNA (Supplementary Figure 10A). Outside of the 3_10_-helix, Zα2^ZBP1^ also experienced very similar exchange regimes to Zα1^ZBP1^ for all of the oligo titrations (Supplementary Figure 7I-L), including split peaks in the interaction interface indicative of both slow and fast exchange contributions (Supplementary Figure 8D).

Despite being in a tethered system, the individual domains of Tandem^ZBP1^ acted very similarly to their isolated Zα1^ZBP1^ and Zα2^ZBP1^ counterparts. CSP magnitudes and directions were nearly identical to the isolated ZBDs (Figure 5D), and bimodal shifts were similarly observed for the Tandem^ZBP1^ – d(CpG)_3_ titration series, although larger complex sizes made many of the bimodal shifts disappear completely by the 1:4 titration point (Supplementary Figure 10B). Interestingly, amide signals in the Zα1 domain of the Tandem^ZBP1^ – m^8^G4 r(CpG)_3_ titration series had much larger CSPs compared to amide signals in the Zα2 domain at early titration points. Zα2 peaks did not experience large CSPs until after the 1:1 titration point, suggesting that Zα1 has a higher apparent affinity for Z-form nucleic acids compared to Zα2 (Figure 5D). This pattern was not observed for either the d(CpG)_3_ or r(CpG)_3_ oligos, suggesting that the apparent affinities for the right-handed B- and A-conformations are similar between the two domains. The exchange regimes were also similar between the isolated and tandem domains for each titration series, however, precise analysis of Tandem^ZBP1^ was difficult due to the much larger complex size (Supplementary Figure 7M-O). Similar to the Zα2^ZBP1^ titrations, addition of oligos to Tandem^ZBP1^ quenched the intermediate exchange caused by exchange of the 3_10_-helix between multiple states.

### 1D Imino Proton Spectra Confirm the Adoption of A-, B-, or Z-conformations Throughout ZBD Titrations

Figure 6A-D shows changes in the 1D imino proton spectra of the m^8^G4 r(CpG)_3_, d(CpG)_3_, r(CpG)_3_, and LNA (CpG)_3_ oligos upon titration of the different ADAR1 and ZBP1 ZBD constructs. The imino resonances were assigned using previous assignments from studies of the d(CpG)_3_ and r(CpG)_3_ oligos [78] and LNA (CpG)_3_ oligos [71]. Assignments of the free m^8^G4 r(CpG)_3_ imino resonances were inferred from their similarity to 1D imino spectra of the d(CpG)_3_ and r(CpG)_3_ oligos under saturating Zα^ADAR1^ concentrations, assuming that the m^8^G4 r(CpG)_3_ oligo is adopting a substantial Z-form population under our experimental conditions. Similar to previous reports [38], the m^8^G4 r(CpG)_3_ is transiently sampling both the A- and Z-conformations or a melted intermediate at 35°C in the absence of protein as the imino resonances for G2 is nonexistent and G4 is heavily broadened, indicating accessibility to and exchange with water which decreases signal intensity (Figure 6A). The addition of any of the ZBD constructs to m^8^G4 r(CpG)_3_ resulted in a relative increase in the G4 imino signal, and the potential appearance of G2 at early titration points when protein saturating conditions allowed for the full stabilization of the duplexed RNA and the quenching of exchange with water. m^8^G4 r(CpG)_3_ imino resonances for all ZBD titrations additionally displayed fast exchange regimes throughout the titration, which largely match the exchange regimes seen for the 2D ^15^N-HSQC titration series, confirming the relatively fast interconversion of the m^8^G4 modification between the A- and Z-conformations. Interestingly, increasing relative concentrations of Zα^ADAR1^ did not cause significant CSPs in the G4 resonance, possibly indicating the adoption of similar Z-conformations in the presence and absence of protein. However, relative increases in Zα1^ZBP1^ or Zα2^ZBP1^ concentrations did result in significantly different downfield and upfield imino resonances, respectively, potentially suggesting different free- and protein-bound Z-RNA conformations. Addition of Tandem^ZBP1^ to the m^8^G4 r(CpG)_3_ oligo resulted in minor downfield shifts perturbations to the G4 imino resonance. The lower overall shifts are likely due to the averaging of Zα1 and Zα2 induced changes; however, Zα1 seems to prefer the already stabilized Z-form, mirroring the ^15^N-HSQC analyses.

**Figure 6.**
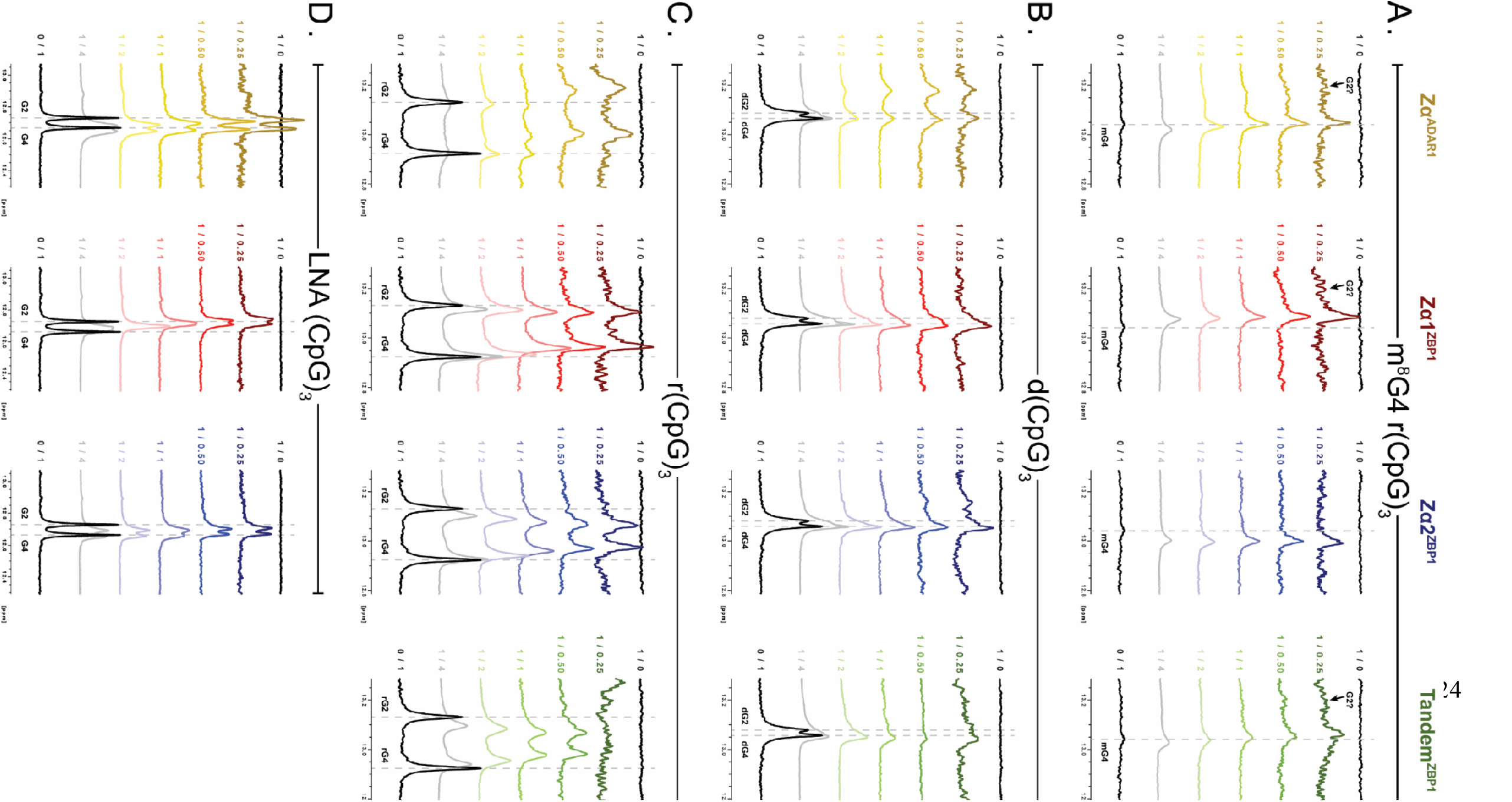
1D imino spectra of different 6-mer oligos upon titration of Zα^ADAR1^ (yellow), Zα1^ZBP1^ (red), Zα2^ZBP1^ (blue), and Tandem^ZBP1^ (green) measured at 35°C. Each spectrum was artificially scaled up/down to account for differences in nucleic acid concentrations and protein/nucleic acid ratios are listed to the left of each spectrum. Grey dashes represent the starting positions of the resonances in the free spectra. **(A)** 1D imino titration series of ZBD constructs with the m^8^G4 r(CpG)_3_ oligo. The resonances of the Z-form oligo are labeled as mG4. **(B)** 1D imino titration series of ZBD constructs with the d(CpG)_3_ oligo. The resonances of the B-form oligo are labeled as dG2 and dG4. **(C)** 1D imino titration series of ZBD constructs with the r(CpG)_3_ oligo. The resonances of the A-form oligo are labeled as rG2 and rG4. **(D)** 1D imino titration series of ZBD constructs with the LNA (CpG)_3_ oligo. The resonances of the A-like oligo are labeled as G2 and G4.

In contrast to the m^8^G4 imino signal intensities, the addition of Zα^ADAR1^, Zα1^ZBP1^, Zα2^ZBP1^, or Tandem^ZBP1^ resulted in a relative decrease of the signal intensities of imino resonances for the d(CpG)_3_, r(CpG)_3_, and LNA (CpG)_3_ oligos (Figure 6B-D). The oligos in the B-form, A-form, and A-like conformations, respectively, are relatively stable under these experimental conditions and do not adopt appreciable Z-form populations or melted intermediates in the absence of protein, even at 42°C [38]. The decrease in imino proton signals could reflect increased exchange with water as a result of active right-to-left-handed conversion processes in the presence of the ZBDs or increased R2 relaxation as a result of increased complex size or increased μs-ms exchange contributions due to active conversion processes. Similar to previous reports [78], titration of Zα^ADAR1^ into the d(CpG)_3_ oligo resulted in the gradual separation of the B-form G2 and G4 imino resonances into new Z-form positions at ~13.20 and ~13.05 ppm, respectively (Figure 6B). Interestingly, G4, but not G2, imino resonances in the Zα1^ZBP1^–d(CpG)_3_ titration series shifted to a position similar to that observed in the Zα^ADAR1^ series. Instead, the G2 resonance gradually broadened as [P]/[NA] ratios were increased. Titration of Zα2^ZBP1^ with the d(CpG)_3_ oligo, in contrast, did result in the appearance of two unique Z-form G2 and G4 resonances at high [P]/[NA] ratios. Tandem^ZBP1^ largely behaved similarly to Zα2^ZBP1^, however, analysis was hampered due to heavy line broadening, likely because of increased molecular size. The increased ability of Zα2^ZBP1^ and Tandem^ZBP1^ to induce a more stable Z-conformation G2 resonance than Zα1^ZBP1^ reflects the CD results that showed that Zα2^ZBP1^ was more efficient in stabilizing the Z-conformation. Furthermore, compared to Zα1^ZBP1^ and Zα2^ZBP1^, the imino resonances of the Zα^ADAR1^ – d(CpG)_3_ titration series were significantly more broadened, indicating more substantial contributions from intermediate exchange, mirroring the results of the 2D ^15^N-HSQC analyses.

Likewise, the exchange regimes of the imino resonances for the 1D Zα^ADAR1^ – r(CpG)_3_ titration series matched the slow exchange regime seen in the ^15^N-HSQCs. Peak positions for the B-form G2 and G4 resonances gradually disappeared and reappeared in the Z-form positions at ~13.2 and ~13.0 ppm, respectively, as relative [P]/[NA] ratios were increased and closely matched the final peak positions of other studies (Figure 6C, [78]). Similarly, exchange regimes for the 1D titration series of Zα1^ZBP1^, Zα2^ZBP1^, and Tandem^ZBP1^ with the r(CpG)_3_ oligo matched the ^15^N-HSQC series and occurred almost exclusively on the fast time scale with relatively minor signal loss, with the exception of Tandem^ZBP1^, which experienced larger signal losses. Additionally, while imino resonances for the Zα^ADAR1^ series resulted in downfield shifts, the ZBP1 constructs displayed different final peak positions for the RNA 6-mer imino resonances, confirming a different binding mode and lack of converting ability for the unmodified r(CpG)_3_ oligo. Imino shifts for the LNA (CpG)_3_ oligo also largely match the trends seen in the ^15^N-HSQCs (Figure 6D). G2 and G4 imino resonances gradually shift throughout the titration series, indicating fast on/off rates, however, unlike every other 1D titration series, largest CSPs occurred at low relative [P]/[N] ratios and gradually shifted back to the positions of free oligo at high relative [P]/[N] ratios. In line with previous reports [71], imino resonances were also heavily broadened at low relative [P]/[N] ratios, and with the multimodal binding behaviors seen in the 2D ^15^N-HSQC spectra indicate that multiple LNA (CpG)_3_ oligos are capable of binding the same Zα^ADAR1^ protein. Interestingly, Zα1^ZBP1^ and Zα2^ZBP1^ show slightly different behaviors in the 1D series, with the G4 imino resonance shifting linearly as relative [P]/[N] ratios increase, whereas the G2 imino resonance has the largest CSP at low relative [P]/[N] ratios and shift back towards free oligo at high relative [P]/[N] ratios. Unlike Zα^ADAR1^, where signal intensities increased as relative protein concentrations increased, increasing the relative concentrations of both Zα1^ZBP1^ and Zα2^ZBP1^ caused a decrease in the height of the imino resonances, suggesting that both domains do not have multiple binding sites for LNA oligos.

### Isothermal Titration Calorimetry Captures Active A/B-to-Z Conversion Processes

From CD and NMR experiments, it is clear that ZBP1’s ZBDs behave much differently than those of ADAR1’s ZBDs due to differences in conversion activities and binding modes. Previous studies have used ITC to investigate the properties of different ZBDs binding to DNA oligos and have shown the method’s potential for capturing active conversion processes [79], dependent on experimental design. ITC is the method of choice for assessing and obtaining thermodynamic parameters such as enthalpy (ΔH), entropy (ΔS), free energy (ΔG), equilibrium binding affinity (K_a_), and stoichiometry [80, 81]; however, modeling the interaction of ZBDs with different nucleic acids is extremely difficult due to the inherent complexity of the system. Most experimental studies suggest that, at least in vitro, ZBDs bind to both left-handed and right-handed oligos and act through an induced fit sequential model (Figure 7; [39–42, 78, 82]). Thus, ITC models must account for the relative affinities of ZBD binding to multiple, potentially non-equivalent, sites on both right- and left-handed duplexes as well as consider the interconversion rates between A/B- and Z-forms for each type of protein-nucleic acid complex. Despite the difficulty in extracting reliable parameters, we set out to measure the titration of each ZBD construct into the m^8^G4 r(CpG)_3_, d(CpG)_3_, and r(CpG)_3_ 6-mer oligos to capture initial binding and conversion processes.

**Figure 7.**
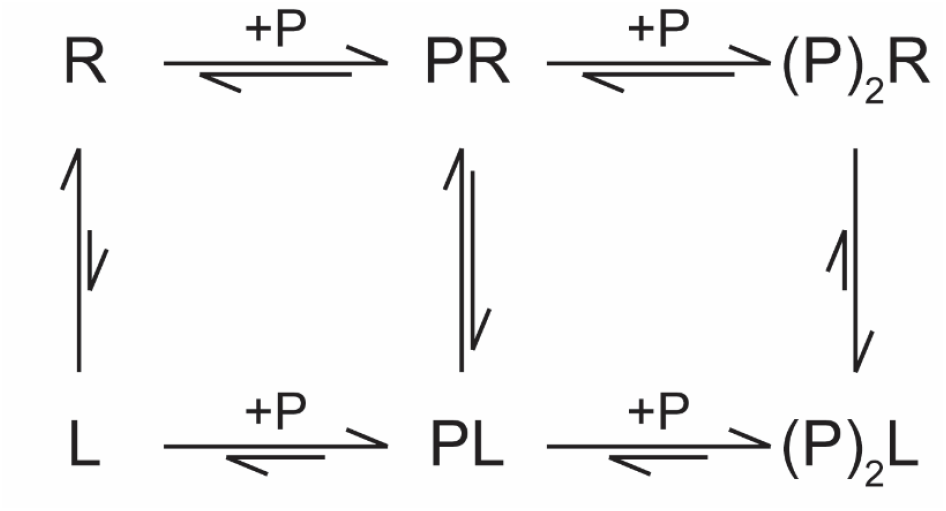
Induced fit sequential binding model describing the binding of ZBDs (denoted as P) to nucleic acids including conformational changes of right-handed (R) duplexes to the left-handed (L) Z-conformation. Equilibria arrows are representative of relative population dynamics and favorabilities but do not represent measured values.

Titration of Zα^ADAR1^ into all three oligos resulted in large heat releases typical for enthalpically favorable interactions (Figure 8A, top panel); however, the amount of heat released and the shape of the resulting isotherms varied greatly between the DNA and RNA 6-mers. Titration into the pre-stabilized Z-form m^8^G4 r(CpG)_3_ oligo resulted in an initial heat of binding of approximately −29 kcal•mol ^−1^, whereas the d(CpG)_3_ and r(CpG)_3_ oligos had much lower (approximately −9 kcal•mol^−1^) initial heats of binding. Additionally, the resulting exotherms differed significantly between all three oligos. The Zα^ADAR1^ – m^8^G4 r(CpG)_3_ exotherm was monophasic and saturated at a molar ratio of 2. In contrast, the d(CpG)_3_ and r(CpG)_3_ titration series were biphasic and consisted of two distinct exotherms that saturated at molar ratios of 1 and 2, respectively. (Supplementary Figure 12A-C). ZBD binding to a pre-formed Z-NA 6-mer should result in two equivalent binding sites and would therefore explain the monophasic nature of the m^8^G4 titration series. The presence of two distinct isotherms in the d(CpG)_3_ and r(CpG)_3_ titration series suggests sequential, non-equivalent binding sites in which the first site must be saturated to allow for binding of the second site.

Although both the d(CpG)_3_ and r(CpG)_3_ oligos had two distinct isotherms, the slopes of the first isotherm differed greatly between them. Titration of Zα^ADAR1^ into d(CpG)_3_ resulted in a shallower isotherm with smaller differences (<1 kcal•mol^−1^) in the heats of binding across the entire first isotherm, whereas titration into r(CpG)_3_ resulted in much larger differences (~4 kcal•mol^−1^). These differences could be reconciled by RNAs having a higher activation energy for adopting the Z-form. The energy of Zα^ADAR1^ binding right-handed oligonucleotides is partially co-opted to deform nucleic acid duplexes into a melted intermediate or deformed structure, where more binding energy is lost in the conversion of RNA compared to its DNA counterpart. Interestingly, this initial binding event must be saturated before the second isotherm appears, as doubling the oligo length doubles the molar ratio at which each isotherm is saturated (Supplementary Figure 13). Additionally, since the m^8^G4 modification results in a significant Z-form population at 35°C, the binding energy is much larger than for the right-handed oligos as no energy is lost in the destabilization of the duplex structure. The secondary structures of each oligo were confirmed to be in the Z-conformation at the end of the experiment by immediately transferring the contents of the ITC cell to the CD spectrophotometer (Figure 8A, bottom panel). Like previous CD results, Zα^ADAR1^ converted all three 6-mer oligos to the Z-form. These findings suggest that under our experimental conditions, the ITC experiments capture Zα^ADAR1^’s initial binding event to A/B-form oligos and its stoichiometry-dependent conversion to the Z-conformation at [protein]:[binding site] molar ratios greater >1. Unfortunately, reliable affinities could not be extracted from the raw isotherms for various reasons, including the presence of overlapping isotherms and lower active fractions of the duplex due to measurements occurring at 35°C. Lowering the measurement temperature nominally increases the active fraction of stable duplex but requires unfeasibly long injection intervals to allow for full conversion of the oligonucleotides (Supplementary Figure 13).

**Figure 8.**
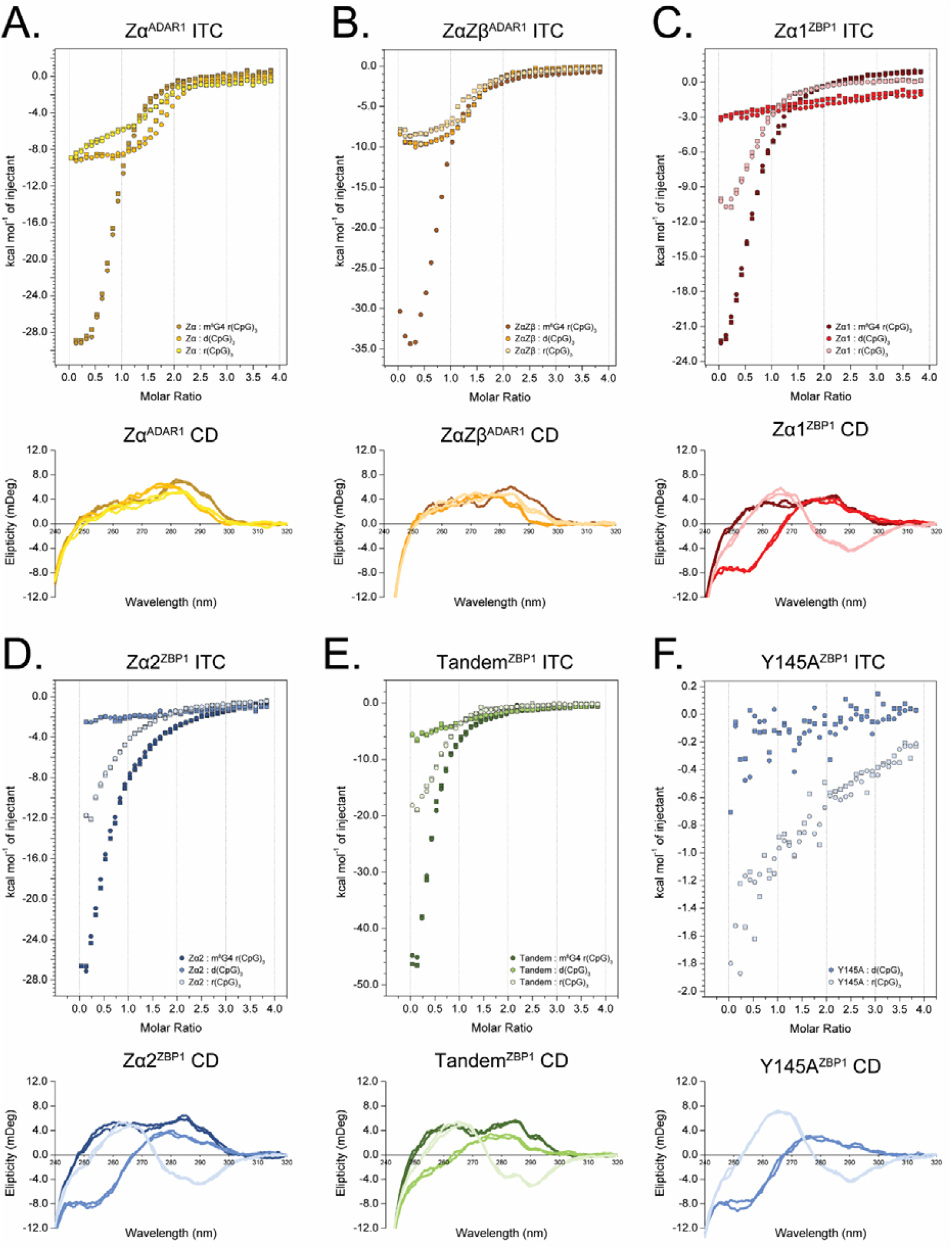
Isothermal Titration Calorimetry isotherms of Zα^ADAR1^ **(A)**, ZαZβ^ADAR1^ **(B)**, Zα1^ZBP1^ **(C)**, Zα2^ZBP1^ **(D)**, Tandem^ZBP1^ **(E)**, with the m^8^G4 r(CpG)_3_, d(CpG)_3_, and r(CpG)_3_ oligos, and Y145A^ZBP1^ **(F)**, with the d(CpG)_3_ and r(CpG)_3_ oligos. All Measurements were performed at 35°C and are colored as indicated. ITC measurements (top panels) were measured in duplicate with individual points represented as circles and squares. CD measurements (bottom panels) were taken immediately after each ITC measurement to confirm the conformation of the nucleic acid.

Titration of ZαZβ^ADAR1^ into the three oligos resulted in isotherms that were largely similar to those of Zα^ADAR1^ with a few noticeable differences. Like Zα^ADAR1^, titration of ZαZβ^ADAR1^ into m^8^G4 r(CpG)_3_ resulted in a similar large heat release (−35 kcal•mol^−1^), however, during the first few injections, ZαZβ^ADAR1^ manifests an initial endothermic phase followed by an exothermic phase. The initial endothermic phase may be indicative of conformational changes in the tandem domain construct as the Zα domain initially binds the nucleic acid or could arise from dissociation of ZαZβ^ADAR1^ molecules at lower concentrations, as the construct is prone to LLPS at high concentrations (data not shown). The initial endothermic phase is likely an inherent feature of ZαZβ^ADAR1^, as it is present in the d(CpG)_3_ and r(CpG)_3_ measurements as well (Figure 8B). Titration of ZαZβ^ADAR1^ into the DNA and RNA 6-mers also produced two exotherms, similar to Zα^ADAR1^, but the slopes of the first exotherm were much more similar in magnitude, suggesting that the presence of the linker or the Zβ domain reduces the energy barrier for adopting the Z-form for the r(CpG)_3_ oligo. CD subsequently confirmed that all three oligos were adopting the Z-conformation by the end of the ITC experiment. For the same reasons listed above, affinities for the ZαZβ^ADAR1^ titration series were not extracted.

### ZBP1 Binds Pre-Stabilized Z-form, but not A/B-form, Duplexes with Similar Thermodynamics as ADAR1

Like previous CD and NMR results, ITC showed that Zα1^ZBP1^, Zα2^ZBP1^, and Tandem^ZBP1^ bound the 6-mer oligo constructs with similar thermodynamic properties but differed greatly from the ADAR1 constructs, with the exception of the pre-stabilized m^8^G4 r(CpG)_3_ oligo. Both Zα1^ZBP1^ and Zα2^ZBP1^’s titration into the m^8^G4 RNA resulted in large exothermic heat releases to a similar extent as ADAR1’s isolated Zα domain (Figure 8C,D), however, while Zα1^ZBP1^ titration series was nearly saturated at a molar ratio of 2, titration of Zα2^ZBP1^ continued to release heat at much higher molar ratios, suggesting nonspecific multivalent interactions occurring at high protein concentrations consistent with the phase separating properties of the domain. Titration of Tandem^ZBP1^ into m^8^G4 r(CpG)_3_ resulted in a much larger heat release (Figure 8E), nearly equivalent to the addition of both Zα1^ZBP1^’s and Zα2^ZBP1^’s combined exothermic contributions, indicating that both domains are binding simultaneously. However, the presence of both domains does not affect the stoichiometry at which the isotherm saturates potentially indicating a different mechanism for the exothermic interaction. It is possible that at early titration points, both domains bind Z-form m^8^G4 oligos in a noncooperative fashion. As titrations continue, one of the ZBDs – likely Zα1 – outcompetes the other for binding. CD confirmed that all three constructs stabilized the modified RNA in the Z-form.

Interestingly, unlike the ADAR1 constructs, titration of Zα1^ZBP1^, Zα2^ZBP1^, or Tandem^ZBP1^ into the d(CpG)_3_ oligo resulted in extremely shallow isotherms with low exothermic contributions (Figure 8C-E). Although shallow isotherms typically signify weak or nonspecific interactions, our CD and NMR results demonstrated significant, albeit unstable, complex formation occurring between ZBP1 ZBDs and d(CpG)_3_ oligos. Given that ADAR1’s ZBDs co-opt initial binding energies into destabilizing duplex structures for subsequent A/B-to-Z conversion, ZBP1’s ZBDs may be doing something similar; however, the high Z-form energy barrier and the relatively unstable ZBP1-DNA complexes result in low measurable heat releases. Furthermore, all ITC measurements were carried out at 35°C to facilitate efficient conversion within reasonable injection intervals, however, ITC measurement temperatures were much closer to the measured T_M_’s (~50°C; [38]), than the previous CD measurements. Although Zα^ADAR1^ and ZαZβ^ADAR1^ are able to stabilize ZBD – Z-DNA complexes at these temperatures, ZBP1’s lower Z-stabilization efficacies result in weak complexes that readily disassociate, which would result in lower overall heat release. CD confirmed the unstable complexes, as the DNA was predominantly in the B-conformation at 35°C with minor perturbations. Interestingly, Tandem^ZBP1^’s titration into d(CpG)_3_ resulted in a more defined isotherm and greater Z-stabilization by CD, suggesting that the constructs’ ZBDs are working cooperatively to stabilize the Z-form under the ITC experimental conditions. Additionally, titration of the Z-binding deficient Y145A^ZBP1^ construct into the DNA 6-mer resulted in an even lower exothermic isotherm (<-1 kcal•mol^−1^) that is more indicative of non-specific interactions with the DNA (Figure 8F).

Titration of Zα1^ZBP1^, Zα2^ZBP1^, and Tandem^ZBP1^ into the r(CpG)_3_ resulted in biphasic isotherms with an initial, short endothermic component followed by a larger exothermic component (Figure 8C-E). Unlike the endothermic components of ZαZβ^ADAR1^’s titration with all three oligos, the initial endotherms of ZBP1’s ZBD constructs is unlikely to be a protein specific phenomenon. Instead, the initial endothermic event suggests a partial desolvation of A-form duplexes due to the titration of the protein, mimicking salting-out or counter-ion stripping. Additionally, the shapes of the r(CpG)_3_ isotherms with the ZBP1 constructs closely match previous titration measurements of Zα^ADAR1^ into an LNA (CpG)_3_ oligo [71], further supporting an A-form-dependent endothermic contribution. Unlike d(CpG)_n_ repeats, ZBP1 is incapable of stabilizing r(CpG)_n_ repeats in the Z-conformation. The lack of A-to-Z conversion ability results in a single exotherm representative of the ZBDs binding to an A-form duplex and confirmed by CD at the end of the measurement. Similar to the d(CpG)_3_ oligo, mutation of Zα2’s tyrosine resulted in a severely abrogated interaction with the r(CpG)_3_ duplex (Figure 8F). Although ZBP1’s ZBDs lack apparent RNA conversion activities, the addition of the domains results in lower A-form characteristic NMR spectral peaks, potentially suggesting that the domains are still partially competent in deforming native A-form duplexes at 35°C. This in agreement with previous reports that flipping-deficient ZBDs promote the adoption of ssRNA intermediates [38].

### 8-Oxoguanine is a Likely Physiologically Relevant Target of ZBP1 Sensing and Activation

Although we have used 8-methylguanine modifications to promote and stabilize Z-form oligonucleotides, m^8^G nucleobase modifications are not prevalent within human RNA or DNA molecules. 8-oxoguanine, a modification with a similar propensity to promote Z-DNA in vitro [68], has recently been proposed to be a specific activator of ZBP1 when incorporated into released fragments of mitochondrial DNA after acute chemotoxic stress in vivo [83]. Similarly, acute or chronic exposure to the chemical stressors cisplatin and arsenic, which partially act through the generation of reactive oxygen species (ROS), greatly promotes the formation of Z-RNA in cells [51, 84]. 8-oxoguanine is the most prevalent oxidative product upon exposure to ROS, and due to the chemical reactivity afforded by the 2⍰OH and its predominant localization to the cytoplasm - and therefore vicinity to ROS production - guanines in RNA are more susceptible to producing o^8^G than guanines in DNA [85, 86]. Consequently, we wondered if ZBP1’s could bind and stabilize o^8^G4-modified RNA oligos similarly to the m^8^G4-modified RNAs.

To our surprise, 8-oxoguanine seems to be a greater natural Z-RNA stabilizer than the 8-methylguanine modification (Figure 9A). In the absence of protein (black line), the o^8^G4 r(CpG)_3_ was already predominantly in a Z-like conformation, even at 15°C. This is in contrast to the m^8^G4 r(CpG)_3_ oligo, which had distinct A- and Z-form populations (Figure 2F). Addition of any of the ZBDs did not drastically change the conformation of the o^8^G4 RNA duplex over time (Figure 9B), including the addition of Y145A^ZBP1^, further suggesting that the 8-oxoguanine stabilizes the Z-conformation to a greater extent than 8-methylguanine. Interestingly, from the CD timecourse, it appears that ZBP1’s ZBD constructs were ~20% more efficient than ADAR1’s constructs were in stabilizing the Z-form; however, without a clear change in the CD signal, it is possible that these differences represent minor changes in protein and nucleic acid concentrations between the experiments. Despite these limitations, it would be interesting to speculate that ZBP1’s ZBDs are specifically attuned to sensing o^8^G4-stablized Z-NAs. To confirm that ZBP1 is engaging the o^8^G4 stabilized Z-RNA in a similar manner to m^8^G4 Z-RNA, we additionally measured ^15^N-HSQC spectra for Zα1^ZBP1^ and Zα2^ZBP1^ at the 1:0.5 [ZBD]:[NA] titration point. Overlaying the resulting o^8^G4 r(CpG)_3_ ^15^N-HSQCs with the m^8^G4 r(CpG)_3_ titration series revealed that both Zα1^ZBP1^ and Zα2^ZBP1^ (Figure 9C,D, respectively) contact the o^8^G4 r(CpG)_3_ oligo in nearly the same manner. Similar CSP directions and magnitudes were seen for Zα1^ZBP1^ and Zα2^ZBP1^ at both 1:0.5 titration points, revealing that both domains are using the same interface and binding mode to both modified oligos. Due to the high prevalence of ROS under the same conditions that activate ZBP1 and the ability of 8-oxoguanine to specifically stabilize the Z-conformation, these results suggest that o^8^G adducts on RNA and DNA are the likely physiological ligand for ZBP1 sensing and activation.

**Figure 9.**
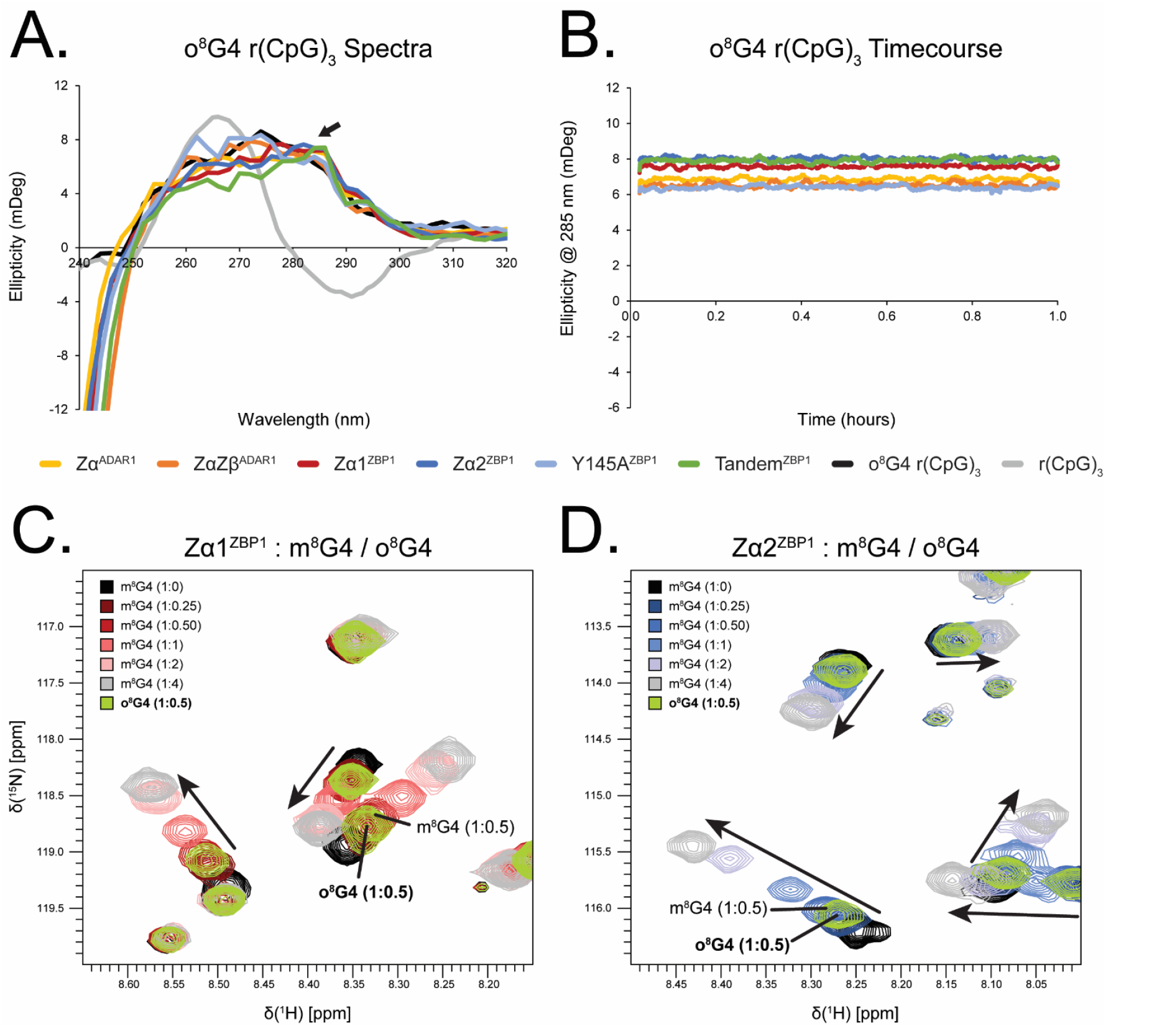
8-oxoguanine is a better Z-stabilizer than 8-methylguanine and is bound by ZBDs in a similar manner. **(A)** CD spectra of the o^8^G4 r(CpG)_3_ oligo in the presence of ADAR1 and ZBP1 ZBD constructs at 15°C, color coded according to the legend. Positive ellipticity at 285 nm (black arrow) indicates the presence of Z-form RNA. **(B)** CD time course measurements of ZBDs with o^8^G4 r(CpG)_3_ oligos. **(C)** Titration series of Zα1^ZBP1^ with m^8^G4 and o^8^G4 r(CpG)_3_ RNA. Titration points for both m^8^G and o^8^G modified oligos are color coded according to the legend above. **(D)** Titration series of Zα2^ZBP1^ with m^8^G4 and o^8^G4 r(CpG)_3_ RNA. Titration points for both m^8^G and o^8^G modified oligos are color coded according to the legend above.

## DISCUSSION AND CONCLUSIONS

ADAR1 and ZBP1 are both critical players of the innate immune response to viral infections, however, they promote drastically different cellular fates. ADAR1 is the master negative regulator of the response, in which it employs an adenosine deaminase domain to edit and destabilize dsRNA duplex structures and dsRBDs and ZBDs to sequester both A- and Z-NAs, respectively, thereby preventing dsRNA sensor activation and the initiation of IFN responses (Extensively reviewed in [1]). ZBP1, on the other hand, utilizes ZBDs to sense foreign nucleic acids in the Z-form to initiate downstream cell death pathways, including PANoptosis [2]. Mutation or depletion of ADAR1p150, the isoform containing the functional Zα domain, results in Z-RNA accumulation and ZBP1 activation. Despite having structurally homologous ZBDs, ADAR1 is able to suppress ZBP1 activation through ZBD-dependent sequestration of available Z-NAs; however, the mechanism by which ADAR1 is able to outcompete ZBP1 is poorly understood.

In this study, our CD, NMR, and ITC results demonstrate that contrary to prior beliefs, the ZBDs of ADAR1 and ZBP1 are not equivalent. The tandem ZαZβ domains of ADAR1 are thought to act as a single, bipartite domain that engage and convert nucleic acids to the Z-form in a cooperative manner [32]. Furthermore, the properties of the functional Zα domain are dramatically changed when expressed with the IDR linker and the Zβ domain. In contrast, the tandem ZBDs of ZBP1 do not appear to act cooperatively in vitro under our experimental conditions. NMR Relaxation experiments suggest that ZBP1’s ZBDs act as independent tethered domains and not as part of a larger bipartite unit, lacking any interdomain interactions. Additionally, CD, NMR, and ITC results suggest that Zα1 outcompetes Zα2 for binding to pre-stabilized Z-NAs and negatively modulates Zα2-dependent phase separation properties, in line with previous reports [21].

To our knowledge, this is the first report of ZBP1’s inability to convert native A-form RNA helices to the Z-form, and it calls into question the physiological ligands responsible for ZBP1’s activation and initiation of downstream signaling events. While ZBP1’s ZBDs are able to stabilize d(CpG)_n_ oligos in the Z-form, their conversion rates are much slower than ADAR1’s ZBDs and require the cooperativity afforded by longer sequences to reach nearly the same level of Z-conversion efficacy. It is likely that the *cis*-*trans* proline-proline, which are absent in both of ZBP1s Zα domains, is necessary for the conversion of r(CpG)_n_ oligos. Proline-to-alanine substitutions in ADAR1’s Zα domain, which cause one subtype of the AGS interferonopathy when paired with an ADAR-null allele [50], recapitulate ZBP1’s inability to convert RNA to the Z-form as well as hampering its DNA conversion activity. Although it is currently unclear what specific structural features of the proline-proline motif are responsible for its outweighed effect on RNA conversion over DNA conversion, previous studies have shown that the identity of residues within the β-wing greatly modulate conversion activities [47]. Crystal structures of ZBDs in complex with Z-RNA strands do not show any specific contacts between the prolines and the 2⍰OHs, making it unlikely that the motif is specifically affecting overall Z-RNA stability. However, disrupted β-wing positioning and/or the lack of specific contacts with A-form RNA may decrease initial energies of binding below the threshold necessary for A-to-Z conversion. The higher activation energy of Z-RNA (~1.2 kcal/(mol•bp) more than Z-DNA), implies that the loss of one or more favorable protein-nucleic acid interactions would abolish activity.

While not capable of flipping unmodified RNAs into the Z-conformation, lowering the activation energy with chemical modifications at the C8 position of guanine nucleobases allowed ZBP1 ZBD constructs to efficiently promote the Z-conformation to nearly the same extent as ADAR1 ZBDs. With the exception of Y145A^ZBP1^, which is incapable of stabilizing the Z-form, all ADAR1 and ZBP1 constructs contacted and stabilized using similar interfaces and with similar thermodynamic properties. Although we primarily used 8-methylguanine modifications to stabilize the Z-conformation in this study, we found that 8-oxoguanine, the most prevalent product of ROS exposure, is an even stronger inducer of Z-RNA. ZBP1 activation and initiation of cell death pathways has been implicated in cases of chronic or acute exposure to chemical stressors [51, 84], cancer/cell crisis [87, 88], ischemia-reperfusion [89, 90], viral infection [60, 91], and mitochondrial damage [92, 93], all of which are characterized by dysregulated mitochondrial function and elevated ROS. Thus, the accumulation of ROS and the generation of o^8^G modifications may allow for the widescale promotion and adoption of Z-RNA necessary for ZBP1 activation.

Other than chemical modifications, certain cellular conditions may also provide suitable ligands for ZBP1 activation. Recently, spliceosome inhibition and the generation of RNA:DNA hybrid duplexes have been shown to activate cell death pathways in a ZBP1-dependent manner [14, 94, 95]. Hybrid duplexes undergo a Zα^ADAR1^-dependent conformational change to the Z-form with much more favorable kinetics when compared to either DNA or RNA duplexes [96]. The lowered amount of 2⍰OH’s likely reduces the energetic barrier of Z-form adoption [3]; however, it is unknown whether ZBP1’s ZBDs would be able to effectively convert the hybrids to the Z-form. In addition to the formation of hybrid duplexes, negative supercoiling is a potent inducer of the Z-conformation [13, 97]. Processive enzymatic helicases, such as DNA and RNA polymerases, generate regions of negative supercoiling in their wake as they plow through dsDNA structures (extensively reviewed in [98]). Monoclonal antibody staining using the Z22 antibody subsequently showed the presence of polymerase- and torsional strain-dependent Z-DNA in vivo [99, 100]. Negative supercoiling in dsRNA would lower the energetic barrier for Z-form adoption and allow for ZBP1 binding and activation. Interestingly, MDA5, a dsRNA sensor with helicase activity, can induce negative helical torsion in long dsRNA substrates during oligomerization and in the absence of ATP hydrolysis [101]. It is interesting to speculate that MDA5 recognition of its physiological substrate may induce potential Z-forming regions for ZBP1 activation; however, there is currently no evidence supporting cooperative binding between MDA5 and ZBP1.

Additionally, our findings help explain how ADAR1p150 is able to prevent ZBP1’s activation purely through competitive binding [2], and why spontaneous activation of ZBP1 does not occur under homeostatic conditions (Figure 10). ZBP1’s ZBDs, being weaker Z-form converters and stabilizers, are easily outcompeted by ADAR1’s ZBDs for the low levels of available Z-prone RNA sequences in cells at steady-state. However, ongoing stressors, including infection or exposure to chemicals, generate ROS and increase the available pool of potentially Z-forming NAs. As exposure to the stressor continues or intensifies, the amount of Z-NAs increases to the point where ADAR1 is no longer able to sequester them away from ZBP1, leading to ZBP1 activation and initiation of cell death pathways. Thus, these two proteins might exist in a delicate balance in which ADAR1 works to salvage cells with low levels of viral load or oxidative damage, and ZBP1 removes cells that are too heavily damaged.

**Figure 10.**
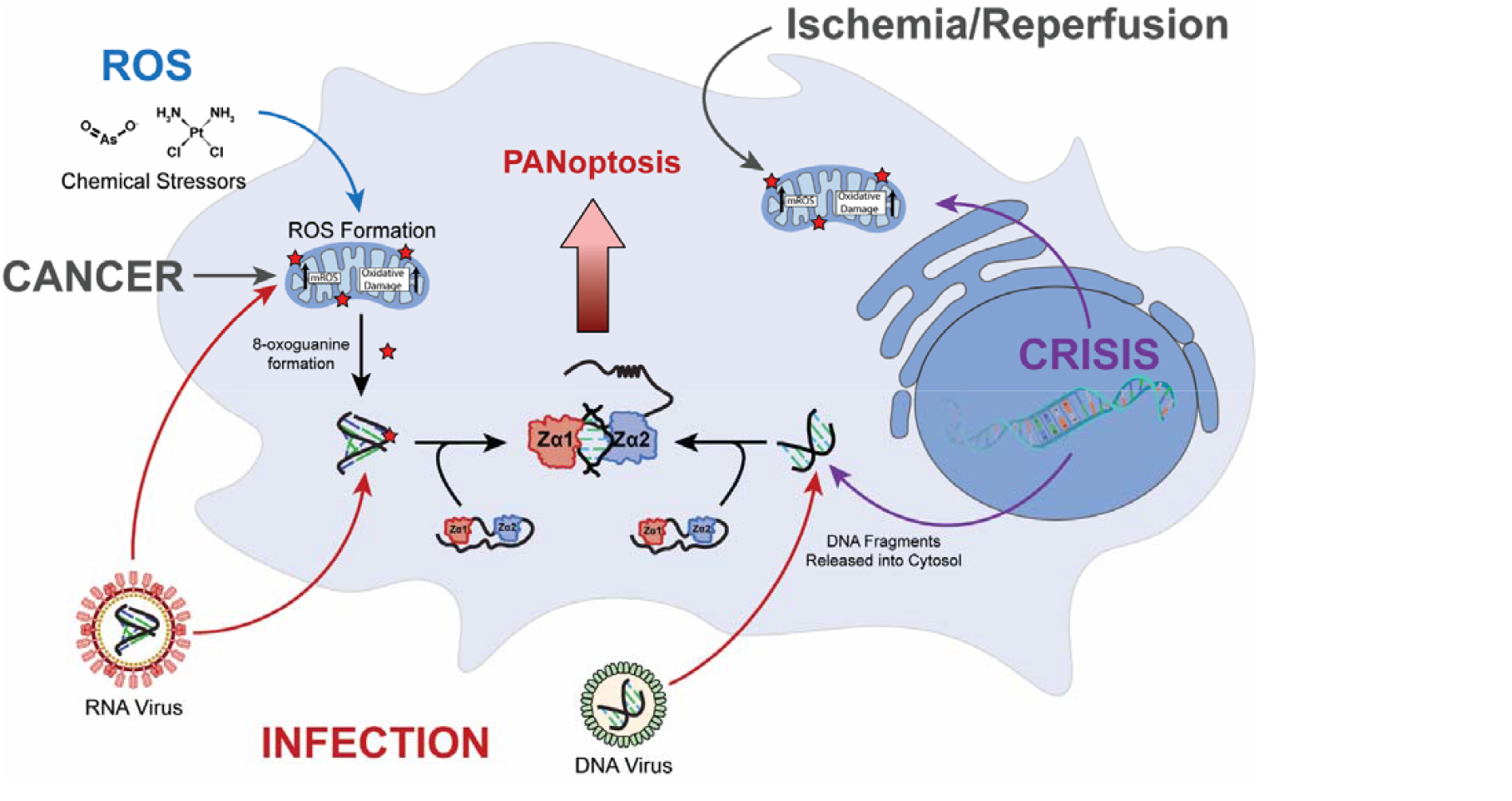
Mechanisms describing 8-oxoguanine production, Z-NA accumulation, and ZBP1 activation. Exposure to chemical stressors, cancer, infection, cellular crisis, ischemia/reperfusion injuries, and other stressors result in the direct production of reactive oxygen species or cause damage to mitochondria leading to the accumulation of ROS. O^8^G4 is the most prevalent oxidative product on nucleic acids and dramatically stabilizes Z-NAs. The accumulation of Z-NAs exceeds ADAR1’s ability to sequester them away from ZBP1, leading to ZBP1’s activation and initiation of cell death pathways.

## ASSOCIATED CONTENT

### Supporting Information

The Supporting Information is available free of charge via the Internet at biorxiv

Details of protein purification, sample preparation, conservation analysis, and CD, ITC, NMR experimental conditions/analysis.

## AUTHOR INFORMATION

### Authors

**Jeffrey B. Krall** *– Department of Biochemistry and Molecular Genetics, University of Colorado Anschutz Medical Campus, Aurora, Colorado, 80045, United States;*

**Lily G. Beck** – *Department of Biochemistry and Molecular Genetics, University of Colorado Anschutz Medical Campus, Aurora, Colorado, 80045, United States;*

**Parker J. Nichols** – *Department of Biochemistry and Molecular Genetics, University of Colorado Anschutz Medical Campus, Aurora, Colorado 80045, United States; Department of Biochemistry, University of Utah School of Medicine, Salt Lake City, Utah, 84112, United States;*

**Quentin Vicens** *– Department of Biology and Biochemistry, Center for Nuclear Receptors and Cell Signaling, University of Houston, Houston, Texas, 77204, United States;*

**Morkos A. Henen** – *Department of Biochemistry and Molecular Genetics, University of Colorado Anschutz Medical Campus, Aurora, Colorado, 80045, United States;*

### Author Contributions

The manuscript was written through contributions of all authors. All authors have given approval to the final version of the manuscript.

### Funding Sources

This project was supported by NSF grant #2153787 to B.V. and Q.V., and NIH grants R01 GM150642 to Q.V. and B.V., University of Colorado Cancer Center Support Grant P30 CA046934, and Biomedical Research Support Shared Grant S10 OD025020.

### Notes

The authors declare no competing financial interests.

## ACKNOWLEDGMENTS

The authors would like to thank current and former Vögeli Lab members and the Biophysics core (Dr. Robb Welty) and NMR facilities (Dr. David Jones) for thoughtful discussions and technical assistance.

## Notes

### Competing Interest Statement

The authors have declared no competing interest.

