## Supplementary Information for "ZBP1’s Inability to Convert Unmodified RNAs to the Z-form Underlies a Balanced Mechanism of RNA Recognition with ADAR1"

### Experimental Section:

#### Protein Purification:

The N-terminal Zα (139-201) domain of *Homo sapiens* ADAR1 (Zα$^{ADAR1}$) in the pet-28a(+) plasmid (N-terminal 6 × His-tag and thrombin cleavage site between His tag and the Zα$^{ADAR1}$ sequence) was a gift from Drs. Peter Dröge and Alekos Athanasiadis. Additionally, a tandem ZαZβ$^{ADAR1}$ (139-366) construct was similarly constructed. The codon optimized construct was synthesized by Genscript and subcloned into the same expression vector, pet-28a(+), using *Nde* I and *BamH* I restriction sites to yield proteins with an N-terminal 6 × His-tag and thrombin cleavage site. Four protein constructs from the N-terminus of *Homo sapiens* ZBP1 (UniProt: Q9H171): Zα1$^{ZBP1}$ (1-70), Zα2$^{ZBP1}$ (103-166), Y145A$^{ZBP1}$ (103-166; Y145A) and Tandem$^{ZBP1}$ (1-166) were generated. The codon optimized constructs were synthesized by Genscript and subcloned into the pet-28a(+) plasmid using *Nco* I and *BamH* I restriction sites to yield proteins with an N-terminal 6 × His-tag, thrombin cleavage site, and T7 tag.

The constructs were expressed and purified similarly to Beck et al., 2024[1]. Briefly, the plasmids were transformed and expressed in LOBSTR-BL21(DE3) *E. coli.* The cell cultures for all protein constructs except for Y145A$^{ZBP1}$ and ZαZβ$^{ADAR1}$ were grown in both M9 minimal media supplemented with 1 g/L $^{15}$N ammonium chloride and Luria Broth (LB); the cell cultures for Y145A$^{ZBP1}$ and ZαZβ$^{ADAR1}$ were grown only in LB media. All cell cultures were grown to an optical density of 0.6 at 600 nm, induced with IPTG at a final concentration of 1 mM, and allowed to express overnight at 21 °C. The cell cultures were centrifuged at 4k RPM for 15 minutes to collect the cell pellets. Pellets were resuspended in lysis buffer (50 mM Tris–HCl (pH 8.0), 300 mM NaCl, 10 mM imidazole), supplemented with 1 mM dithiothreitol (DTT) and 1 mM phenylmethylsulfonyl fluoride (PMSF), and subjected to sonication for cell disruption. The lysate was clarified by centrifugation at 15k RPM for 30 minutes to clear cell debris. The clarified supernatant was applied to 5 X 1 mL HisTrap FF columns (Cytiva, Marlborough, MA), washed with 5 column volumes (CVs) of lysis buffer, 5 CVs of wash buffer (50 mM Tris–HCl (pH 8.0), 1 M NaCl, 10 mM imidazole), and eluted in 1 mL fractions over 3 CVs of elution buffer (50 mM Tris–HCl (pH 8.0), 300 mM NaCl, 500 mM imidazole). The eluents were concentrated to ~4 mL and further purification was completed on a size-exclusion HiLoad 16/600 Superdex 75 pg (Cytiva, Marlborough, MA) in 20 mM potassium phosphate (pH 6.4), 100 mM NaCl, 1 mM DTT, 0.5 mM EDTA. All protein fractions were collected and concentrated to ~1-3 mM stocks using 3000 MWCO concentrator (Millipore-Sigma, Burlington, MA).

#### DNA and RNA Constructs and Preparations:

All DNA and LNA constructs were synthesized by Integrated DNA Technologies. All RNA constructs except for the m$^8$G4 and o$^8$G4 r(CpG)$_3$ oligos were synthesized by Dharmacon (a part of Horizon Discovery). Both the m$^8$G4 and o$^8$G4 r(CpG)$_3$ constructs were synthesized by the Yale School of Medicine oligo synthesis resource. All nucleic acid oligos in this study were resuspended to a stock concentration of 2 mM duplex and heat annealed prior to use at 95°C for 10 minutes, followed by slow cooling at room temperature for 30 min.

For NMR experiments, the constructs were dialyzed and buffer matched in 20 mM potassium phosphate (pH 6.4), 100 mM NaCl, 1 mM DTT, and 0.5 mM EDTA buffer to the $^{15}$N isotopically labeled protein constructs using 1 kDa cut-off mini dialysis kits (Cytiva). Oligo concentrations were measured using a NanoDrop 2000 Spectrophotometer (Thermo Scientific) and diluted to their final titration point concentrations of 125 µM for the free oligo (0:1), and 31.25 µM, 62.5 µM, 125 µM, 250 µM, or 500 µM for the 1:0.25, 1:0.5, 1:1, 1:2, and 1:4 titration points, respectively.

For Circular Dichroism (CD) measurements, 6-mer and 12-mer nucleic acid constructs were diluted from the 2 mM stocks to a final concentration of 50 µM and 24-mer nucleic acid constructs were diluted to a final concentration of 25 µM in 20 mM potassium phosphate (pH 6.4), 100 mM NaCl, 1 mM DTT, and 0.5 mM EDTA buffer.

For Isothermal Titration Calorimetry (ITC) measurements, all nucleic acid constructs were extensively dialyzed and buffer matched in 20 mM potassium phosphate (pH 6.4), 100 mM NaCl, and 0.5 mM EDTA buffer to the protein constructs using 1 kDa cut-off mini dialysis kits (Cytiva). Oligo concentrations were measured using a NanoDrop 2000 Spectrophotometer (Thermo Scientific) and diluted to a final concentration of 50 µM.

#### Domain Conservation Analysis

Z-Binding Domains (ZBDs) from human ADAR1 (*uniprot*: P55265), human ZBP1 (*uniprot:* Q9H171), goldfish PKZ (*uniprot*: Q7T2M9), vaccinia virus E3L (*uniprot*: P21605), and Cyprinid herpesvirus 1 ORF112 (*uniprot*: K7PC93) were aligned using the Clustal Omega Multiple Sequence Alignment tool [2] with default settings. The aligned sequences were then used to generate a consensus sequence using the EMBOSS Cons tool with default settings [2] and a conservation score using the

Protein Residue Conservation Prediction tool using the Property Entropy scoring method, a window size of 3, and the BLOSUM62 substitution matrix [3]. Alignments, conservation scores, and consensus sequences can be found in **Figure 1D.**

**Circular Dichroism and Absorbance Spectroscopy:**

All CD and absorbance measurements were collected simultaneously using a JASCO J-815 CD spectrophotometer (run using Spectra Manager version 2 (JASCO)) in a 0.1 cm quartz cuvette. CD and absorbance timecourses were measured by rapidly pipetting saturating amounts of each protein construct with the nucleic acid constructs and then measuring the ellipticity and absorbance at 264 nm for DNA constructs or 285 nm for RNA and LNA constructs. Saturating protein conditions were assumed to occur at a 1:(2n) molar ratio of nucleic acid:protein, where n is the number of ZBD binding sites on the duplex, similar to previous studies [4]. This molar ratio was not changed for the constructs with multiple ZBDs ($Z\alpha Z\beta^{ADAR1}$ and $Tandem^{ZBP1}$) despite having two times the number of ZBDs. The deadtime between the addition of protein and beginning the CD measurement was ~15 seconds. Temperatures were varied to try and match conversion rates between the faster DNA / modified RNA constructs (measured at 15°C) and the much slower RNA constructs (measured at 42°C), according to previous findings [4]. LNA constructs were similarly measured at 42°C, despite their inability to convert to the Z-form. Each timecourse was measured for a minimum of 1 hour if Z-conversion had reached equilibrium or for a maximum of 2 hours. Protein and nucleic acid stock solutions were preincubated separately at the measurement temperatures for at least 10 minutes before mixing. Full spectrum scans were performed directly after each time-course to validate the adoption of the Z-conformation. Circular dichroism and absorbance spectra were collected in 1-nm steps from 320 to 240 nm. Three scans were collected and averaged.

For many of our measured CD timecourses, especially as nucleic acid length was increased, pseudo-first order association constants failed to capture the experimental data. Therefore, the timecourse data was sequentially fit to one, two, or three rate exponential decay equations using the software package OriginLab. F-tests were performed between each increase in the number of exponential rates fitted to validate the need to move to more complex statistical models. Despite the apparent need for multiple rates, y-min constrained pseudo-first order plateaus were acquired by fitting the time-course data to a one phase association equation using the software package Prism to estimate the equilibrium population of Z-form nucleic acids as a percentage of the well-known $Z\alpha^{ADAR1}$ domain. Y-min constants were determined using the CD spectra of the free oligos to set a realistic starting ellipticity to account for measurement deadtime. Fitted multiexponential rates ($k$) and percent Z-form adoption can be found in **Supplementary Table 1** and **Table 2**, respectively.

**Isothermal Titration Calorimetry:**

Nucleic acid and protein constructs were buffer matched by dialyzing twice overnight in the same beaker at 4°C into 20 mM potassium phosphate (pH 6.4), 100 mM NaCl, and 0.5 mM EDTA using 1 kDa cut-off mini dialysis kits (Cytiva). The concentrations after dialysis were measured using a NanoDrop 2000 Spectrophotometer (Thermo Scientific). Protein stocks and nucleic acid stocks were diluted down to a final concentration of 1 mM and 50 µM, respectively. Binding heat for all 6-mer oligo constructs were measured on a Malvern ITC200 instrument (run using ITC 200 version 1.26.1 (Malvern)) at 35°C and a stirring speed of 750 rpm, with 120 s injection delays and a reference power of 9 $\mu cals^{-1}$. 12-mer $r(CpG)_6$ titrations were performed at both 35°C and 55°C with 120 s or 600 s injection delays, respectively. Otherwise, all other experimental conditions were the same as the 6-mer oligos. All titrations were measured with 37 consecutive 1 µL injections of 1 mM protein into 50 µM nucleic acid, with an initial injection of 0.4 µL.

**NMR Spectroscopy:**

All NMR samples were measured in 20 mM potassium phosphate (pH 6.4), 100 mM NaCl, 1 mM DTT, and 0.5 mM EDTA buffer at 35°C. All experiments were carried out on a Bruker 600 MHz spectrometer (run using TopSpin 4.2.0 (Bruker)) equipped with a 5/3 mm triple resonance $^{1}H/^{13}C/^{15}N/^{19}F$ cryoprobe (CP2.1 TCI). All of the $^{1}H$ carrier frequencies were centered on water.

*Relaxation Experiments*

For all relaxation experiments, $Z\alpha1^{ZBP1}$, $Z\alpha2^{ZBP1}$, and $Tandem^{ZBP1}$ were prepared at a final concentration of 500 µM in a total volume of 300 µL, supplemented with 5% $D_2O$, and measured in a 5 mm Shigemi tube (Wilmad Glass). $^{15}N$ $R_1$ relaxation experiments were collected with 2048 ($^{1}H$) × 100 ($^{15}N$) complex points, a recycle delay of 1 s, 32 scans, and spectral widths of 16 and 35 ppm for the $^{1}H$ and $^{15}N$ dimensions, respectively. The relaxation delays were 20, 60, 100, 200, 400, 600, 800, and 1200 ms. $^{15}N$ $R_{1\rho}$ relaxation experiments were run with 2048 ($^{1}H$) × 128 ($^{15}N$) complex points, a recycle delay of 1 s, 32 scans, and spectral widths of 16 and 35 ppm for the $^{1}H$ and $^{15}N$ dimensions, respectively. The relaxation delays under a spin-locking field strength of 1500 Hz were 12, 24, 48, 50, 72, 96, 120, 144, 165, 180, 210, and 250 ms. $^{15}N$ $R_2$ relaxation experiments were run with 2048 ($^1$H) × 128 ($^{15}$N) complex points, a recycle delay of 1 s,32 scans, and spectral widths of 16 and 35 ppm for the $^1$H and $^{15}$N dimensions, respectively. The relaxation delays were 16, 32, 64, 128, 160, 192, 224, and 256 ms. For the determination of effective overall tumbling times $\tau_C$, $R_2$ was calculated from $R_1$ and $R_{1\rho}$ to reduce increased relaxation contributions from the μs-ms time scale. R2 was calculated using the equation:

$$R_2 = R_{1\rho} + \left(R_{1\rho} - R1\right) tan^2(\theta) \tag{1}$$

where $\theta = \tan(\gamma_N B_1/2\pi\Delta v)$, $\Delta v$ is the resonance offset, $|\gamma_N B_1/2\pi|$ is the strength of the spin-lock field $B_1$, and $\gamma_N$ is the gyromagnetic ratio of the $^{15}$N spin. $\tau_C$ was then calculated from the $R_2/R_1$ ratio.

*$^{15}$N-HSQC and 1D Imino Nucleic Acid Titrations*

All $^{15}$N-HSQC spectra were measured with a final $^{15}$N-isotopically labeled protein concentration of 125 μM and a $D_2O$ concentration of 5% in a volume of 105 μL. Each titration point was made new with a constant protein concentration and incubated with the nucleic acid oligos for 30 min at 42°C immediately before being put into 3 mm NMR tubes (Norell) and measured. The $^{15}$N-HSQC spectra were acquired with a nonuniform sampling (NUS) scheme generated by the NUS@HMS scheme generator[5] employing 2048 complex data points in the direct dimension and 50% sampling of the original 400 complex points in the indirect $^{15}$N dimension. The spectral widths were 13.7 and 35 ppm for the $^1$H and $^{15}$N dimensions, respectively, with a recycle delay of 1 s and 16 scans. The $^{15}$N-HSQC assignments of $Z\alpha^{ADAR1}$, $Z\alpha1^{ZBP1}$, $Z\alpha1^{ZBP1}$, and $Tandem^{ZBP1}$ have been carried out previously [1, 6]. Chemical shift perturbations (CSPs) from the $^{15}$N-HSQC titrations of oligo into protein were calculated using the following equation [7, 8]:

$$\text{CSP} = \sqrt{\left(\delta_{H,free} - \delta_{H,bound}\right)^2 + 0.2\left(\delta_{N,free} - \delta_{N,bound}\right)^2} \tag{2}$$

where $\delta_H$ and $\delta_N$ are the chemical shifts of a peak in the $^1$H and $^{15}$N dimensions, respectively.

Immediately after measurement of each $^{15}$N-HSQC spectra, 1D $^1$H spectra were measured for analysis of the imino protons of the nucleic acids. The 1D $^1$H spectra were acquired with 256 scans. A-, B-, and Z-form imino proton peak positions have been previously assigned for the $d(CpG)_3$, $r(CpG)_3$, and LNA $(CpG)_3$ oligos ([9, 10]), and inferred for the $m^8$G4 $r(CpG)_3$ oligo based on its resemblance to $Z\alpha^{ADAR1}$-induced Z-form $r(CpG)_3$.

### SUPPLEMENTAL FIGURES

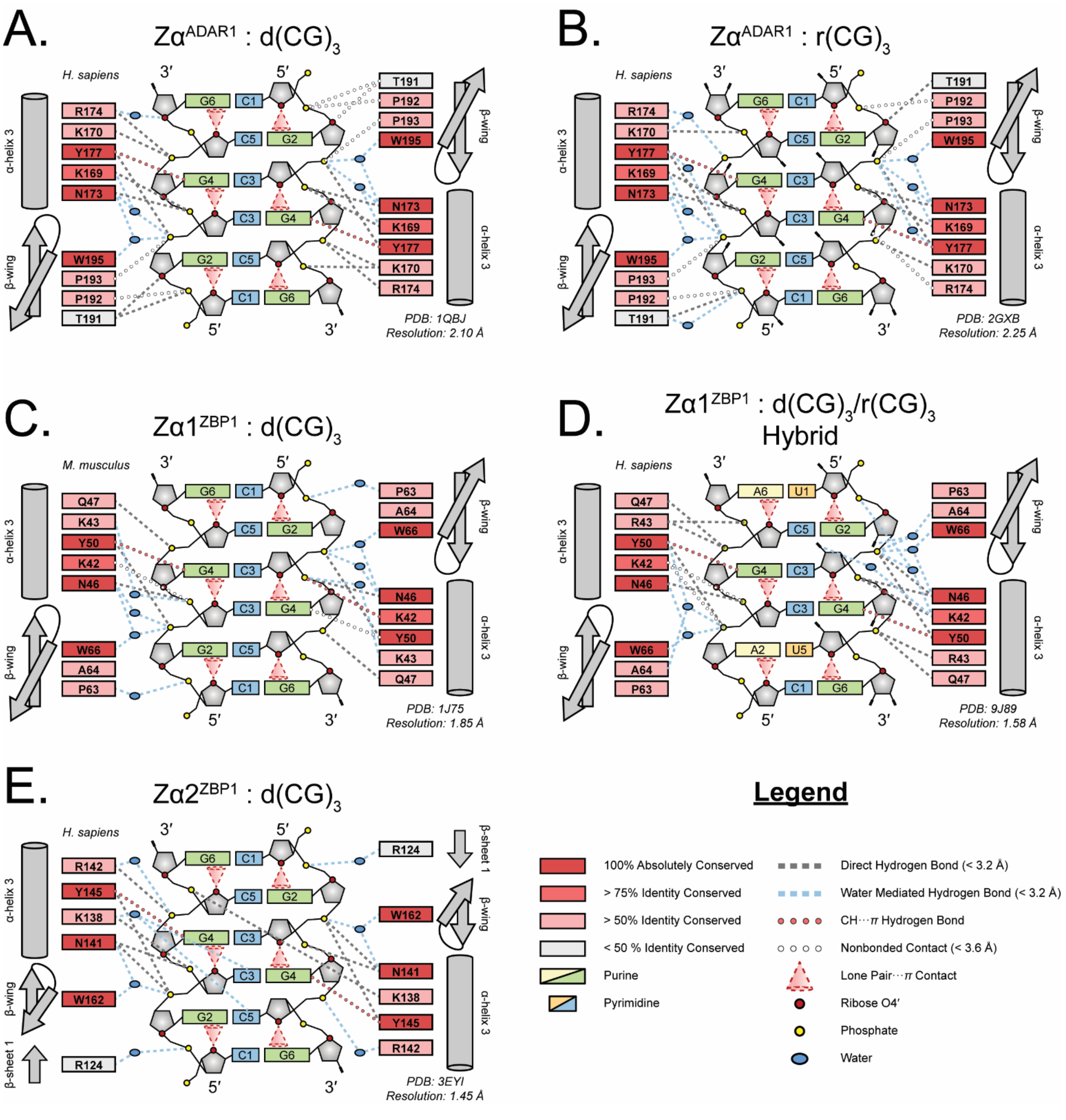


**Supplementary Figure 1.** Binding modes of mammalian Zα domains with DNA and RNA generated by NUCPLOT [11]. Moderately stringent optimal hydrogen bond distances (<3.2 Å), hydrogen bond geometries, and van der Waals contacts (<3.6 Å) were chosen because of the high resolution (<2.5 Å) structures available. **(A, B)** Binding mode of the $Z\alpha^{ADAR1}$ domain with self-complementary $d(CG)_3$ and $r(CG)_3$ oligos as seen in the crystal structures 1QBJ and 2GXB, respectively [12, 13]. **(C, D).** Binding mode of $Z\alpha1^{ZBP1}$ with a self-complementary $d(CG)_3$ and a hybrid $d(CG)_3$:$r(CG)_3$ oligo as seen in the crystal structures 1J75 and 9J89, respectively [14, 15]. It is important to note that our analysis of the crystal structures differ from those reported by the authors in [15], likely due to different restraints used in the analysis of bonds/contacts. **(E).** Binding mode of $Z\alpha2^{ZBP1}$ with a self-complementary $d(CG)_3$ oligo as seen in the crystal structure 3EYI [16].

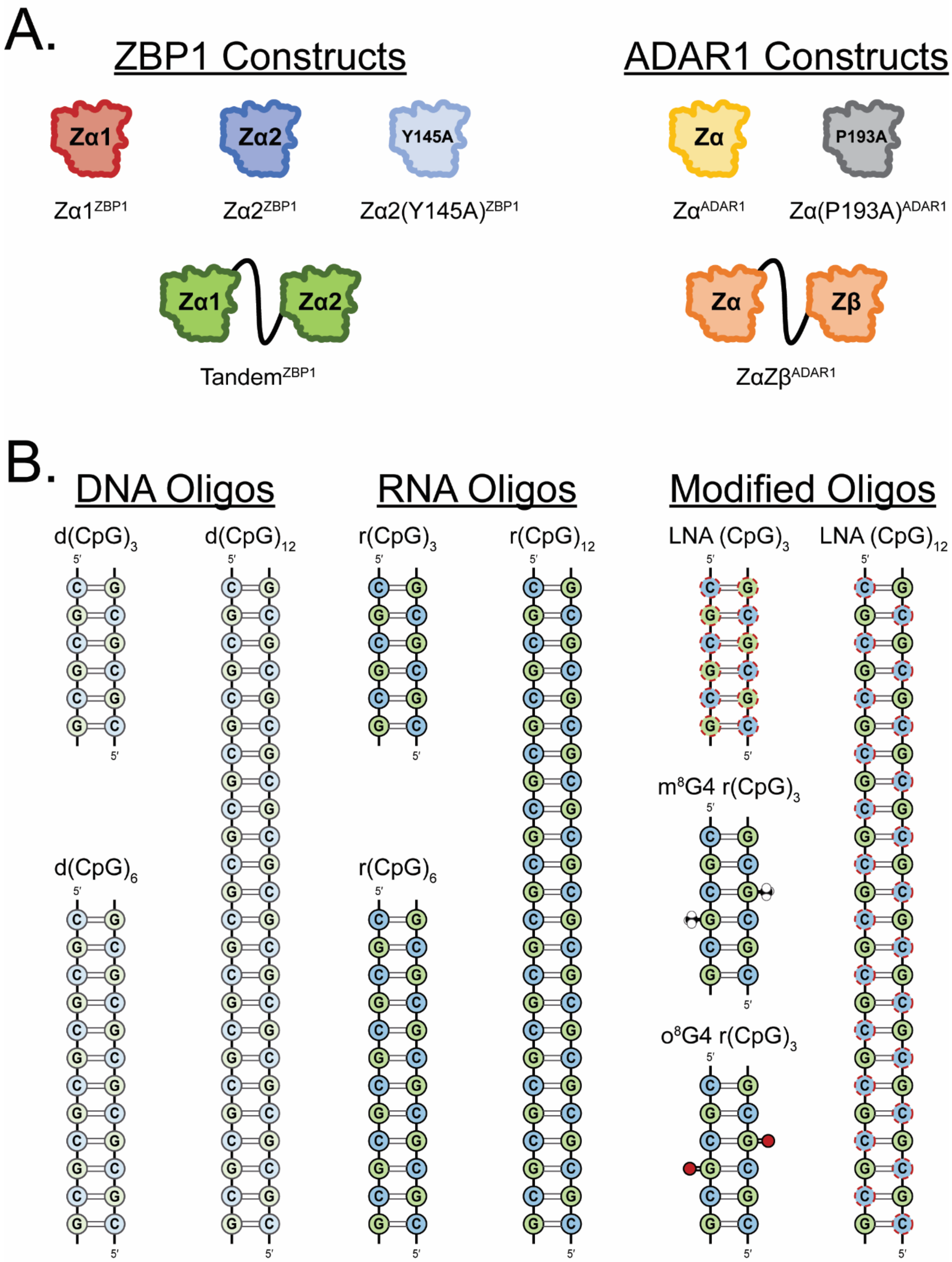


**Supplementary Figure 2.** Protein and oligo constructs. **(A)** ZBP1 and ADAR1 protein constructs and the names referenced throughout the manuscript. ZBP1 constructs include the isolated $Z\alpha1^{ZBP1}$ and $Z\alpha2^{ZBP1}$ domains, $Y145A^{ZBP1}$ in which the critical tyrosine residue is mutated and renders the domain nonfunctional, and $Tandem^{ZBP1}$ construct which contains both Zα1, Zα2, and the interconnected WT linker. ADAR1 constructs include the isolated $Z\alpha^{ADAR1}$ domain, $P193A^{ADAR1}$ in which one of the residues in the *cis-trans* proline-proline motif is mutated, and the tandem $Z\alpha Z\beta^{ADAR1}$ construct. Color-schemes for the proteins remain consistent throughout the manuscript. **(B)** Nucleic acid oligo constructs and the names referenced throughout the manuscript. All oligos are self-complimentary and were annealed immediately prior to measurements. Locked nucleic acid (LNA) sugar moieties are represented as red dashed circles. 8-methylguanine ($m^8G$) modifications are represented as black and white methyl space-filled models and 8-oxoguanine ($o^8G$) modifications are represented as red circles.

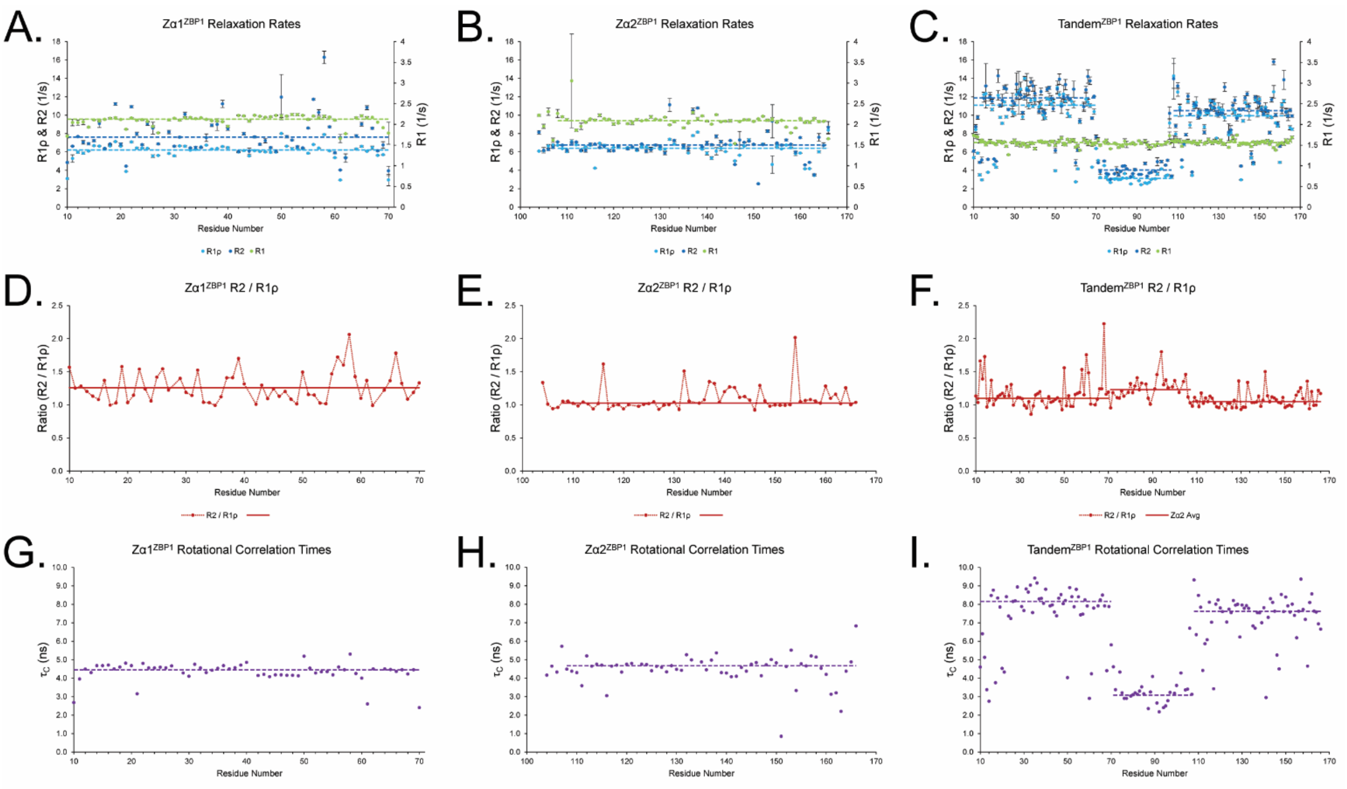


**Supplementary Figure 3.** Measured relaxation rates and calculated relaxation parameters of ZBP1 constructs. **(A-C)** Overlayed R2, R1ρ, and R1 relaxation rates for each residue within the $Z\alpha1^{ZBP1}$, $Z\alpha2^{ZBP1}$, and $Tandem^{ZBP1}$ constructs, respectively. Average R2, R1ρ, and R1 rates reflect small, well-folded domains for the individual $Z\alpha1^{ZBP1}$ and $Z\alpha2^{ZBP1}$. The low R2 and R1ρ rates for the intrinsically disordered linker, as well as the larger rates for the ZBDs in the larger $Tandem^{ZBP1}$ construct reflect a tethered domain system with minimal interaction between ZBDs, and match previous reports [1]. **(D-F)** Ratio of R2 / R1ρ can be used to inform on regions experiencing increased R2 rates as a result of domain motions and/or exchange processes. $Z\alpha2^{ZBP1}$ has a region of increased $R2_{Ex}$ between residues 137-142, likely as a result of the dynamic folding/unfolding of a $3_{10}$ helix. Interestingly, the IDR linker of $Tandem^{ZBP1}$ also experiences increased R2 / R1ρ ratios, indicating contributions from exchange processes. **(G-I)** Calculated rotational correlation times ($\tau_C$) of each ZBP1 construct. $Z\alpha1^{ZBP1}$ and $Z\alpha2^{ZBP1}$ $\tau_C$ averages (4.45 ± 0.07 ns and 4.66 ± 0.11 ns, respectively) reflect small, well-folded domains. Similarly, $\tau_C$ domain averages of $Tandem^{ZBP1}$ reflect a tethered domain system.

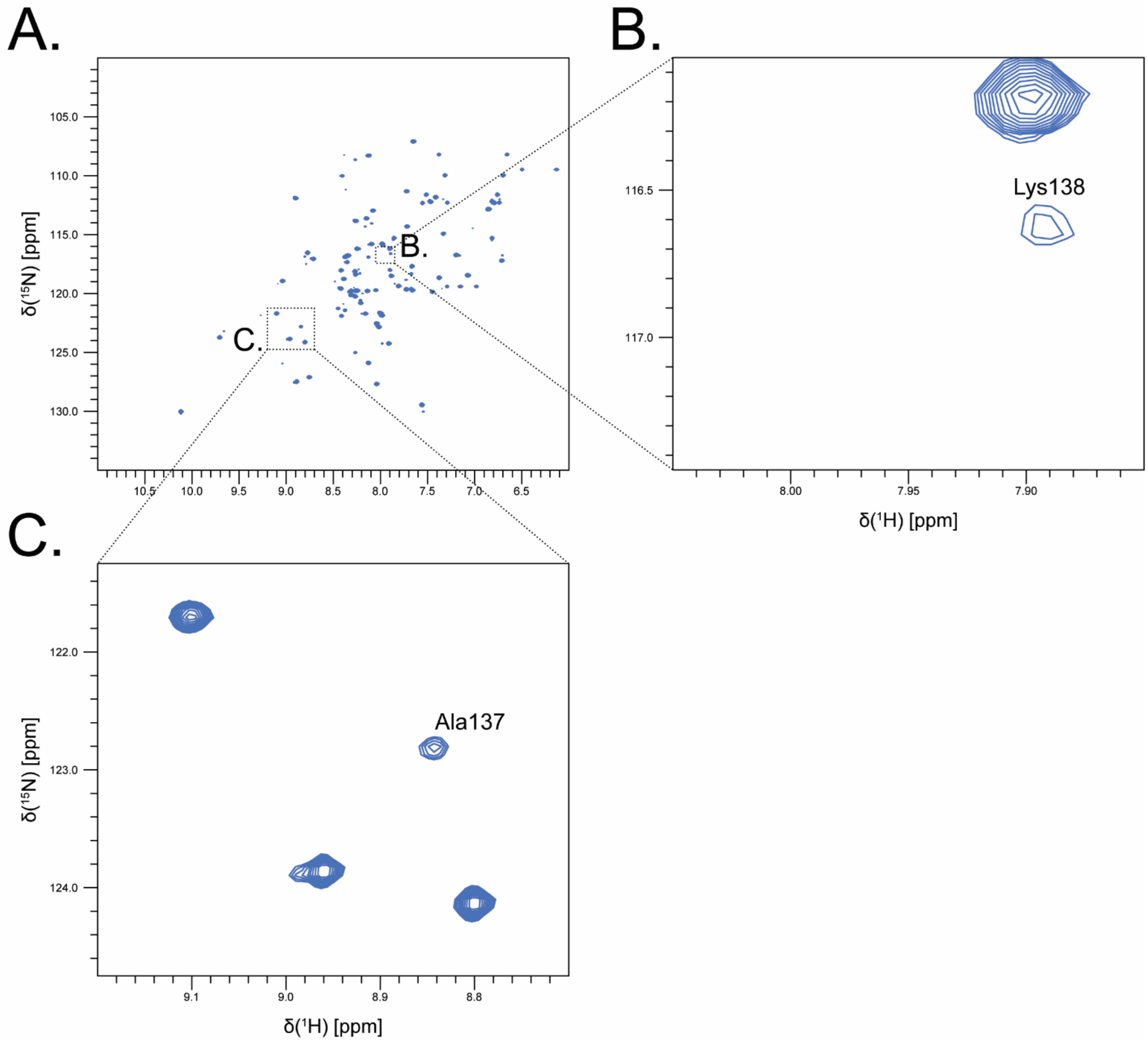


**Supplementary Figure 4.** $^{15}$N-HSQC of Zα2$^{ZBP1}$. **(A)** Full Spectrum $^{15}$N-HSQC of Zα2 shows many dispersed peaks, indicative of a well-folded protein. **(B-C)** Zoomed in regions surrounding Lys138 and Ala137, respectively. Both residues are heavily broadened compared to other peaks in the spectrum, indicating the presence of μs-ms exchange contributions.

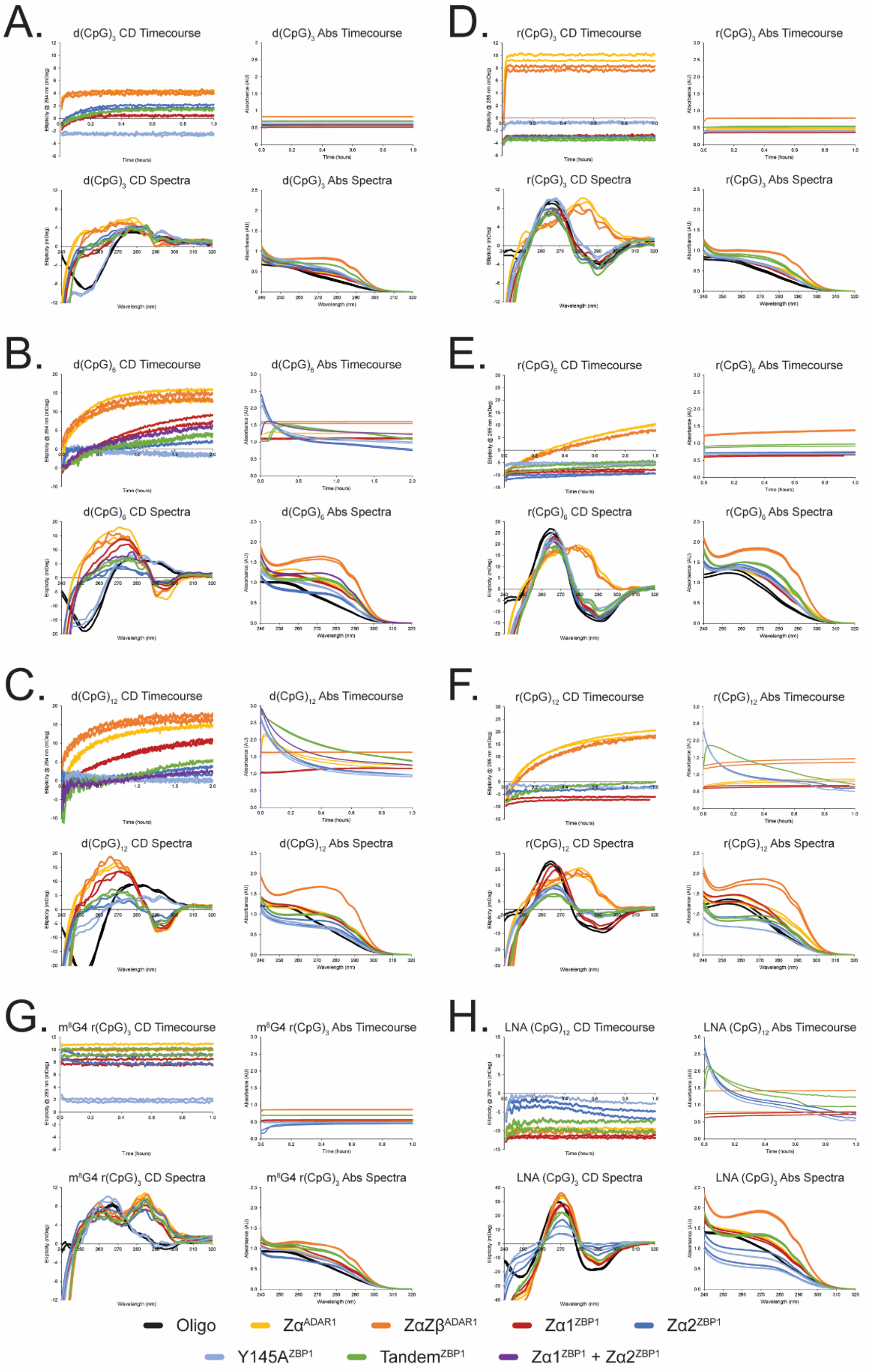


**Supplementary Figure 5.** Concurrent circular dichroism and absorbance measurements for each oligo tested in **(A-H)** under saturating protein concentrations. CD and absorbance timecourse measurements were taken simultaneously and reflect similar processes. CD and absorbance spectra were taken immediately after the timecourses. All plots represent the raw data collected except for the CD timecourses which instead depict the moving average (period of 20) of the data. **(A-C)** DNA oligos were measured at 15°C for at least one hour or until equilibrium was reached. **(D-F)** RNA oligos were measured at 42°C for at least 1 hour to roughly match the rate of the DNA oligos at 15°C according to previous findings [4]. **(G)** m$^8$G4 r(CpG) oligos were measured at 15°C in order to slow down the conversion rate. **(H)** LNA oligos were measured at 42°C.

**Supplementary Table 1. Multi-Rate Exponential Fits of ZBD Constructs Extracted from Circular Dichroism Timecourses**

| | Oligo | # Fits | Amplitude 1 | Amplitude 2 | Amplitude 3 | Rate 1 (hr$^{-1}$) | Rate 2 (hr$^{-1}$) | Rate 3 (hr$^{-1}$) |
|---|---|---|---|---|---|---|---|---|
| $Z\alpha^{ADAR1}$ | d(CG)3 | Two-Rate | -2.40 ± 0.24 | -0.72 ± 0.13 | | 122.97 ± 12.36 | 11.18 ± 1.22 | |
| | d(CG)6 | Three-Rate | -12.87 ± 0.45 | -2.51 ± 0.65 | -2.51 ± 0.63 | 2.67 ± 0.05 | 10.17 ± 1.60 | 47.85 ± 9.69 |
| | d(CG)12 | Two-Rate | -9.77 ± 0.67 | -7.22 ± 0.75 | | 1.51 ± 0.10 | 7.18 ± 0.57 | |
| | r(CG)3 | One-Rate | -12.90 ± 0.22 | | | 96.97 ± 1.36 | | |
| | r(CG)6 | Three-Rate | -25.17 ± 0.24 | -3.82 ± 0.37 | -0.98 ± 0.24 | 0.91 ± 0.02 | 4.97 ± 0.41 | 56.94 ± 12.03 |
| | r(CG)12 | Three-Rate | -12.15 ± 1.22 | -12.24 ± 0.71 | -6.83 ± 0.90 | 1.64 ± 0.07 | 5.47 ± 0.47 | 21.05 ± 1.51 |
| | LNA (CG)12 | One-Rate | -3.08 ± 0.28 | | | 104.26 ± 13.23 | | |
| $Z\alpha Z\beta^{ADAR1}$ | d(CG)3 | Two-Rate | -2.82 ± 0.35 | -0.56 ± 0.16 | | 109.15 ± 14.08 | 8.69 ± 1.94 | |
| | d(CG)6 | Two-Rate | -10.83 ± 0.23 | -2.70 ± 0.43 | | 2.17 ± 0.05 | 23.31 ± 3.96 | |
| | d(CG)12 | Three-Rate | -9.44 ± 0.56 | -2.60 ± 0.65 | -2.81 ± 1.18 | 2.25 ± 0.10 | 13.02 ± 3.89 | 163.57 ± 69.79 |
| | r(CG)3 | One-Rate | -11.77 ± 0.31 | | | 91.49 ± 1.99 | | |
| | r(CG)6 | Two-Rate | -24.69 ± 0.20 | -2.99 ± 0.36 | | 0.85 ± 0.02 | 7.60 ± 0.93 | |
| | r(CG)12 | Two-Rate | -19.00 ± 0.47 | -10.58 ± 0.51 | | 1.82 ± 0.04 | 9.53 ± 0.42 | |
| | LNA (CG)12 | One-Rate | -3.03 ± 0.59 | | | 94.38 ± 25.59 | | |
| $Z\alpha 1^{ZBP1}$ | d(CG)3 | One-Rate | -1.85 ± 0.08 | | | 11.58 ± 0.41 | | |
| | d(CG)6 | Two-Rate | -14.99 ± 0.12 | -0.97 ± 0.20 | | 1.06 ± 0.02 | 8.94 ± 1.71 | |
| | d(CG)12 | Two-Rate | -13.15 ± 0.13 | -1.68 ± 0.25 | | 0.99 ± 0.02 | 15.69 ± 2.58 | |
| | r(CG)3 | One-Rate | -1.04 ± 0.13 | | | 58.96 ± 5.58 | | |
| | r(CG)6 | Two-Rate | -1.26 ± 0.24 | -0.99 ± 0.09 | | 85.72 ± 15.78 | 6.08 ± 0.52 | |
| | r(CG)12 | Two-Rate | -1.00 ± 0.27 | -1.54 ± 0.11 | | 93.98 ± 25.33 | 6.99 ± 0.42 | |
| | LNA (CG)12 | One-Rate | -2.21 ± 0.34 | | | 96.79 ± 11.97 | | |
| $Z\alpha 2^{ZBP1}$ | d(CG)3 | Two-Rate | -2.54 ± 0.14 | -0.46 ± 0.22 | | 8.59 ± 0.32 | 71.75 ± 34.92 | |
| | d(CG)6 | One-Rate | -4.66 ± 0.38 | | | 0.62 ± 0.07 | | |
| | d(CG)12 | | | | | | | |
| | r(CG)3 | One-Rate | -1.02 ± 0.15 | | | 65.84 ± 7.86 | | |
| | r(CG)6 | Two-Rate | -2.38 ± 0.29 | -1.74 ± 0.31 | | 0.96 ± 0.13 | 132.62 ± 19.76 | |
| | r(CG)12 | Two-Rate | -0.99 ± 0.58 | 0.01 ± 0.04 | | 3.12 ± 2.11 | 4.52 ± 1.84 | |
| | LNA (CG)12 | Two-Rate | -5.86 ± 0.54 | 21.50 ± 21.59 | | 46.08 ± 3.95 | 0.21 ± 0.17 | |
| $Y145A^{ZBP1}$ | d(CG)3 | | | | | | | |
| | d(CG)6 | Two-Rate | -5.56 ± 0.34 | 2.28 ± 0.13 | | 23.70 ± 1.48 | 1.11 ± 0.14 | |
| | d(CG)12 | Two-Rate | -9.16 ± 0.45 | 2.76 ± 0.26 | | 23.62 ± 1.28 | 2.30 ± 0.21 | |
| | r(CG)3 | One-Rate | -1.21 ± 0.15 | | | 58.11 ± 6.11 | | |
| | r(CG)6 | One-Rate | -0.84 ± 0.21 | | | 52.71 ± 10.49 | | |
| | r(CG)12 | | | | | | | |
| | LNA (CG)12 | Two-Rate | -8.16 ± 0.37 | 7.93 ± 5.32 | | 44.61 ± 3.66 | 0.42 ± 0.36 | |
| $Tandem^{ZBP1}$ | d(CG)3 | Two-Rate | -2.45 ± 0.09 | -0.86 ± 0.35 | | 5.82 ± 0.21 | 175.84 ± 68.18 | |
| | d(CG)6 | One-Rate | -8.81 ± 0.14 | | | 1.37 ± 0.04 | | |
| | d(CG)12 | One-Rate | -12.62 ± 0.68 | | | 0.64 ± 0.05 | | |
| | r(CG)3 | One-Rate | -0.94 ± 0.19 | | | 89.80 ± 14.93 | | |
| | r(CG)6 | Two-Rate | -4.79 ± 0.14 | -1.14 ± 0.20 | | 0.94 ± 0.04 | 34.41 ± 5.47 | |
| | r(CG)12 | Two-Rate | -5.13 ± 0.38 | -3.83 ± 0.75 | | 1.79 ± 0.61 | 11.38 ± 2.59 | |
| | LNA (CG)12 | One-Rate | -2.55 ± 0.45 | | | 27.41 ± 4.05 | | |

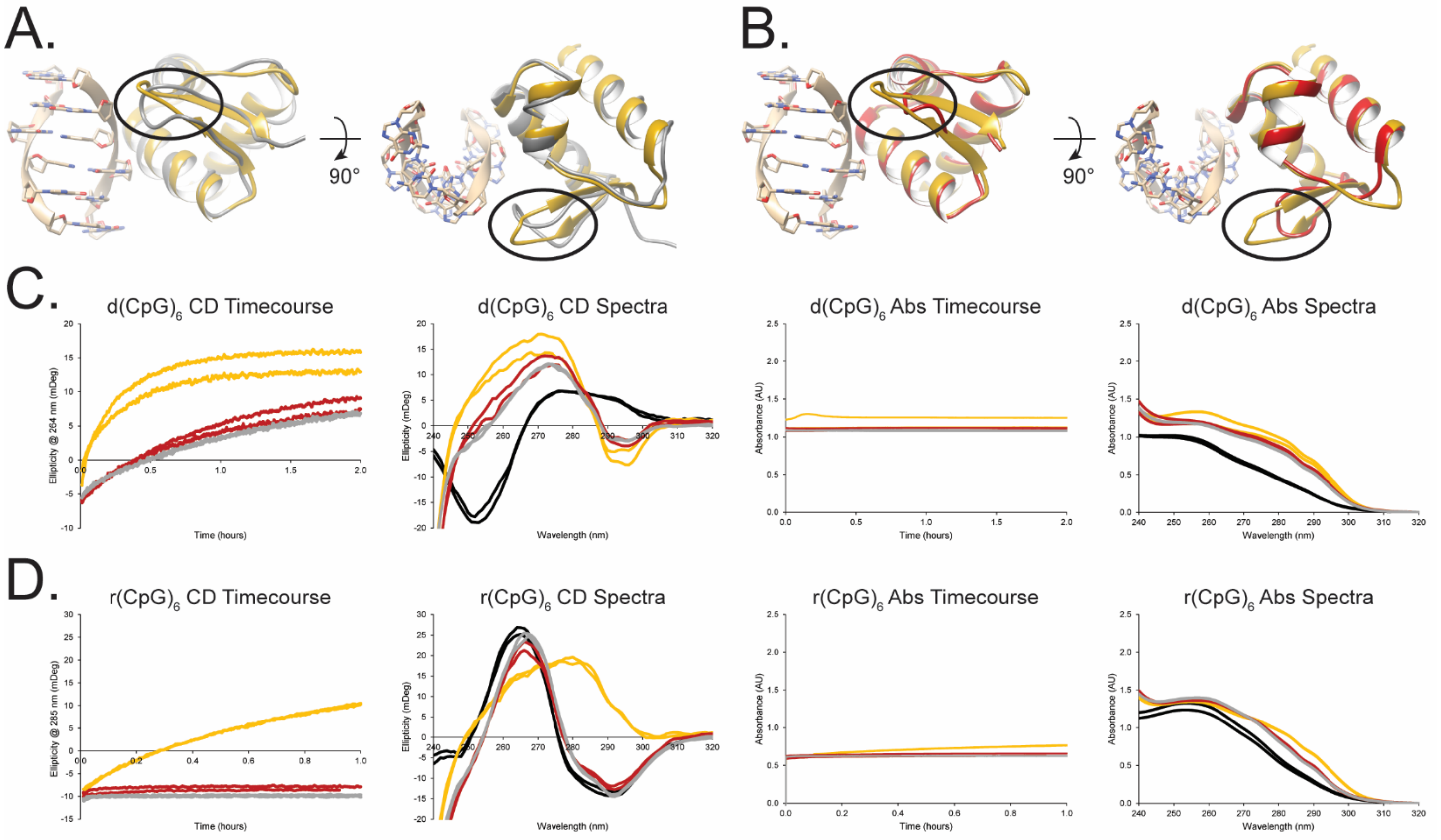


**Supplementary Figure 6.** *Cis-trans* proline-proline motif is critical for the β-wing fold and A-to-Z conversion activity. **(A)** Superposition of the crystal structure of ADAR1's Zα domain in complex with a $d(CpG)_3$ oligo (*PDB*: 1QBJ) with the NMR solution structure of $P193A^{ADAR1}$ (*PDB*: 8GBD) showcasing the misfolding of the β-wing (black circles) caused by the P193A point mutation. **(B)** Superposition of the same crystal structure (*PDB*: 1QBJ) with the crystal structure of the Zα1 domain of ZBP1 (*PDB*: 1J75) showcasing the different fold of the shortened β-wing (black circles). **(C-D)** CD and absorbance measurements of $Z\alpha^{ADAR1}$ (yellow), $Z\alpha1^{ZBP1}$ (red), and $P193A^{ADAR1}$ (grey) with $d(CpG)_6$ and $r(CpG)_6$ oligos (C and D, respectively). Experimental conditions were the same as in Supplementary Figure 5 and black lines represent the spectra of the free oligos.

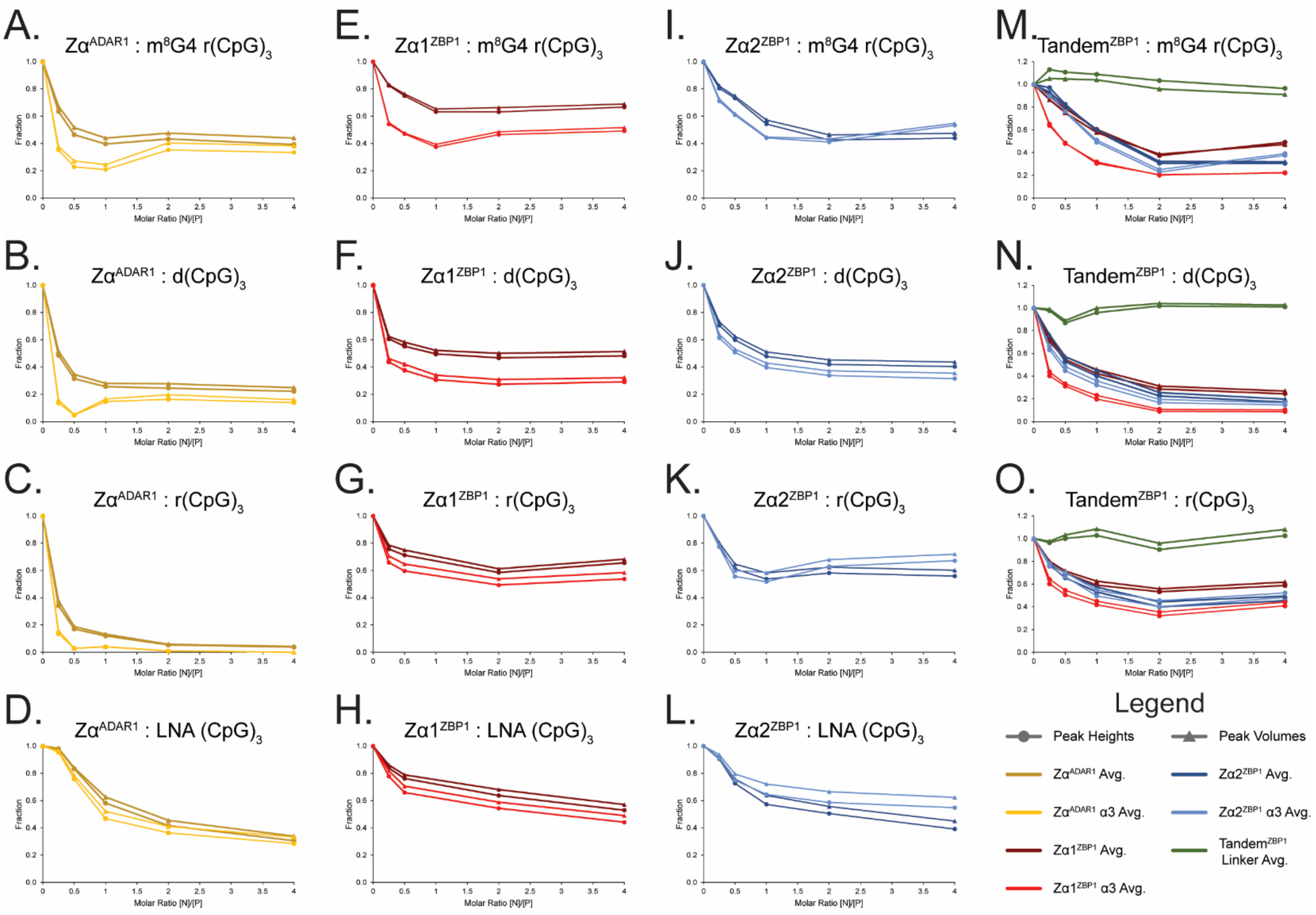


**Supplementary Figure 7.** Average peak heights and volumes for amide signals in each ZBD – oligo $^{15}$N-HSQC titration series represented by circles and triangles, respectively. **(A-D)** Average peak heights and volumes of each Zα$^{ADAR1}$ titration series across the entire domain (gold) and across just the α3-helix (yellow). **(E-H)** Average peak heights and volumes of each Zα1$^{ZBP1}$ titration series across the entire domain (maroon) and across just the α3-helix (red). **(I-L)** Average peak heights and volumes of each Zα2$^{ZBP1}$ titration series across the entire domain (navy) and across just the α3-helix (blue). **(M-O)** Average peak heights and volumes of each Tandem$^{ZBP1}$ titration series across the entire Zα1 domain (maroon) or across Zα1's α3-helix (red), across the entire IDR linker (green), and across the entire Zα2 domain (navy) or across Zα2's α3-helix (blue).

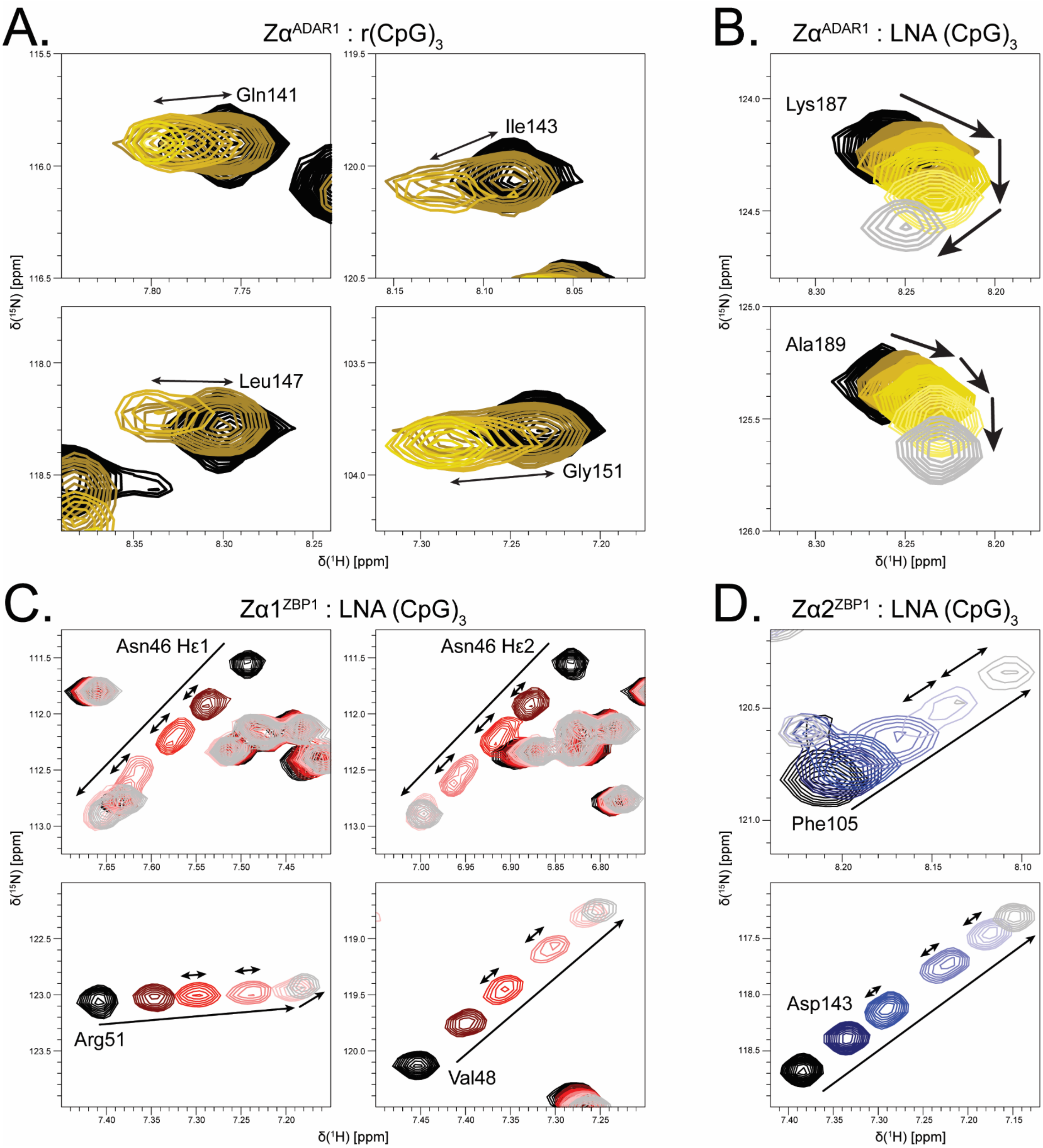


**Supplementary Figure 8**. Atypical shift behaviors identified in the ADAR1 and ZBP1 oligo titrations. **(A)** Characteristic examples of slow exchange in the Zα$^{ADAR1}$ – r(CpG)$_3$ $^{15}$N-HSQC titration series. The amide signal of free protein (black) slowly disappears, and the amide signal of bound protein (shades of yellow and grey) slowly appears as oligo concentrations are increased. **(B)** The Zα$^{ADAR1}$ – LNA $^{15}$N-HSQC titration series show nonlinear, multimodal binding modes consistent with multiple oligos binding the same ZBD [9]. **(C, D)** Amide signals in the interaction interface of Zα1$^{ZBP1}$ (red, **C**) and Zα2$^{ZBP1}$ (blue, **D**) show both fast and slow exchange processes at intermediate titration points in the LNA (CpG)$_3$ titration series.

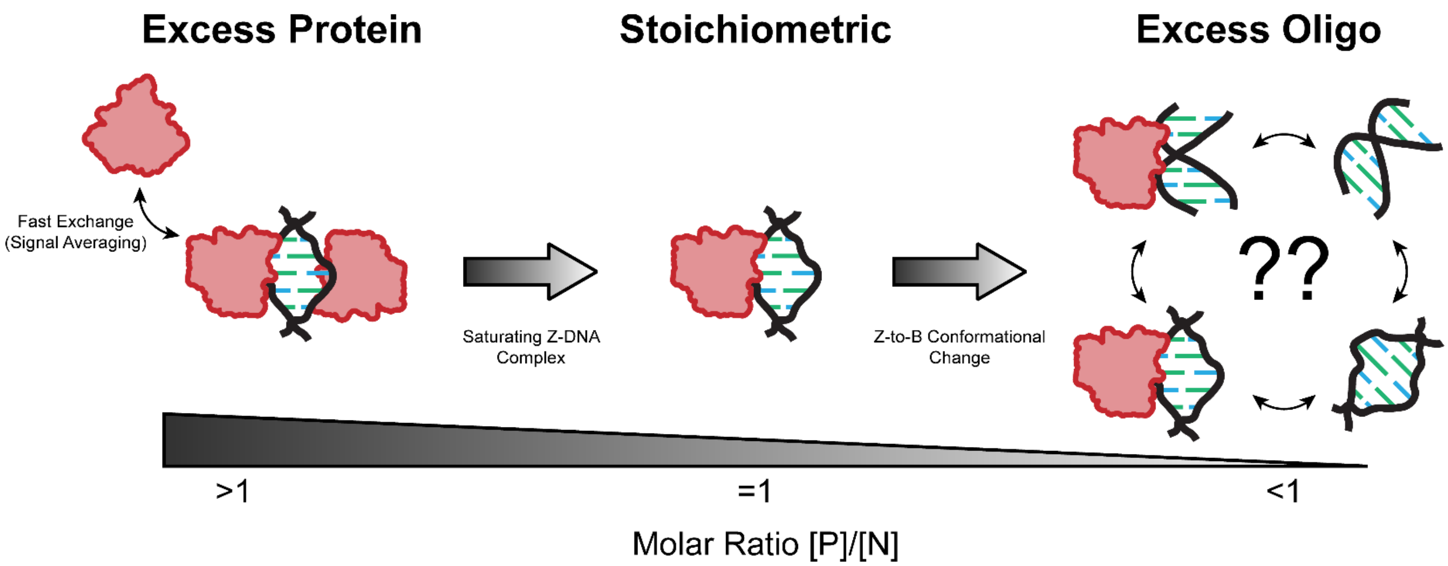


**Supplementary Figure 9**. Graphical model explaining bimodal binding shifts seen in ZBP1 – d(CpG)$_3$ oligo titrations. Under protein saturating conditions, ZBP1 ZBDs are able to fully saturate the B-DNA oligos and convert them to the Z-conformation. Fast/intermediate exchange between free and bound ZBDs reduce the overall magnitude of shifts at early titration points. As nucleic acid concentrations increase up to equimolar stoichiometries, ZBD – Z-DNA complexes are saturated, and first binding mode reaches maximal shifts. Because ZBP1's ZBDs form unstable complexes with short 6-mer DNAs, as oligo concentrations are further increased, ZBDs begin to be titrated off and lose the ability to stabilize the Z-conformation, resulting in a switch to the second, right-handed binding mode. It is currently unclear whether the oligos revert on the ZBD or if dissociation from the ZBD results in a reversion to the B-form.

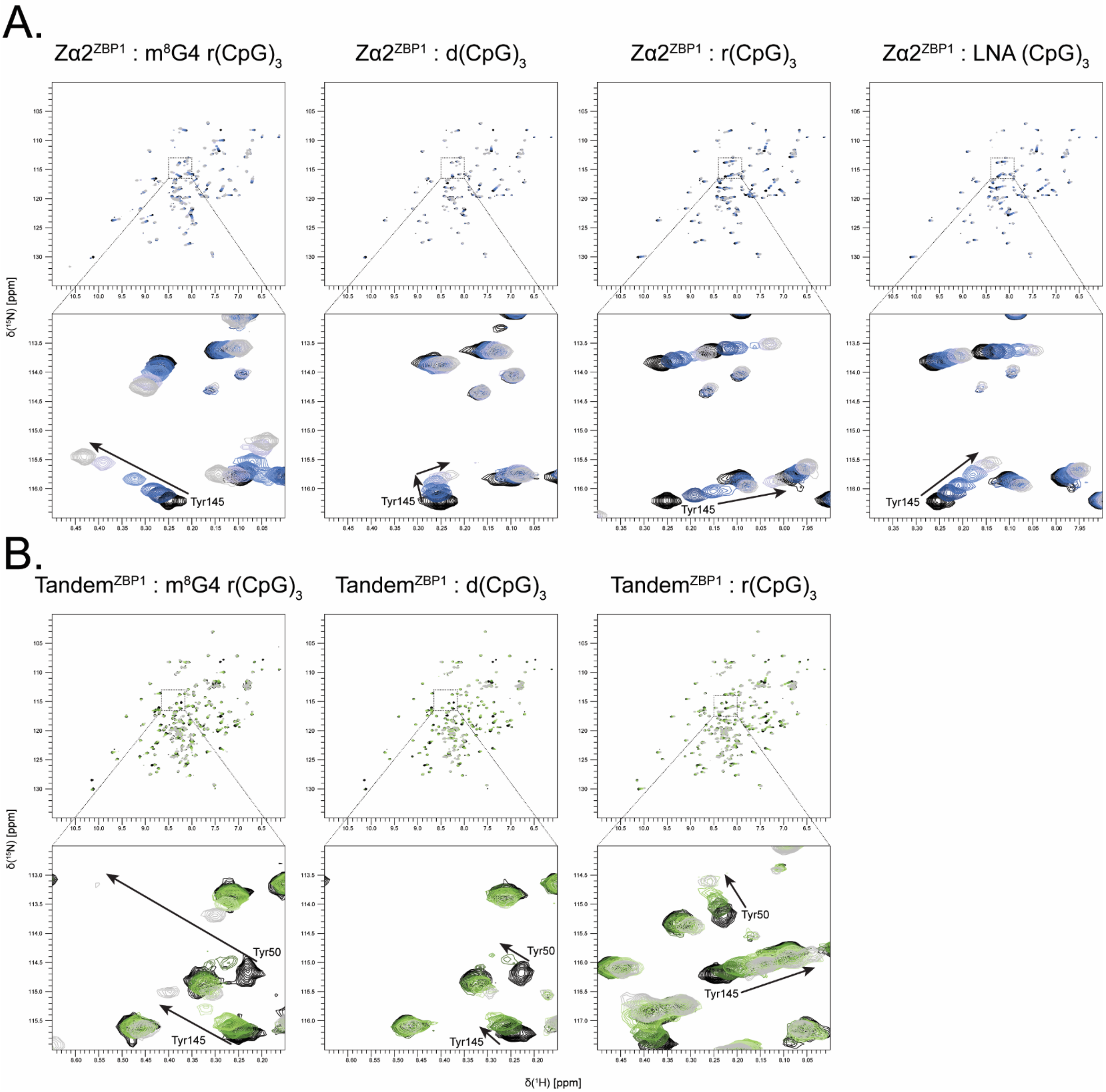


**Supplementary Figure 10.** $^{15}$N-HSQC titration series of Zα2$^{ZBP1}$ and Tandem$^{ZBP1}$ with different 6-mer DNA/RNA oligos measured at 35°C. **(A)** Titration series of Zα2$^{ZBP1}$ with the Z-form m$^8$G4 r(CpG)$_3$, B-form d(CpG)$_3$, A-form r(CpG)$_3$, and A-Like LNA (CpG)$_3$. Titration points are color coded from black (Apo), through shades of blue (1:0.25, 1:0.50, 1:1, 1:2), to grey (1:4). Inset boxes highlight residues Y145 which show significant peak shifts and are critical for flipping abilities. Zα2$^{ZBP1}$ – d(CpG)$_3$ titration series has bimodal binding modes similar to Zα1$^{ZBP1}$. **(B)** Titration series of Tandem$^{ZBP1}$ with the same oligos seen in (A) with the exception of the LNA (CpG)$_3$. Titration points are color coded from black (Apo), through shades of green (1:0.25, 1:0.50, 1:1, 1:2), to grey (1:4). Inset boxes highlight the CSPs of the critical Y50 and Y145 residues of Zα1 and Zα2, respectively. Peaks for Zα1 and Zα2 shift to similar positions as seen with the Zα1$^{ZBP1}$ and Zα2$^{ZBP1}$ titration series but are heavily broadened due to increased complex size

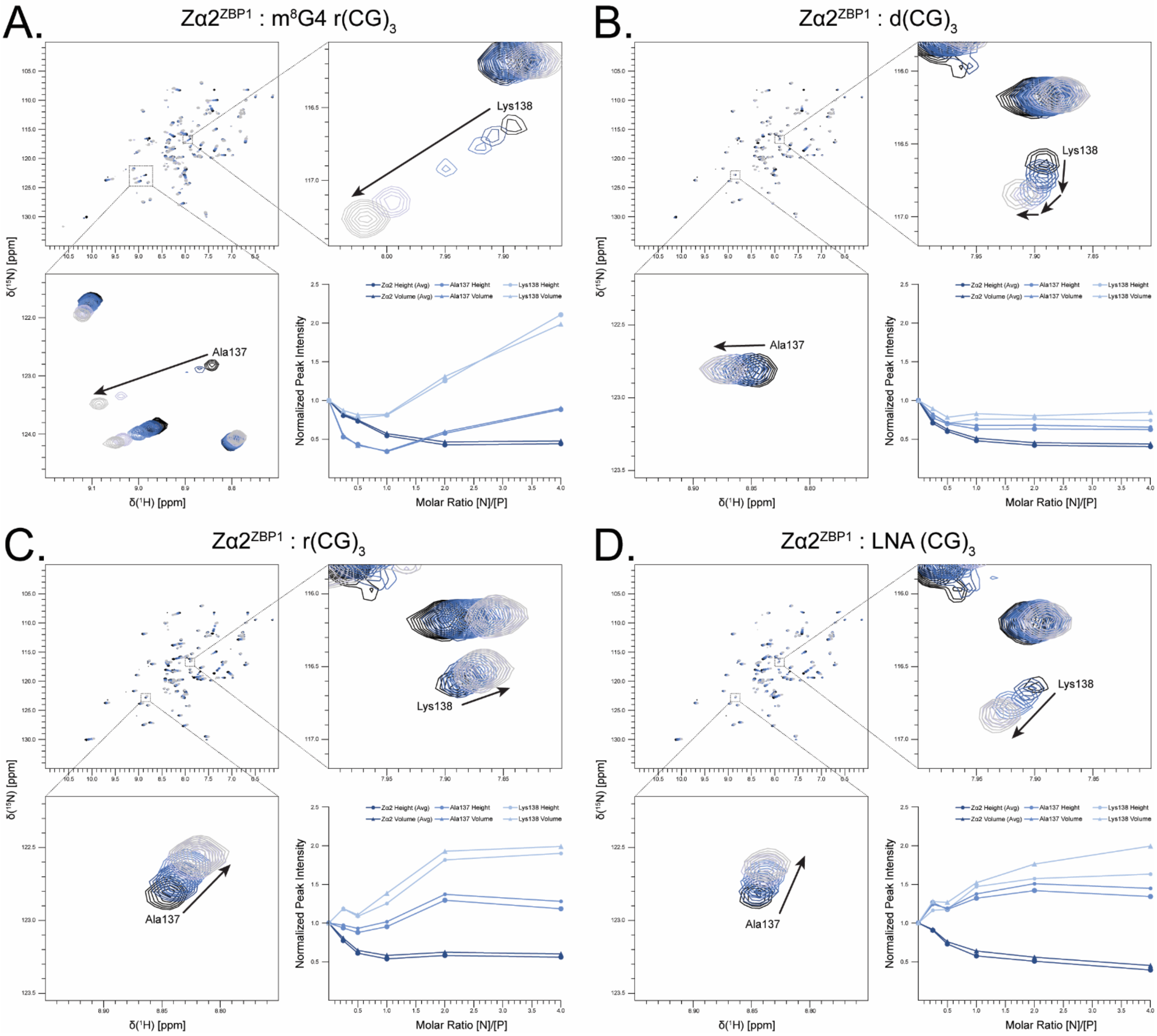


**Supplementary Figure 11.** Intermediate exchange seen in residues within the $3_{10}$-helix at the N-terminus of Zα2[ZBP1]'s recognition helix is quenched upon addition of increasing concentrations of m[8]G4 r(CpG)$_3$ **(A)**, r(CpG)$_3$ **(C)**, and LNA (CpG)$_3$ **(D)** oligos. Titration of d(CpG)$_3$ **(B)** oligo into Zα2[ZBP1] does not quench intermediate exchange, but displays unique, multimodal binding modes for Lys138.

**Supplementary Figure 12.** Raw Isothermal Titration Calorimetry measurements of Zα^ADAR1 **(A-C)**, ZαZβ^ADAR1 **(D-F)**, Zα1^ZBP1 **(G-I)**, Zα2^ZBP1 **(J-L)**, and Tandem^ZBP1 **(M-O)** into m⁸G4 r(CpG)$_3$, d(CpG)$_3$, and r(CpG)$_3$ oligos, respectively. Y145A^ZBP1 **(P,Q)** was titrated into d(CpG)$_3$ and r(CpG)$_3$ oligos, respectively. All measurements were done in duplicate and measured at 35°C with 2 min injection intervals.

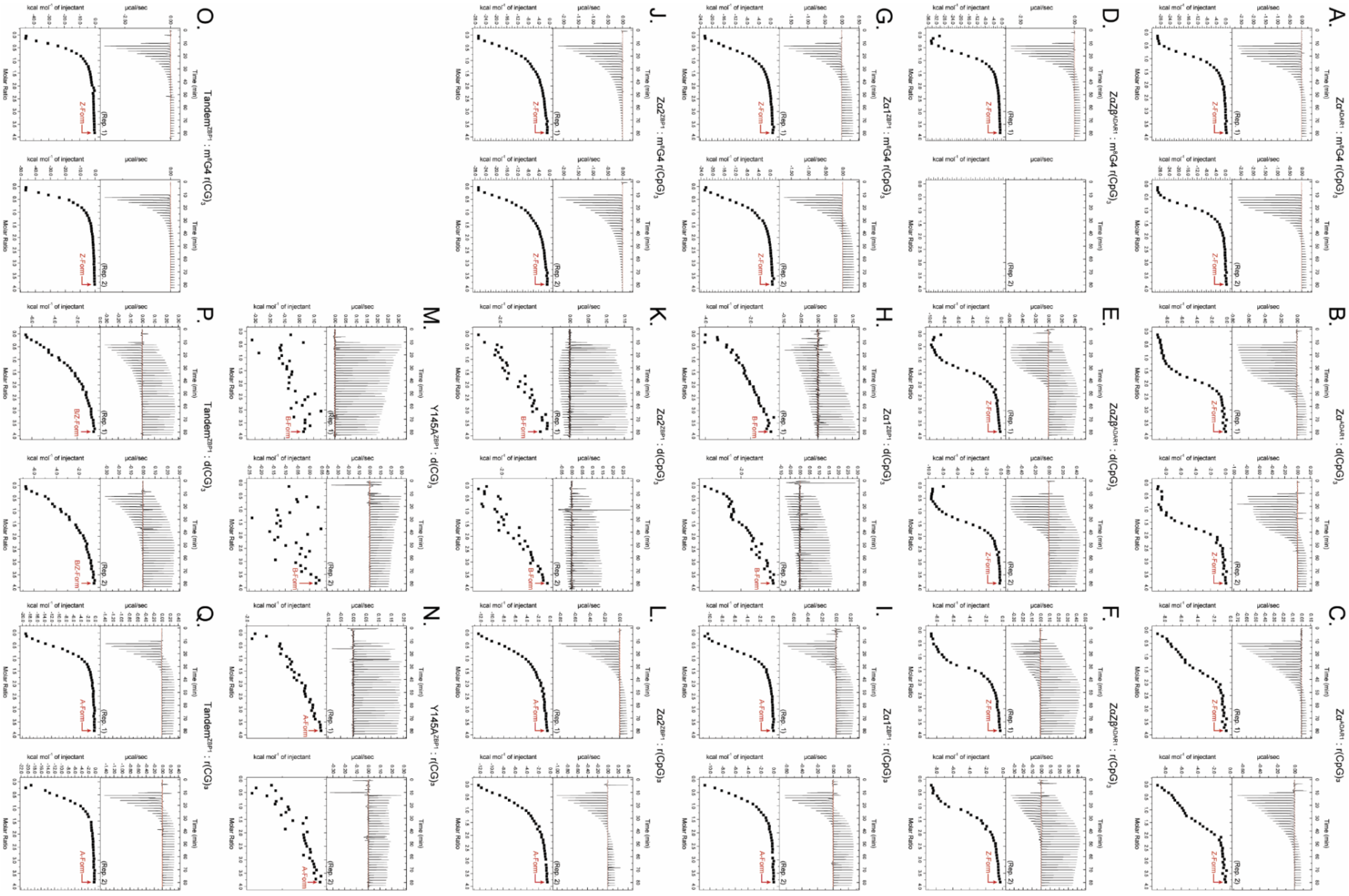

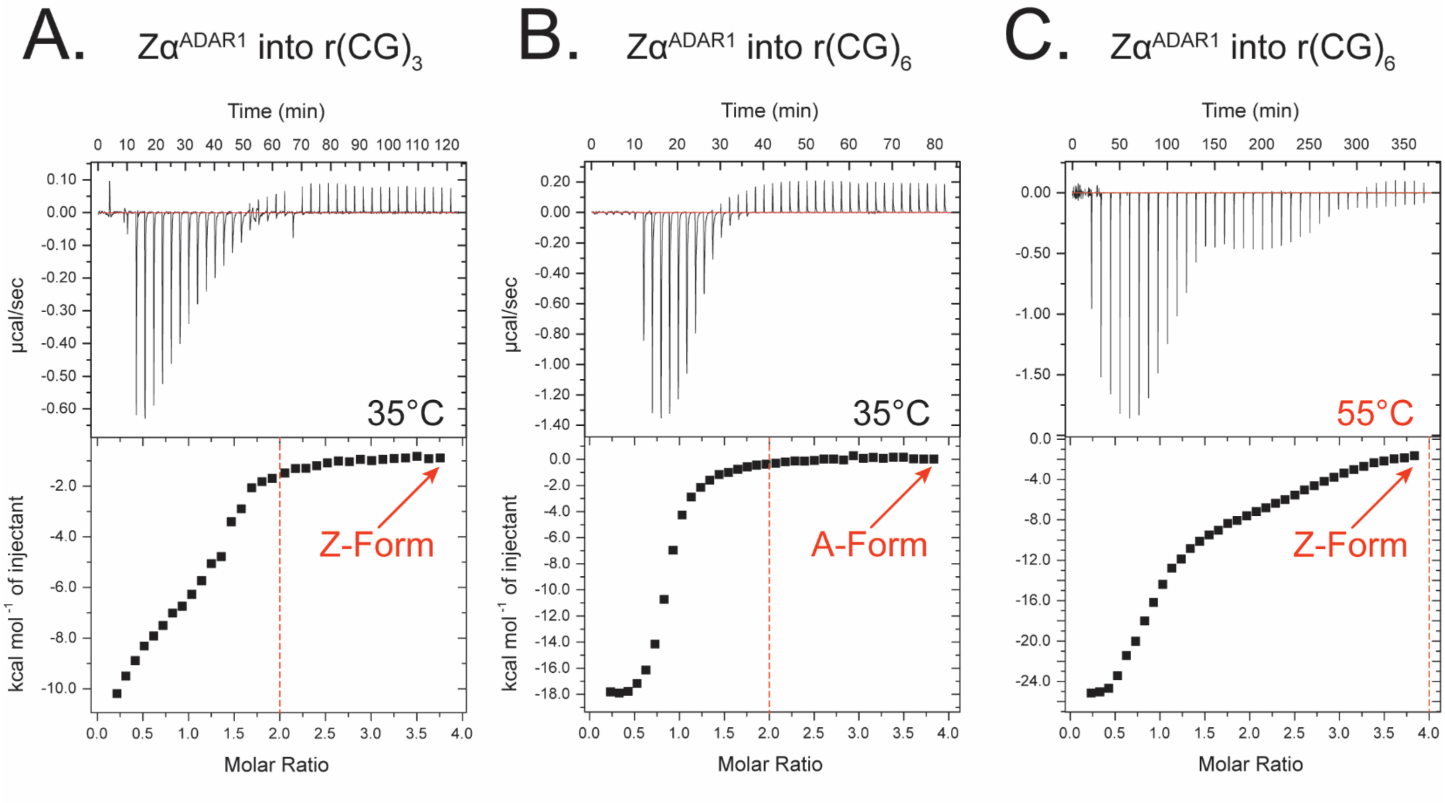


**Supplementary Figure 13.** Raw Isothermal Titration Calorimetry measurements comparing differences in stoichiometry between a 6-mer r(CpG)$_3$ oligo at 35°C and 2 min injection intervals **(A)**, a 12-mer r(CpG)$_6$ oligo at 35°C and 2 min injection intervals **(B)**, and the same 12-mer r(CpG)$_6$ oligo at 55°C and 10 min injection intervals **(C)**.
